# Emergent coexistence and the limits of reductionism in ecological communities

**DOI:** 10.1101/2025.05.15.654235

**Authors:** Guim Aguadé-Gorgorió, Sonia Kéfi

**Affiliations:** ISEM, University of Montpellier, CNRS, IRD, EPHE, Montpellier, France; Santa Fe Institute, 1399 Hyde Park Road, Santa Fe, NM 87501, USA

## Abstract

Understanding if pairwise interactions explain the species composition of communities is a central goal in ecology. This question has been challenged by the observation of *emergent coexistence*, where microbial communities contain species that cannot coexist in pairs, suggesting the presence of non-pairwise mechanisms. Instead, we show that emergent coexistence arises naturally in species-rich models with pairwise interactions. Strikingly, this phenomenon does not require additional mechanisms like intransitive or higher-order interactions; rather, coexistence arises from dense networks of indirect effects. As diversity increases, we show that indirect effects become so intricate that pairwise interactions no longer predict community composition, revealing a fundamental limit to reductionist explanations of coexistence. Like chaos emerging from simple rules, our findings provide theoretical foundations to understand how unexpected species coexistence can emerge from pairwise interactions.

---

*In this great chain of causes and effects, no single fact can be considered in isolation*.

- Alexander Von Humboldt

Ecological systems are inherently complex, because of the number of constituent species and the diverse ways in which they depend on each other. Typically, ecological communities are conceptualized as a set of species connected by a network of pairwise interactions [1]. Despite knowing that additional mechanisms other than pairwise interactions may be in place [2], this framework continues to dominate how ecological communities are conceptualized and studied. A central question in ecology is whether pairwise interactions alone are sufficient to understand and predict community properties such as coexistence and stability.

Microbial ecosystems offer a powerful experimental platform for investigating how species interactions govern fundamental community properties [3; 4; 5; 6; 7]. Their unique tractability enables researchers to isolate individual species and perform controlled co-culture experiments, which allows inferring pairwise ecological relationships [8; 9; 10] (Fig. 1). This capability naturally leads to a fundamental question: Can knowledge of pairwise interactions reliably predict which combinations of these species will form a stable community, especially so as diversity increases (Fig. 1)? Specifically, does pairwise coexistence translate to coexistence in larger communities, while species pairs exhibiting competitive exclusion are inherently incompatible? A confirmation of these patterns would support reductionist views of community assembly, while observed deviations could indicate the presence of other complex ecological phenomena that transcend pairwise relationships.

**FIG. 1.**
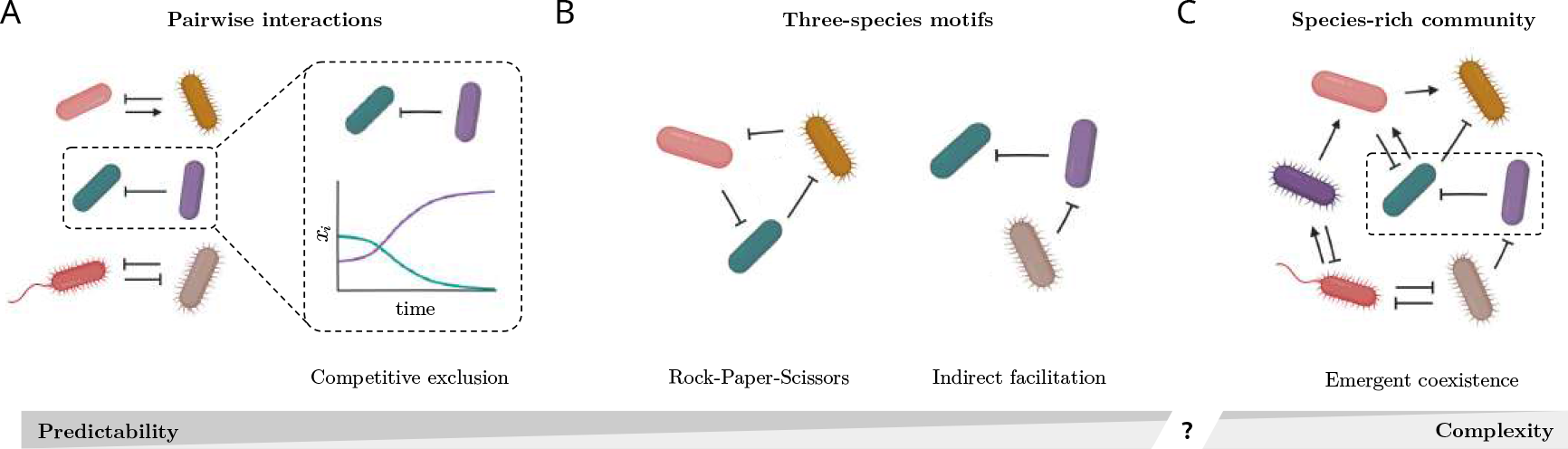
Can we predict community composition from pairwise interactions? (A) Experimental studies of microbial communities are able to grow species by pairs and infer if they coexist or exclude one another (dashed box, [9]). (B) In three-species systems, competitive exclusion can be avoided with rock-paper-scissors loops or indirect facilitation [11]. (C) Yet, species-rich microbial communities can harbor many excluding pairs and no rock-paper-scissors loops, hinting to the possibility that coexistence is an emergent, community-level property [9].

A significant step forward has been the experimental observation that stable microbial communities often contain many species pairs that, when isolated, fail to coexist due to strong competition [8; 9; 10; 12] (Fig. 1A). This phenomenon, in which community composition seems to contradict expectations from pairwise interactions, has recently been labeled as *Emergent Coexistence* (EC, [9], Fig. 1C). While cooperative interactions are common in microbial species [13], the apparent incompatibility of many of their constituent species raises questions about the mechanisms that support stable coexistence in ecological communities. This apparent paradox suggests that pairwise interactions alone may be insufficient to predict species coexistence, pointing towards more complex, community-level mechanisms.

Mathematical models have long been a cornerstone in exploring how species interactions shape their coexistence [14; 15]. Many mechanisms have been proposed to circumvent the problem of competitive exclusion, including positive interactions [16], non-random network structures [17] or the role of space in reducing effective competition [15]. In laboratory observations of EC, however, pairwise competition can be abundant and space does not seem to play a major role [8; 9; 10; 12]. In this context, typical explanations involve higher-order interactions or intransitive competition [1; 18]. Higher-order interactions emerge when a third species can modulate the competitive strength between the first two species [2; 18]. Intransitive competition is characterized by a lack of competitive hierarchy, for example in a rock-paper-scissors loop (Fig. 1B, [19]). Both of these mechanisms could buffer pairwise exclusions through the presence of a third species. However, their empirical prevalence and explanatory power in natural and laboratory ecosystems remain unclear, and EC has been observed without intransitive loops [1; 9; 12; 20; 21].

In the absence of higher-order or intransitive mechanisms, a species-rich community is still pervaded by indirect effects, by which a species’ growth is influenced by chains of cumulative effects through other species (Fig. 1C, [11; 22; 23]). These indirect effects, exemplified by the principle “the enemy of my enemy is my friend”, have been extensively studied in small-scale interaction motifs [11; 22; 24] (Fig. 1B). However, as community complexity increases, quantifying the magnitude and significance of these indirect effects in both theoretical models and empirical systems remains a formidable challenge [23; 24; 25; 26; 27] (Fig. 1C).

Here, we study if EC is possible in species-rich models of random pairwise interactions due to indirect effects alone, or else if additional mechanisms need to be invoked. In the main text we explore EC in the generalized Lotka-Volterra (GLV) model, a simple phenomenological description of a community of many interacting species [6; 7; 10; 28], and analyse other dynamical models in ESM I.N. Our main goal is to understand how pairwise interactions shape –and allow us to predict– the species composition of a community. To focus on the role of interactions, a common assumption is that growth rates are uniform across species (*r*_*i*_ = *r*), and that interaction strengths are rescaled relative to self-regulation (*A*_*ij*_ = *a*_*ij*_*K*_*i*_; [27; 30], ESM I.A). In this setting, the abundance dynamics of species *i* in the GLV model follows:

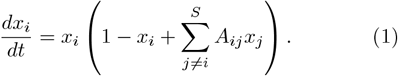

The growth of species *i* is affected by linear replication, quadratic self-regulation and the interaction with each of the other species, mediated by *A*_*ij*_. However, estimating all the *S*(*S* − 1) interactions for a species-rich system is an utterly complex task [10; 32; 33]. Instead, a common procedure is to assume that we can only infer the statistical properties of *A* such as the interaction strength (*µ*), heterogeneity (*σ*) and network connectivity (*C*) (ESM I.C-D; [28; 34; 35]. While interaction networks can be highly structured in macro-organisms [17], the random interaction assumption has been successful at characterizing microbial community properties [7; 30; 36; 37] (ESM I.C-D). Starting from a large pool of *S* randomly interacting species, characterized by an interaction matrix *A*, are there subsets of species that can coexist stably even if some of their constituent species do not coexist in pairs (Fig. 1)? Here, we study whether this is possible, the mechanisms behind it, and its consequences for the predictability of community composition.

## RESULTS

### Emergent coexistence is common in models with random pairwise interactions

We explore the presence of stable communities with EC in the GLV model (Eq. 1). We start from a pool of *S* species and random interaction coefficients *A*_*ij*_ defined by *µ* and *σ*, and study if the system reaches a stable state where at least one pair of the coexisting species would not coexist outside of the community due to competitive exclusion (*A*_*ij*_ *<* −1, ESM I.B,I.E). Our results reveal that EC is a common outcome in the GLV model, provided that pairwise interactions are strong and heterogeneous. More specifically, we identify a large domain of the phase space, inside of which a large fraction of stable communities harbor species pairs that cannot coexist in isolation (Fig. 2A,B), and we provide analytical estimates for its boundaries (ESM II.A).

**FIG. 2.**
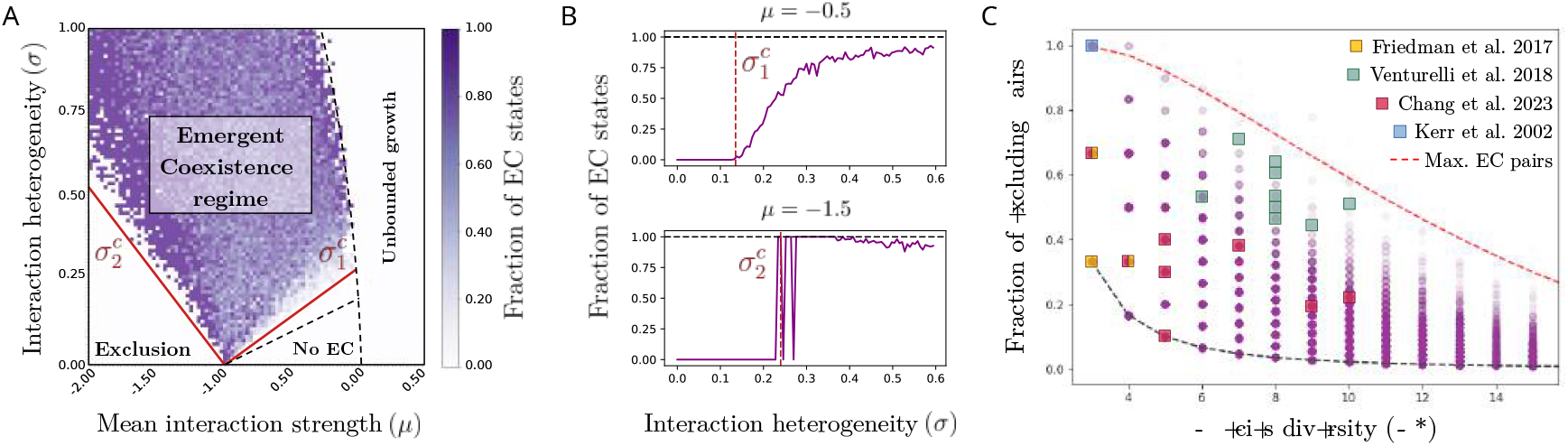
Emergent coexistence in models of pairwise interactions. (A) Simulations of the GLV model starting from a pool of *S* = 80 species and interactions randomly sampled from a normal distribution (ESM I.E). The GLV model has a rich phase space with four dynamical regimes depending on the values of *µ* and *σ* [28; 29; 30]. Most stable communities harbor EC in an area delimited by two analytically estimated boundaries (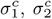, red lines, ESM II.A). This EC regime is found inside the multistability regime, one of the four dynamical regimes of the GLV model (delimited by dashed lines, [31]). (B) Vertical slices of (A) at *µ* = −0.5 and *µ* = −1.5 showing these transitions towards the regime where EC is prevalent. (C) Diversity and fraction of excluding pairs in communities with EC found in the simulations of (A). Simulated communities (purple circles) and empirical data [8; 9; 10; 19] (colored squares) are consistent with the predicted maximum fraction of excluding pairs that stable communities can sustain (red and dark dashed lines, ESM II.A).

Notably, EC occurs inside a regime of the GLV model characterized by multiple stable states [28; 31] (Fig. 2A). This is consistent with empirical observations that have linked EC to the presence of alternative community states depending on initial conditions [9; 38]. We also find that EC is a prevalent outcome beyond the assumption of fully random interactions, when *A*_*ij*_ have predator-prey structure [39], are highly skewed or sparse [33; 36; 40], or have nested or single-resource structures (Fig. S2, ESM II.A). Finally, we show that stable communities with EC are also present in models with multilayer interactions, saturating responses or sublinear growth (ESM I.N,II.A), while very recent work has found similar outcomes in models with growth-competition and competition-colonization trade-offs [41]. Taken together, these results suggest that EC is not particular to the GLV model or the random interaction assumption, but rather is a general outcome in species-rich models once many interactions are competitive and heterogeneous.

### Analytical predictions match observations of the fraction of excluding pairs

A striking observation of EC experiments is that as many as 60 ∼ 70% of species pairs in a stable community do not coexist in co-culture [9; 10]. For each community with EC that we find, we measure the fraction of species pairs that interact via competitive exclusion and hence would not coexist if isolated (Fig. 2C). We find that the maximum fraction of excluding pairs in a community decays with species diversity, and that this is related to a theoretical limit on competition strength and heterogeneity beyond which a species-rich system becomes unstable (Fig. 2C, ESM II.A) [28; 34]. Using recent corrections of this limit for communities of moderate size [31], we provide an analytical estimate for the maximum fraction of excluding pairs as a function of species diversity (Fig. 2C, ESM II.A). Our result is consistent with experimental data [8; 9; 10] (ESM I.G), and confirms that the fraction of excluding pairs can be very large in communities of moderate size: a community of 10 species could sustain as many as 60% of species pairs that do not coexist in co-culture. A natural question is to ask how all these species can coexist if so many interactions are strongly competitive and there are no higher-order effects.

### Indirect effects limit the predictability of species coexistence

Early work on three-species models showed that positive indirect effects of length two, as in “the enemy of my enemy is my friend”, could allow competitors to coexist ([11; 24], ESM I.B). Yet, in networks of many interacting species and long chains of indirect effects, it is unclear what determines who is friend and who is enemy. Importantly, recent work has shed new light to our understanding of indirect effects in species-rich communities [27].

Given a pool of *S* species interacting through matrix *A*, equilibrium states of the GLV model will be those in which at least a subset of *S*^∗^ species with interactions encoded in *A*^∗^ coexist with positive abundances. This translates to finding a subset of species with positive abundances by solving (ESM I.B):

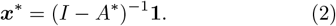

Here 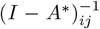 encodes the net effect of species *j* on the equilibrium abundance of species *i* [24; 25; 26; 27] (ESM I.B). The link between direct and net effects can be better understood by using the Neumann series [27]:

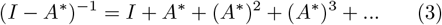

so that the equilibrium abundance of a given species becomes

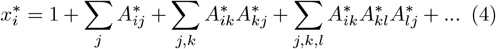

The abundance of a species is affected by direct interactions, but also by the cumulative effects of indirect interactions through all other species [27]. A fundamental consequence of this is that, even when direct interactions are all competitive, we find that half of the net effects between species in a community can be positive (Fig. S5, ESM II.B). The reductionist expectation is that if we can estimate *A*_*ij*_ experimentally, we can assemble a community by selecting species pairs that can coexist [8]. Yet, if most species are competitors, but half of the net effects are positive, how can we determine who is a friend of whom in the context of the full community?

The validity of the reductionist approach is determined by the *collectivity* metric *ϕ*, the spectral radius of the interaction matrix *A*^∗^ ([27], ESM II.B). In strongly interacting and highly connected communities, chains of indirect effects become stronger as they increase in length, until the series in equations (3),(4) does not converge. This happens when *ϕ* is larger than 1 [42]. Even when *ϕ* is below 1, but close to it, an accurate prediction of species abundances would require estimating many powers of *A*^∗^ (Eq. 4, ESM II.B). We find that most communities with EC found in the previous section have *ϕ* close to or larger than 1, with the smallest possible *ϕ* increasing as *ϕ* ∼ *S*^∗^|*µ*^∗^| (Fig. 3A, ESM II.B, [27; 39]). An explanatory consequence of high collectivity is that direct and net interactions become uncorrelated: species *j* can compete strongly with species *i*, but, through indirect effects across the community, exert a positive net effect on it (Figs. 3B, S6). Collectivity therefore provides the mathematical basis for why EC is not predictable from co-culture observations.

**FIG. 3.**
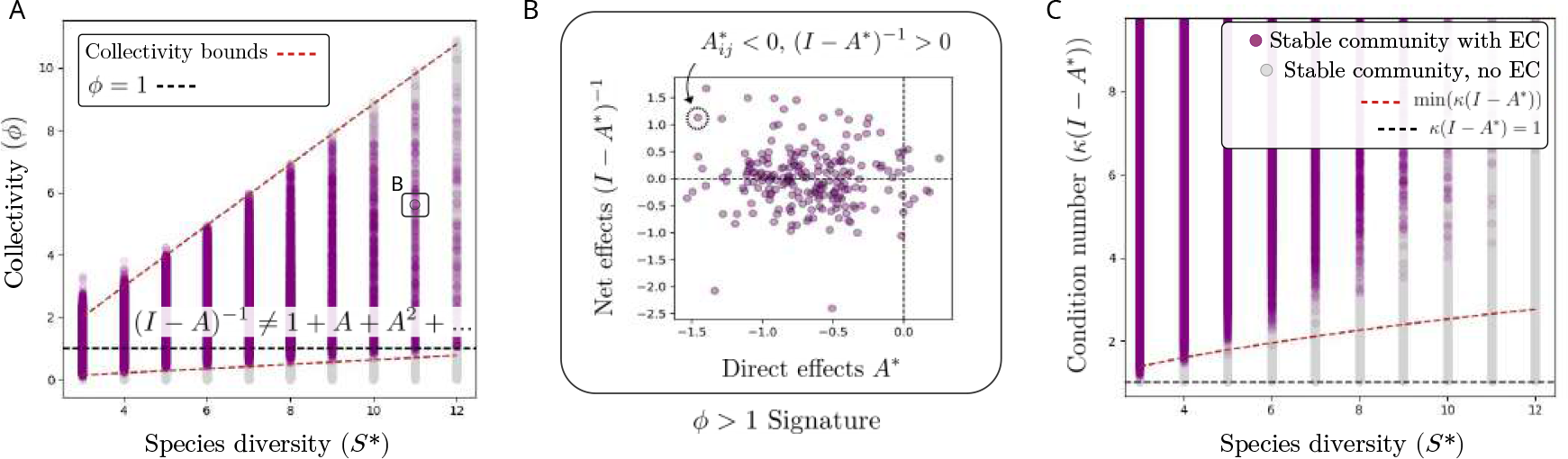
Indirect effects limit the predictability of community composition. Collectivity (A) and Condition number (C) for stable communities with (purple) and without (gray) EC. (A) Most communities with EC have collectivity (*ϕ*) close to or larger than 1 (dark dashed line), with analytical estimates predicting *ϕ* ∼ *S*^∗^|*µ*^∗^| (red dashed lines, ESM II.B). (B) For *ϕ >* 1, direct interactions do not correlate with net species effects: a species can be a strong competitor, but still provide a positive net effect through interactions with the rest of the community (dashed circle). (C) Most communities with EC also have a condition number larger than 1, with the lowest expectation increasing as *κ* ∼ *S*^∗^ (ESM II.B).

If net effects cannot be directly inferred from pairwise interactions, an alternative approach could be to estimate them by calculating (*I* − *A*^∗^)^−1^ numerically (Eq. 2). The feasibility of this method can be assessed using the *condition number κ*(*I* − *A*^∗^), which quantifies how small measurement errors in *A*^∗^ are amplified when inverting the matrix, with *κ* = 1 being the best-case scenario (ESM II.B, [43; 44; 45]). Our analysis reveals that for communities exhibiting EC, the lowest *κ* increases with species diversity as *κ* ∼ *S*^∗^, while most communities display significantly higher *κ* values (Figs. 3C, S7). This means that small errors in the measurements of pairwise interactions (*A*^∗^) can lead to substantial errors in the inverse matrix and therefore a decreased predictability of species abundances at equilibrium (Fig. 3C, ESM II.B).

These results align with evidence that reductionist assembly rules work well for small communities of three or four species [8], but can break down in more diverse communities where indirect effects become pervasive [9]. The simplicity of estimating *ϕ* and *κ* also suggests that these parameters offer valuable metrics for assessing ecological predictability, as demonstrated in two experimental case studies in ESM II.B.

### Balanced feedback loops maintain stability under strong competition

The results above describe the mechanisms by which strongly competing species can coexist due to indirect effects. Beyond coexistence, experimental communities with EC have also been found to be dynamically stable [8; 9], and we study here the mechanisms that guarantee this stability. Mathematically, linear stability can be captured by the eigenvalues of the Jacobian matrix at equilibrium having negative real part (ESM II.C) [39].

Theoretical results propose that a community will have positive eigenvalues (and hence, become unstable) once diversity, interaction strength and heterogeneity overcome a predictable threshold [28; 34; 39]. Interestingly, we find that communities with EC can remain stable even if competition is above this threshold (Fig. S9, ESM II.C, [28]). This is because random matrix predictions only operate for very large communities (ESM II.C, [31]). For smaller communities like those found in EC experiments (Fig. 2C), not only the aggregate statistics of interactions, but also their species-level organization can play a role in their dynamics (ESM II.C). We find that communities with moderate diversity and strong interactions can be linearly stable provided that they fulfill the Routh-Hurwitz criteria (Fig. S9). These criteria require that interactions are structured so that negative, stabilizing loops control positive loops that would instead amplify perturbations (ESM I.K, [23; 46]). Our results show that community stability is possible under strong competition and high collectivity, provided that specific loop architectures are maintained (ESM II.C, [42]).

### Emergent coexistence does not require intransitive competition

Coexistence under strong competition has often been linked to intransitivity in theoretical models [1; 18; 47; 48]. Intransitivity can be defined by a lack of hierarchy in species competition, by which no species can become the most dominant of the community. This can be assessed by the presence of rock-paper-scissors loops or exclusions by species with a lower rank in the competitive hierarchy, among many possible metrics (ESM I.L). Yet, in empirical communities with more than three species, the presence and role of these motifs remain unclear. While some communities appear highly intransitive [49], evidence indicates that intransitivity is in fact rare across species-rich systems [1; 20; 21]. Experimental communities with EC have been observed with strikingly low intransitivity percentages (0 − 0.3% of triplets are RPS, 1 − 3% of exclusions are LRE in [9]-[12]).

Here, we find that intransitivity patterns in communities with EC become rarer as species diversity increases (Figs. 4, S12, S13). In small communities, specific intransitive motifs are common and necessary for coexistence (Fig. 2C). In more complex and collective communities, many interaction chains can rescue species from extinction, and the presence of intransitive motifs simply approaches the random expectation (Fig. 4, ESM II.D). However, rock-paper-scissors motifs are almost absent from diverse EC communities assembled in the lab [9; 12], which contradicts the random expectation of ∼ 25% of triplets being rock-paper-scissors (Fig. 4C,D, ESM II.D). One possible explanation could be a fully nested interaction matrix *N*^∗^, where all interactions are located in the upper triangle of the matrix (Fig. 4C, ESM I.M). Fully hierarchical competition matrices, although unrealistic, can generate communities with EC and no rock-paper-scissors, proving that intransitive loops are not necessary for EC (ESM II.A, [41]).

**FIG. 4.**
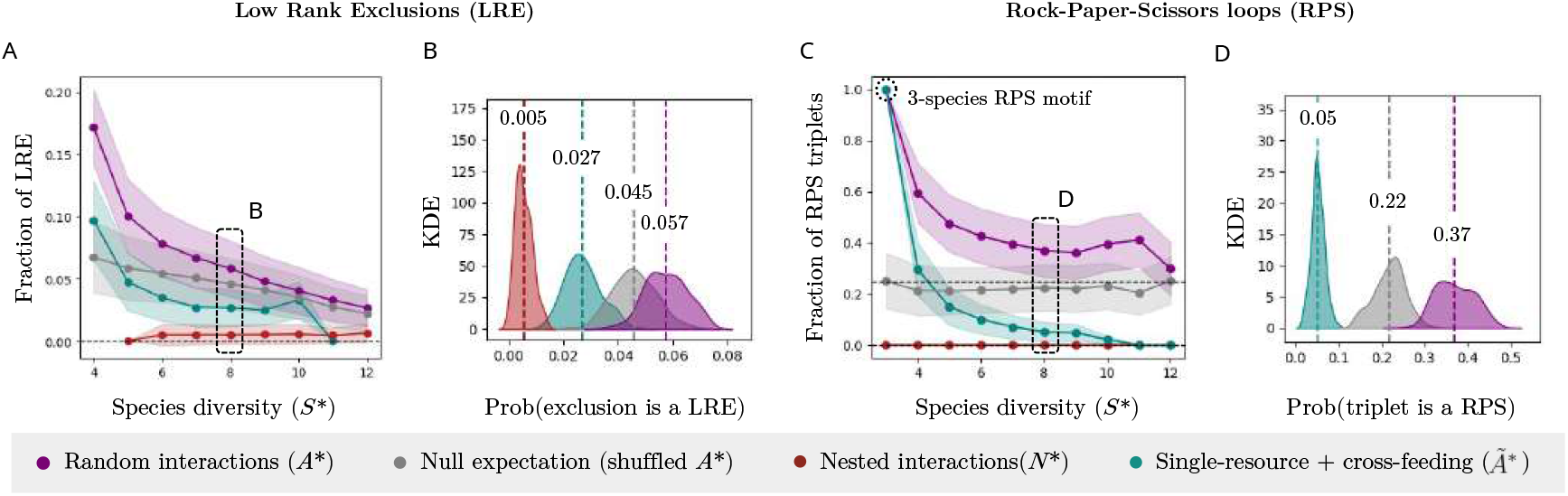
Emergent coexistence does not require intransitive competition. (A,B) Fraction of exclusions where a lower-ranked species excludes a higher-ranked one (LRE), where the rank is defined as the number of wins minus number of losses divided by the number of interactions (ESM I.L). (C,D) Fraction of triplets that are rock-paper-scissors (RPS), where a triplet is a trio of species all connected by competitive exclusion interactions (ESM I.L). We compare intransitivity in communities with EC found under random interactions (*A*^∗^, purple), reshuffled *A*^∗^ (gray), fully nested interactions (*N*^∗^, red) and single-resource competition with cross-feeding (*Ã*^∗^, green). (B,D) Kernel Density Estimates (KDE, ESM I.L) allow us to visualize the expected probability distribution of intransitivity metrics in communities with *S*^∗^ = 8 species.

Observations of transitive EC could also be linked to specific experimental conditions, in which species compete for a single resource and cooperate via cross-feeding [9; 38; 50]. In the scenario where competition is defined by a single trait (the capacity to consume the available resource, *γ*_*i*_), competition becomes strictly hierarchical [51]. On top of this, we can incorporate additional weak competition mechanisms *a*_*ij*_ [52], together with positive cross-feeding *C*_*ij*_ *>* 0 [38; 50]. The modified competition matrix *Ã* writes (ESM I.M)

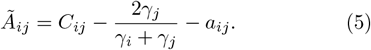

When two species compete, the species with the lowest *γ*_*i*_ perceives stronger competition, resulting in a hierarchical structure where the most efficient consumer dominates (ESM I.M, [14]). We find that *Ã* can generate communities with EC and very low intransitivity values consistent with laboratory observations of EC (Fig. 4, teal, ESM II.A,II.D).

## DISCUSSION

Species-rich microbial communities assembled in laboratory conditions frequently harbor species that fail to coexist in isolated pairs due to strong competition [8; 9; 10]. This observation of emergent coexistence (EC) challenges the reductionist approach of predicting community composition solely from pairwise interactions and suggests the potential importance of non-pairwise, higher-order mechanisms [1]. Our research, however, demonstrates that EC is a common outcome in species-rich models of pairwise interactions, with our analytical and numerical predictions aligning with experimental observations. We have showed how indirect effects can maintain species coexistence, but become increasingly complex as species diversity increases. Beyond a predictable threshold in interaction strength and species diversity, direct interactions alone can no longer predict community coexistence. This finding provides a mathematical foundation for EC and imposes an explicit limit to the reductionist approach in community assembly. These results further emphasize the need for coarse-grained models that can capture global community properties without relying on precise species-level information.

Our work confirms that EC is intrinsically an emergent phenomenon [53]: in the presence of strong indirect effects, community composition is driven by, but cannot be analytically predicted from, pairwise interactions between species. This places EC in a similar conceptual category as deterministic chaos, where simple rules can produce systems with complex and unpredictable dynamics [54]. Our results also align with a broader challenge across complex systems science: understanding the limits of our ability to predict a system’s future configuration based on its current state [55]. In ecology, this challenge is often framed in terms of forecasting temporal responses to perturbations [45; 56; 57]. Our work extends this perspective by identifying computable limits to the predictability of stable community composition. These insights have practical implications for the rational design and bottom-up assembly of microbial consortia, with direct applications in fields such as biomedicine [58] and environmental restoration [59].

Ecological communities are undoubtedly shaped by intricate processes beyond random pairwise interactions, including consumer-resource dynamics [50], higher-order effects [2] and non-random interactions [17]. Our work does not claim that these mechanisms are not relevant for ecological dynamics. Rather, we demonstrate that, even in their absence and assuming the simplest possible pairwise interaction models, ecological complexity can already give rise to unexpected collective properties. By examining empirical observations of EC, our research highlights the crucial role of indirect effects in molding emergent community behavior.

## METHODS

All mathematical and numerical methods are discussed in detail in the Electronic Supplementary Material. All codes for simulations are found at https://github.com/GuimAguade/EmergentCoexistence.

### Rescaling the GLV model

We study the GLV model of *S* interacting species [6; 28; 31], with species labeled as *i* = 1, 2, …*S* and abundance dynamics following

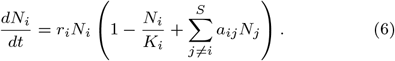

To focus on the role played by interspecies interactions, a typical procedure (see e.g. [27; 31]) is to assume homogeneous growth rates *r*_*i*_ = *r*, rescale time as *t* → *rt*, divide species abundances by carrying capacities (*x*_*i*_ ≡ *N*_*i*_*/K*_*i*_) and rescale interaction strengths relative to self regulation (*A*_*ij*_ ≡ *a*_*ij*_ */a*_*jj*_ = *a*_*ij*_ *K*_*j*_). This allows us to reach equation (1) where *A*_*ij*_ are the only controling parameters of the dynamics. We present other dynamical models in I.N.

### Simulating GLV dynamics

Simulations start by defining a random matrix of *S* = 80 interacting species. *A*_*ij*_ are sampled from a Gaussian distribution with *µ* ∈ (−2, 0.5) and *σ* ∈ (0, 1) (Fig. 2A-C), while other interaction structures are described in ESM I.M. We set random initial conditions *x*_*i*_(*t* = 0) ∈ *U* [0, 1], integrate the system for ∆*t*_1_ = 3 *·* 10^3^ timesteps and check if abundances are the same after ∆*t*_2_ = 10^2^ additional timesteps to test for stationarity [29; 31]. Once a final state is reached, we identify those *S*^∗^ ≤ *S* species that have survived with positive abundance and capture their interactions in *A*^∗^. A state has EC if *S*^∗^ ≥ 3, and there is at least one exclusionary interaction 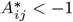. The fraction of excluding pairs (Fig. 2C) is the number of pairs for which 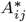 or 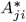 are smaller than −1, divided by the total number of pairs of surviving species, *S*^∗^(*S*^∗^ − 1)*/*2.

### Sampling stable subsets of *A*^∗^

We use a complementary analytical technique to infer stable states from Eq. (1) more efficiently [31] (Figs. 3A-C,4A-D). We define *A* as above, and define a random subset of *S*^∗^ ≤ *S* species and their interactions *A*^∗^. We measure the equilibrium abundances by solving Eq. (3) numerically, and check if all abundances are positive, and eigenvalues of the Jacobian matrix 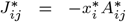 are negative (ESM I.E.2). The subsets of *A*^∗^ that fulfill these two conditions are stable solutions of equation (1) in the absence of migration [31], and we use the same test as above to evaluate EC.

### The boundaries of the EC regime

To estimate the boundaries of the EC regime in the (*µ, σ*) phase space (Fig. 2A,B red lines), we study two *necessary* conditions (ESM II.A). First, we search for the minimal (*µ, σ*) combination for which at least one element within *S*(*S* − 1) elements of *A* will be exclusionary (*A*_*ij*_ *<* −1, ESM II.A.1). The probability of *A*_*ij*_ *<* −1 is *P* = 1 − (1 − Φ (−(1 + *µ*)*/σ*))^*S*(*S*−1)^, where Φ(*z*) is the cumulative distribution function (CDF) of the standard normal distribution. If *S* is large, this function predicts a sharp transition from *P* = 0 to *P* = 1 for given (*µ, σ*) values (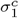 in Fig.2). Second, using a similar approach, we search for the maximal (*µ, σ*) combination for which at least one species pair is not exclusionary (*A*_*ij*_,*A*_*ji*_ *>* −1) and can rescue a third excluded species (ESM II.A.2).

### The maximum fraction of excluding pairs

To compute the fraction of excluding pairs as a function of *S*^∗^ (Fig. 2C), we use recent analytical results to compute the largest species richness *S*^∗^ given (*µ, σ*) (ESM II.A.3, [31]). We also use the method explained above to estimate the fraction of 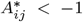 elements given (*µ, σ*). The maximum fraction of *A*^∗^ *<* −1 elements is, at most, the fraction of *A*_*ij*_ *<* −1 elements in the pool. Taken together, for each (*µ, σ*), we can predict both the largest possible community (*S*^∗^) and the maximum fraction of excluding pairs it can contain (ESM II.A.3).

### Analytical bounds on collectivity *ϕ*

In Fig. 3A we plot the spectral radius *ϕ* of *A*^∗^ of stable communities with (purple) and without (gray) EC, found from numerically solving the GLV model with (*µ, σ*) in the range of Fig. 2A and measuring the modulus of the largest eigenvalue of *A*^∗^ with `numpy.linalg.eigvals`. The analytical maximum and minimum *ϕ*(*S*^∗^), we use a result from random matrix theory [27; 39] (ESM II.B.3). Again, we use the methods from [31] to find the largest and smallest *S*^∗^, |*µ*^∗^|, *σ*^∗^ values given each original (*µ, σ*), which define the largest and smallest *ϕ* values for communities with EC (ESM II.B.3).

### Analytical bounds on condition number *k*

In Fig. 3C we plot the condition number of the interaction matrix *A*^∗^ of stable communities with (purple) and without (gray) EC, found from numerically solving the GLV model with (*µ, σ*) in the range of Fig. 2A. The condition number follows *k*(*A*^∗^) = *s*_*M*_ (*A*^∗^)*/s*_*m*_(*A*^∗^), where *s*_*i*_ are the largest (*M*) and smallest (*m*) singular values of the matrix *A*^∗^ (ESM I.H). The smallest possible *k*(*A*^∗^) in a set of systems with different *A*^∗^ happens for the smallest *s*_*M*_ and largest *s*_*m*_. RMT provides estimates for these two values, allowing us to estimate the minimal condition number of a set of states with EC (ESM II.B.4). [43].

### Stability and feedback loops

A state is linearly stable if it recovers from infinitessimal perturbations, *i*.*e*. all eigenvalues of the Jacobian matrix evaluated at that equilibrium point are negative [39]. A state with random interactions becomes unstable once *S, µ* or *σ* overcome a predictable threshold [28; 34], yet we find that this threshold does not apply for communities with a handful of species and strong interactions (ESM II.C). The Routh-Hurwitz criteria link linear stability to conditions on the coefficients *C*_*i*_ of the characteristic polynomial of the Jacobian matrix, found by solving det(*J* − *λI*) = 0 with 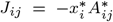 (ESM I.K, [23; 46]). We find *C*_*i*_ numerically with numpy.poly and measure the second Routh-Hurwitz stability condition that requires Λ_2_ = *C*_1_*C*_2_ − *C*_3_ *>* 0 (Fig. ESM 10).

### Intransitive competition motifs

A triplet is a set of three species connected by competitive exclusion 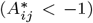. A rock-paper-scissors triplet has a non-dominant architecture (A excludes B, B excludes C, C excludes A, ESM I.L). The rank of a species is measured as the number of wins minus the number of loses divided by the total number of interactions, following e.g. [9; 12]. A low rank exclusion happens when a species with lower rank excludes a species with higher rank, which requires at least 4 exclusions (ESM II.D.1). To visualize the probability distribution of rock-paper-scissors and low rank exclusion fractions (Fig. 4), we find states with EC as described above and select subsets of 100 communities with EC, measure the distribution of observed intransitivity values across them and plot kernel density estimates (KDE) for these distributions (ESM I.L). This sampling is chosen to replicate an experimental setting where intransitivity is measured across a set of different communities [9; 12]. We find similar results with other intransitivity metrics in ESM I.D.3.

### Nested and single-nutrient competition matrices

The matrix *N*^∗^ is generated from *A*, but imposing that all elements in the lower triangle of *A* are zero (*A*_*ij*_ = 0 if *i > j*). For *Ã*, positive *γ* values are sampled from *U* [0.3, 0.7] and sorted by decreasing order to build a hierarchical matrix; *a*_*ij*_ are sampled from *N* (0.1, 0.01); *C*_*ij*_ are sampled from *U* [0.0, 1.0] (ESM I.M). We do not have empirical support for these parameters and use them only as a proof of concept to explain how hierarchical competition and random cross-feeding can lead to transitive communities with EC (ESM II.A, II.D). If the strength or heterogeneity of *a*_*ij*_ increases, the model becomes similar to the random interactions model. If the strength or heterogeneity of *C*_*ij*_ increases, the model transitions towards the outgrowth regime of fig. 2A (ESM II.A).

## ACKNOWLEDGEMENTS

The initial inspiration for this work comes from a talk given by Hyunseok Lee in Paris, in a seminar hosted by Daniel R. Amor in June 2024. We are most grateful to H. Lee for the presentation and the insightful discussions that followed. We also thank specially J.-F. Arnoldi, I. Lajaaiti and B. Pichon with whom we have discussed many of these ideas throughout the years, R. Solé, M. Barbier, J. Piñero, V. Maull, L. Arola-Fernandez, M. Sireci and J. Leigh for their support, and the hospitality of the Santa Fe Institute where much of this work was done. Special thanks to F. Browne and S. Kea for inspiration. Drawings in Figure 1 are created with BioRender.com. G.A-G. was supported by a 2022 postdoctoral fellowship of the Fundación Ramón Areces and a Marie Skłodowska-Curie Actions Postdoctoral Fellowship under project FRAGILEPRINTS - 101105029. Views and opinions expressed are however those of the author(s) only and do not necessarily reflect those of the European Union or the CNRS. Neither the European Union nor the CNRS can be held responsible for them.

## REFERENCES

[1] J. M. Levine, J. Bascompte, P. B. Adler, and S. Allesina, Nature 546, 56 (2017).

[2] I. Billick and T. J. Case, Ecology 75, 1529 (1994).

[3] K. Z. Coyte, C. Rao, S. Rakoff-Nahoum, and K. R. Foster, PLoS biology 19, e3001116 (2021).

[4] S. Arya, A. B. George, and J. O’Dwyer, Current Opinion in Microbiology 83, 102580 (2025).

[5] S. Widder, R. J. Allen, T. Pfeiffer, T. P. Curtis, C. Wiuf, W. T. Sloan, O. X. Cordero, S. P. Brown, B. Momeni, W. Shou, et al., The ISME journal 10, 2557 (2016).

[6] N. I. van den Berg, D. Machado, S. Santos, I. Rocha, J. Chacón, W. Harcombe, S. Mitri, and K. R. Patil, Nature ecology & evolution 6, 855 (2022).

[7] J. Hu, D. R. Amor, M. Barbier, G. Bunin, and J. Gore, Science 378, 85 (2022).

[8] J. Friedman, L. M. Higgins, and J. Gore, Nature ecology & evolution 1, 0109 (2017).

[9] C.-Y. Chang, D. Bajić, J. C. Vila, S. Estrela, and A. Sanchez, Science 381, 343 (2023).

[10] O. S. Venturelli, A. V. Carr, G. Fisher, R. H. Hsu, R. Lau, B. P. Bowen, S. Hromada, T. Northen, and A. P. Arkin, Molecular systems biology 14, e8157 (2018).

[11] J. M. Levine, Ecology 80, 1762 (1999).

[12] L. M. Higgins, J. Friedman, H. Shen, and J. Gore, BioRxiv, 175737 (2017).

[13] J. Kehe, A. Ortiz, A. Kulesa, J. Gore, P. C. Blainey, and J. Friedman, Science advances 7, eabi7159 (2021).

[14] D. Tilman, Resource competition and community structure, 17 (Princeton university press, 1982).

[15] P. Chesson, Annual review of Ecology and Systematics 31, 343 (2000).

[16] S. P. Hart, Journal of Ecology 111, 2094 (2023).

[17] P. C. De Ruiter, A.-M. Neutel, and J. C. Moore, Science 269, 1257 (1995).

[18] L. Gallien, N. E. Zimmermann, J. M. Levine, and P. B. Adler, Ecology Letters 20, 791 (2017).

[19] B. Kerr, M. A. Riley, M. W. Feldman, and B. J. Bohannan, Nature 418, 171 (2002).

[20] O. Godoy, D. B. Stouffer, N. J. Kraft, and J. M. Levine, “Intransitivity is infrequent and fails to promote annual plant coexistence without pairwise niche differences,” (2017).

[21] F. Koch, A.-M. Neutel, D. K. Barnes, K. Tielborger, C. Zarfl, and K. T. Allhoff, Communications Biology 6, 690 (2023).

[22] J. T. Wootton, Annual review of ecology and systematics, 443 (1994).

[23] R. Levins, Annals of the New York Academy of Sciences 231, 123 (1974).

[24] S. H. Levine, The American Naturalist 110, 903 (1976).

[25] J. M. Dambacher, H. W. Li, and P. A. Rossignol, Ecology 83, 1372 (2002).

[26] J. Montoya, G. Woodward, M. C. Emmerson, and R.V. Solé, Ecology 90, 2426 (2009).

[27] Y. R. Zelnik, N. Galiana, M. Barbier, M. Loreau, E. Galbraith, and J.-F. Arnoldi, Ecology Letters 27, e14358 (2024).

[28] G. Bunin, Physical Review E 95, 042414 (2017).

[29] G. Aguadé-Gorgorió, J.-f. Arnoldi, M. Barbier, and S. Kéfi, Ecology Letters 27, e14413 (2024).

[30] E. Mallmin, A. Traulsen, and S. De Monte, Proceedings of the National Academy of Sciences 121, e2312822121 (2024).

[31] G. Aguadé-Gorgorió and S. Kefi, Journal of Physics: Complexity (2024).

[32] B. Rosenbaum and E. A. Fronhofer, Ecosphere 14, e4503 (2023).

[33] S. Arya, A. B. George, and J. P. OâDwyer, Proceedings of the National Academy of Sciences 120, e2307313120 (2023).

[34] R. M. May, Nature 238, 413 (1972).

[35] M. Barbier, J.-F. Arnoldi, G. Bunin, and M. Loreau, Proceedings of the National Academy of Sciences 115, 2156 (2018).

[36] J. Camacho-Mateu, A. Lampo, M. Sireci, M. A. Muñoz, and J. A. Cuesta, Proceedings of the National Academy of Sciences 121, e2309575121 (2024).

[37] J. Pasqualini, A. Maritan, A. Rinaldo, S. Facchin, E. Savarino, A. Altieri, and S. Suweis, arXiv preprint 2406.07465 (2024).

[38] J. E. Goldford, N. Lu, D. Bajić, S. Estrela, M. Tikhonov Sanchez-Gorostiaga,, D. Segré, P. Mehta, A. Sanchez, Science 361, 469 (2018).

[39] S. Allesina and S. Tang, Nature 483, 205 (2012).

[40] F. Koch, A.-M. Neutel, D. K. Barnes, and K. T. Allhoff, bioRxiv, 2024 (2024).

[41] Z. R. Miller and D. Max, bioRxiv, 2025 (2025).

[42] The collectivity limit is not equivalent to May’s classical limit on stability (May, 1972). A community can be stable within May’s bounds, but have strong indirect effects lea ding to EC (ESM XXX, Zelnik et al. 2024).

[43] A. Edelman, SIAM journal on matrix analysis and applications 9, 543 (1988).

[44] L. N. Trefethen and D. Bau, Numerical linear algebra (SIAM, 2022).

[45] W. Gilpin, arXiv preprint arXiv:2403.19186 (2024).

[46] J. M. Dambacher, H.-K. Luh, H. W. Li, and P. A. Rossignol, The American Naturalist 161, 876 (2003).

[47] S. Allesina and J. M. Levine, Proceedings of the National Academy of Sciences 108, 5638 (2011).

[48] R. A. Laird and B. S. Schamp, The American Naturalist 168, 182 (2006).

[49] S. Soliveres, F. T. Maestre, W. Ulrich, P. Manning, S. Boch, M. A. Bowker, D. Prati, M. Delgado-Baquerizo, J. L. Quero, I. Schöning, et al., Ecology letters 18, 790 (2015).

[50] S. Estrela, J. C. Vila, N. Lu, D. Bajić, M. Rebolleda-Gómez, C.-Y. Chang, J. E. Goldford, A. Sanchez-Gorostiaga, and Á. Sánchez, Cell Systems 13, 29 (2022).

[51] M. Barbier, C. De Mazancourt, M. Loreau, and G. Bunin, Physical Review X 11, 011009 (2021).

[52] M. Ghoul and S. Mitri, Trends in microbiology 24, 833 (2016).

[53] O. Artime and M. De Domenico, Philosophical Transactions of the Royal Society A 380, 20200410 (2022).

[54] R. M. May, Nature 261, 459 (1976).

[55] G. Boffetta, M. Cencini, M. Falcioni, and A. Vulpiani, Physics reports 356, 367 (2002).

[56] E. A. Bender, T. J. Case, and M. E. Gilpin, Ecology 65, 1 (1984).

[57] K. Kawatsu, Proceedings of the National Academy of Sciences 121, e2322939121 (2024).

[58] A. Eng and E. Borenstein, Current opinion in biotechnology 58, 117 (2019).

[59] R. Solé, V. Maull, D. R. Amor, J. P. Mauri, and C.-P. Núria, ACS Synthetic Biology (2024).

