## Supplementary Material for "Emergent coexistence and the limits of reductionism in ecological communities"

(Dated: May 15, 2025)

The structure of the following Supplementary Material is as follows: First, we present and define the fundamental problem under study and describe the numerical simulations and mathematical methods to explore it. Second, we present a detailed analysis of the results presented in the main text together with additional mathematical developments, figures and considerations. Third, we present a discussion that extends the conclusions of the main text.

All codes discussed in the present Supplementary Material are available at:

https://github.com/GuimAguade/EmergentCoexistence.

### CONTENTS

I. METHODS AND PRELIMINARY CONSIDERATIONS 4

A. Description of the problem and the GLV model 4

B. Pairwise exclusions, equilibria, net interactions and indirect effects 5

1. Pairwise exclusions 5

2. Indirect effects in three-species toy models 6

3. Equilibria, net interactions and the Jacobian matrix 9

C. Species interactions from random distributions 11

D. Network structure and connectivity in microbial communities 14

E. Finding stable communities 16

1. Simulating ecological dynamics 16

2. Sampling and testing random species subsets 17

F. Evaluating emergent coexistence and the fraction of excluding pairs (Figure 2) 19

G. Gathering data on the fraction of excluding pairs (Figure 2C) 20

H. Measuring predictability properties of the community $(\phi, \kappa)$ (Figure 3) 24

I. Designing empirical tests on predictability 25

J. Measuring May's threshold for stability in smaller systems 26

K. The Routh-Hurwitz criteria 27

L. Competitive hierarchies and intransitivity (Figure 4) 31

1. Competitive ranks and Low Rank Exclusion 32

2. Competitive triplets and Rock-Paper-Scissors 33

3. Other metrics of intransitivity 34

4. Building Figure 4 35

M. Nested interactions, single-resource competition and cross-feeding 36

N. Other dynamical models 40

1. Saturating interactions with a Holling response 41

2. Multilayer interactions with Allee Effects 42

3. Sublinear growth dynamics coupled to linear interactions 43

II. MAIN RESULTS AND ADDITIONAL OUTCOMES 45

A. Mathematical results for emergent coexistence in the Generalized Lotka-Volterra phase space 45

1. The minimal competition necessary for emergent coexistence 45

2. The maximum competition allowed for emergent coexistence 47

3. The maximum fraction of excluding pairs that a community can sustain 49

4. Emergent coexistence under other interaction matrices 53

5. Emergent coexistence in other dynamical models with pairwise interactions 54

6. Sparse interactions and species richness 58

B. Indirect effects, collectivity and condition number 60

1. Interaction patterns in $A^*$ and coexistence 60

2. All-to-all competition but positive net effects 62

3. Indirect effects and an estimate for minimal and maximal collectivity 63

4. An estimate for minimal condition number $\kappa$ 65

5. Empirical applications of using $\phi$ and $k$ to infer predictability 69

C. Feedback loops and stability 70

D. Intransitivity and emergent coexistence 72

1. Low rank exclusions require 4 exclusions, and even then are rare 72

2. Rock-Paper-Scissors loops are equivalent to a random expectation 73

3. Equivalent results with the Laird-Schamp index 74

E. On the role of migration and reinvasions 75

F. Extended Discussion 76

G. Emergent coexistence from chains of pairwise interactions 76

H. Emergent coexistence is an emergent phenomenon 76

I. The limits of bottom-up predictability in species coexistence 77

J. Incorporating non-random and non-pairwise interactions 78

K. Conclusion 78

Figures 80

Bibliography 94

References 94

### I. METHODS AND PRELIMINARY CONSIDERATIONS

#### A. Description of the problem and the GLV model

The original question of this work can be expressed in the following way: *Will a species-rich community model with pairwise interactions between species generate stable states, in which some of the constituent species do not coexist in co-culture?*

The motivation comes from recent work in multispecies microbial communities, demonstrating that stable communities can harbor coexisting pairs of species that do not coexist when isolated from the community due to strong competition (Chang *et al.*, 2023; Friedman *et al.*, 2017; Higgins *et al.*, 2017; Lele *et al.*, 2024; Venturelli *et al.*, 2018). Because coexistence of this excluding pairs is not expected from co-culture observations, this phenomenon has recently been labeled as *Emergent Coexistence* (EC) (Chang *et al.*, 2023).

Our first question is then followed by other corollary questions: *(i) If a model of pairwise interactions can generate EC, what mechanisms allow these excluding pairs to coexist in a larger community?* and *(iii) Given these unexpected observations of an emergent community phenomenon, can we really infer species coexistence and community composition from pairwise information alone?*

To tackle these questions, we study the generalized Lotka-Volterra (GLV) model as the simplest possible model for multispecies pairwise interactions, which has recently received a considerable amount of attention and progress (see e.g. (Aguadé-Gorgorió and Kefi, 2024; Altieri *et al.*, 2021; Barbier *et al.*, 2018; Bunin, 2017; Galla, 2018; Kessler and Shnerb, 2015; Mallmin *et al.*, 2024; Serván *et al.*, 2018; Zelnik *et al.*, 2024)). In one of the several variations of this model, the abundance of a given species $N_i$ follows

$$\frac{dN_i}{dt} = r_i N_i \left( 1 - \frac{N_i}{K_i} + \sum_{j \neq i}^{S} a_{ij} N_j \right). \tag{1}$$

where $r_i$ is the intrinsic growth rate, $K_i$ the intrinsic carrying capacity and $a_{ij}$ is an interaction term describing how the growth of species' abundance $N_i$ is affected by the abundance of any other species $N_j$. As described for example in (Aguadé-Gorgorió and Kefi, 2024; Mallmin *et al.*, 2024; Zelnik *et al.*, 2024), it is often interesting to rescale the parameters of this equation so that we can focus on the role of interactions $a_{ij}$.

To do so, a typical procedure is to assume homogeneous growth rates $r_i = r$ and rescale time as $t \to rt$. It will also become interesting to work with adimensional units, as explained in detail in (Zelnik *et al.*, 2024) and in the section discussing indirect effects and collectivity below. To do so, we rescale interaction strengths relative to self-regulation ($A_{ij} \equiv a_{ij}/a_{jj} = a_{ij}K_j$). $A_{ij}$ therefore captures how much interactions with individuals of another species affect growth, compared to interactions with individuals of the same species. Such a notion of inter- vs intra-species interactions is on its own a central topic in theoretical ecology (Barabás *et al.*, 2017; Hatton *et al.*, 2024).

It is important to remark that, in our specific notation and that of (Zelnik *et al.*, 2024), the matrix $A$ contains the off-diagonal, interspecies interactions, whereas it has zeroes in the diagonal. The complete matrix of interactions, containing both self-regulation and inter-species interactions, is therefore $-I + A$, where $I$ is the identity matrix where the diagonal is $A_{ii} \equiv a_{ii}/a_{ii} = 1$. This specific choice of notation does not have any specific implications, but will become useful later on once we want to invert the matrix $(I - A)^{-1}$ and find ourselves facing the Neumann series (Zelnik *et al.*, 2024).

Now the GLV model describes the dynamics of the relative yield of a species (see (Lajaaiti *et al.*, 2024)) as

$$\frac{dx_i}{dt} = x_i \left( 1 - x_i + \sum_{j \neq i}^{S} A_{ij} x_j \right). \tag{2}$$

In this context, the central question becomes: can this model generate stable communities of multiple coexisting species, in which some species pairs do not coexist in co-culture? If so, additional questions are in place. If it does not predict EC, the research should likely move into adding so-called higher-order effects (Billick and Case, 1994; Gallien *et al.*, 2017; Grilli *et al.*, 2017b).

#### B. Pairwise exclusions, equilibria, net interactions and indirect effects

##### 1. Pairwise exclusions

A starting point in the problem above is to understand the conditions by which a pair of interacting species coexists or undergoes exclusion in the absence of any additional species. This is a classical and well-known problem in theoretical ecology (Chesson, 2000; Strogatz, 2018).

Following the notation above, the question is framed in understanding the outcomes of

$$\frac{dx_1}{dt} = x_1 \left( 1 - x_1 + A_{12} x_2 \right) \tag{3}$$

$$\frac{dx_2}{dt} = x_2 \left( 1 - x_2 + A_{21} x_1 \right) \tag{4}$$

It is easy to show that the Jacobian of this system at an equilibrium $(x_1^*, x_2^*)$ is

$$J = \begin{pmatrix} 1 - 2x_1^* + A_{12} x_2^* & x_1^* A_{12} \\ x_2^* A_{21} & 1 - 2x_2^* + A_{21} x_1^* \end{pmatrix}$$

The two exclusion fixed points are $(x_1^* = 1, x_2^* = 0)$ and $(x_1^* = 0, x_2^* = 1)$. For the first, eigenvalues follow from solving (Strogatz, 2018)

$$\det(J - \lambda I) = -(1 + \lambda)(1 + A_{21} - \lambda) = 0. \quad (5)$$

We find $\lambda_1 = -1$ and $\lambda_2 = 1 + A_{21}$. The state will be stable under the well-known result of $\lambda_2 = 1 + A_{21} < 0$, which gives $A_{21} < -1$. Species 1 will exclude species 2 if competition is stronger (more negative) than self-regulation (take into account the original scaling, $A_{ij} \equiv a_{ij}/a_{ii} = a_{ij}K_j$), a classical and somehow intuitive result. The same can be said about the symmetric fixed point where species 2 excludes species 1 and, on the contrary, about the coexistence state. Exclusion of species $i$ by species $j$ happens for $A_{ij} < -1$, while coexistence of both species happens if both $A_{ij}$ and $A_{ji}$ are larger than -1 (Strogatz, 2018). Bistable exclusion, where both outcomes are possible and depend on what species starts with advantage in terms of abundance, is possible provided that both $A_{ij}$ and $A_{ji}$ are more negative than -1 (Aguadé-Gorgorió *et al.*, 2024b). This frames the question above slightly better: can the GLV model generate stable communities of multiple coexisting species, in which some species pairs $i, j$ do not coexist in co-culture because $A_{ij}$ or $A_{ji}$ are smaller than -1? From here onwards we will call these species pairs *excluding pairs*, in the sense that they exclude one another when grown outside of the community context.

### 2. Indirect effects in three-species toy models

As discussed in the main text, our central argument is that indirect effects can, on their own, generate stable communities with excluding pairs and therefore EC. While intransitive competition or higher-order effects might be present, they are not necessary, and only strong indirect effects need to be in place. Therefore it seems natural to continue by introducing what indirect effects are and what do we know about how to analyse them.

Simple, few-species models provide an easy way to understand the potential link between indirect effects and emergent coexistence, and numerous research in the last decades has used these methods in more or less mathematically-oriented works to uncover the role of indirect effects in simple-enough systems when more than two species are considered (Bender *et al.*, 1984; Dambacher *et al.*, 2003; Levine, 1999, 1976; Levins, 1974; Strauss, 1991; Vandermeer, 1980; Wootton, 1994; Yodzis, 1988). A great general discussion on these topics as well as intransitivity and higher-order interactions can be found in (Levine *et al.*, 2017). Let us try here to describe a minimal scenario for how emergent coexistence can be driven by indirect effects in simple and tractable three-species cases. Suppose two species, 1 and 2, of which we know that 2 can exclude 1, while 1 has for example no impact on 2. This could be a limit-case scenario for two species that compete for the same resource, where 2 is a much better consumer than 1, to the point that the consumption rate of species 1 is negligible from species 2. Another great review discussing the links between such a mechanistic description of interactions through resources and a phenomenological description of direct, species-level interactions can be found in (van den Berg *et al.*, 2022). In any case, this competitive scenario translates into $a_{12} < a_{11} < 0$, so that $A_{12} < -1$. Now, a third species 3 competes with both of them, but is not affected by them. This imposes a perfectly nested interaction matrix $A$, where 3 competes with 1 and 2, 2 excludes 1, and 1 has no impact on others, such as those discussed in Section I.M. If each species replicates and competes following a Lotka-Volterra model, the system of equations for the abundance of each species $x_i$ follows this simple structure:

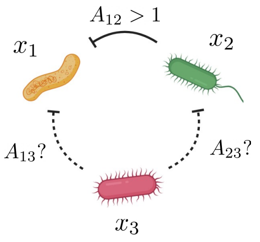


$$A = \begin{pmatrix} 0 & A_{12} & A_{13} \\ 0 & 0 & A_{23} \\ 0 & 0 & 0 \end{pmatrix}$$

$$\frac{dx_1}{dt} = x_1\left(1 - x_1 - A_{12}x_2 - A_{13}x_3\right)$$

$$\frac{dx_2}{dt} = x_2\left(1 - x_2 - A_{23}x_3\right)$$

$$\frac{dx_3}{dt} = x_3\left(1 - x_3\right)$$

In this toy model we have written $A_{ij}$ competition terms with a minus sign in front. In general we prefer writing interactions as $+A_{ij}$, meaning that a positive element means cooperation, and a negative element means competition. Yet, for the sake of this single specific example, we use the opposite notation, which will be useful for the reader to visualize the game of signs that emerges with indirect effects. This means that here positive values of $A_{ij}$ imply competitive interactions, and so $A_{ij} > 1$, because of the negative sign in front of it, implies competitive exclusion. Remember again that the matrix of interactions $A$ contains the off-diagonal elements by the way we define it, so that we include negative self-regulation as an identity matrix or simply $-1$ elements *outside*, which will become useful below. The simplicity of this example makes it easy to isolate the equilibrium abundances of the coexistence state by equating all dynamical equations to zero:

$$x_1^* = 1 - A_{13} - A_{12} + A_{12}A_{23}; \qquad x_2^* = 1 - A_{23}; \qquad x_3^* = 1 \tag{6}$$

This result clearly shows the simplest possible scenario in which indirect effects emerge in competitive models, and is a limit-case scenario of the much more complex equation (4) of the main text. The equilibrium abundance of species 1 is reduced by direct competition, $-A_{12}$ and $-A_{13}$, but improved by the positive effect of the 3→2→1 interaction chain, $A_{12}A_{23}$. Here is where the sign convention used for this specific example allows for a visual understanding of the presence of positive indirect effects

Under which conditions can species 3 rescue species 1 from extinction, without excluding species 2? In other words, under which conditions will all three species abundances at equilibrium in the previous equation be positive? This happens provided

$$A_{23} < 1; \qquad A_{13} < 1 - A_{12}(1 - A_{23}). \tag{7}$$

In other words, feasibility (a term used to describe a scenario where all species abundances are positive, (Grilli *et al.*, 2017a; Marcus *et al.*, 2024)) requires $A_{23} \lesssim 1$ to repress species 2 without excluding it, and $A_{13} \gtrsim 0$ to avoid excluding species 1. The stronger

$A_{12}$ is, the more pressure on species 1 due to competition. This implies that a stronger $A_{12}$ means that species 3 needs to repress species 2 even more, and repress species 1 much less. Mathematically, as competition $A_{12}$ becomes stronger, the more different $A_{13}$ and $A_{23}$ have to be to guarantee the coexistence of the trio. This provides a clear intuition of why interaction heterogeneity ($\sigma$) is required in order to ensure positive indirect effects and emergent coexistence. Let us assume that both interaction terms $A_{13}$ and $A_{23}$ are sampled e.g. from a uniform distribution $\mathcal{U}(c-w, c+w)$, where $c$ is the center of the distribution and $w$ its half-width. Then, in the best case scenario to ensure coexistence, $A_{13} = c - w$ and $A_{23} = c + w$. Plugging this best case expectation in the conditions for coexistence described above, we find that coexistence is ensured provided that the width of the distribution (the heterogeneity of interactions) is larger than a given limit,

$$w > (1-c)\frac{A_{12}+1}{A_{12}-1}. \tag{8}$$

This means that, for coexistence to be possible, there is a necessary heterogeneity $w$ in the impacts of species 3 on 1 and 2. If both interactions had the same strength, coexistence would fail and species 1 would go extinct. Overall, this first simple test provides an intuitive notion of why interaction heterogeneity is required to observe emergent coexistence. This is because heterogeneous interactions can counteract the impacts of competition by generating positive indirect effects. Interestingly, this condition can be tested by generating such a simple model numerically: by sampling interactions as proposed and asking for three-species coexistence, one can easily see there is a minimal heterogeneity in the effects of species 3 to ensure this simple example of emergent coexistence (not shown).

This simple picture provides a starting point for our discussion: First, it shows that emergent coexistence requires a certain degree or amount of interaction heterogeneity, thanks to which chains of interactions can emerge and maintain excluded species within the community. Furthermore, it fosters the later intuition by which intransitivity might not be necessary for emergent coexistence, as indirect effects can be simply linear and come from a higher ranked species without necessarily requiring more complicated or fine-tuned architectures: note that the interaction matrix of this system is perfectly triangular and there are no rock-paper-scissors loops nor upwards exclusion effects. This will be a toy model for the nested matrix $N^*$ studied in later parts of this work.

Finally, this toy model recapitulates the importance of weak effects in the sense of competitive but non-exclusionary interactions. Empirically, weak links might appear irrelevant, such as when observed in pairwise coexistence between two species that do not seem to play a strong effect on one another (Chang *et al.*, 2023). Yet, as empirically explored in (Neutel *et al.*, 2002) and described mathematically throughout our work, weak interactions can still be fundamental to community behavior, functioning or coexistence, by being key elements in longer chains of collective effects. As discussed in Section I.L, the role of weak effects (or, actually, their absence), also explains why earlier work on intransitivity using tournament matrices with binary interactions are not able to capture the nuanced role of chains of indirect effects (Allesina and Levine, 2011; Laird and Schamp, 2006).

This three-species example is also simple thanks to the fact that half of the network of interactions is sparse, meaning that most links are not realized and most chains of indirect effects are therefore short. Yet, how do indirect effects scale up as we consider more interacting species and less zeros in their interaction matrices? It is easy to see that the number of potential links and chains will increase rapidly as one considers more species interacting via a non-triangular interaction matrix. In the main text and throughout this supplementary material we study recent developments that provide a framework to characterize indirect effects in such high-dimensional systems (Zelnik *et al.*, 2024).

#### 3. Equilibria, net interactions and the Jacobian matrix

The GLV model provides potentially the simplest description of a community build upon pairwise interactions between species. One of the benefits of this simple description is that bilinear mass-action interactions, in the form of $x_i A_{ij} x_j$, make the search for equilibria a linear problem. Given the original equation

$$\frac{dx_i}{dt} = x_i \left( 1 - x_i + \sum_{j \neq i}^{S} A_{ij} x_j \right), \tag{9}$$

a fixed point or equilibrium state of the system with $S$ species will fulfill

$$x_i \left( 1 - x_i + \sum_{j \neq i}^{S} A_{ij} x_j \right) = 0 \qquad \forall i \in S. \tag{10}$$

As for the 2-species case above, this can be fulfilled by species being extinct, $x_i^* = 0$, or by the term inside parenthesis being zero. If we study the case where a species has not gone extinct and so we search for its positive abundance, the condition for the equilibrium writes as

$$\left( 1 - x_i + \sum_{j \neq i}^{S} A_{ij} x_j \right) = 0, \tag{11}$$

and, in vector notation, becomes

$$1 - (I - A)\boldsymbol{x} = 0 \tag{12}$$

where $A$ has zeros in the diagonal, so that self-regulation is captured by the identity matrix $I$ alone. This allows us to isolate

$$(I - A)\boldsymbol{x} = 1. \tag{13}$$

We highlight here a key point in our work and notation consistent with previous works (Aguadé-Gorgorió and Kefi, 2024; Liautaud *et al.*, 2019). Not all of the $S$ species of the original pool will survive. In fact, in the regime where we find EC, previous work has shown that there can exist a multiplicity of stable states, all with different subcombinations of the original pool of species. The question, therefore, is to find the stable equilibrium abundances of species that can survive together. As discussed later on, this is a particular scenario for the GLV model without external migration, in which species go extinct for good (Liautaud *et al.*, 2019; Mallmin *et al.*, 2024). The effect of species migration and reinvasion in the community is discussed in section II.E. We note this final subset of coexistors with a $^*$ as in (Liautaud *et al.*, 2019), while it was noted with a $'$ in (Aguadé-Gorgorió and Kefi, 2024). The question is to find the equilibrium abundances for a subset of $S^*$ species, $x^*$, so that the subset of interactions between these species $A^*$ yields all-positive abundances in the equation

$$\boldsymbol{x^*} = (I - A^*)^{-1}1. \tag{14}$$

In this equation and because of the vector product with the unitary vector 1, the row sums of $(I - A)^{-1}$ are the equilibrium abundance of each species, yielding the key result:

$$x_i^* = \sum_j (I - A^*)_{ij}^{-1}. \tag{15}$$

Now, an equilibrium state might have some species extinct, and others present and interacting. In fact, we know that coexistence of all species is a rare scenario at very low interaction strength (Bunin, 2017). As set before and following our notation, finding an equilibrium state of the GLV model then accounts for finding a subset of $S^* \leq S$ species, interacting via a subset of the original matrix $A^* \in A$, for which the row sums of $(I - A^*)^{-1}$ are all positive. In this scenario, $S - S^*$ species have gone extinct, and the remaining $S^*$ can coexist via their self-regulation (the identity matrix, $I$) and the effects of all the remaining interactions.

As described in different degree of detail in e.g. (Arnoldi *et al.*, 2022; Levine, 1999, 1976; Liautaud *et al.*, 2019; Strauss, 1991; Wootton, 1994; Zelnik *et al.*, 2024), the matrix $(I - A^*)^{-1}$ then captures the net effects between species, described as the long-term change in the stable abundance of a species, $x_i^*$, due to a change in the growth rate of another species $j$. We refer the reader to those works for deeper discussion on the ecology and mathematics behind direct and net effects.

This is a fundamental component of ecological communities of many interacting species: the matrix $A^*$ captures the interactions between each pair of species, which we can obtain by isolating species and comparing their growth-when-alone to the growth in pairwise coculture e.g. as described in (Pennekamp *et al.*, 2018; Venturelli *et al.*, 2018). Yet, the stable state of the system once we consider multiple species is not a direct function of $A^*$, but of the net interactions $(I - A^*)^{-1}$. Again, the linearity of the GLV model provides a powerful way of understanding this concept. As shown in (Zelnik *et al.*, 2024), net effects can be decomposed using Neumann's series

$$(I - A^*)^{-1} = 1 + A^* + (A^*)^2 + (A^*)^3 + ... \tag{16}$$

which, as discussed in the main text, implies that the stable abundance of a given species is

$$x_i^* = 1 + \sum_j A_{ij}^* + \sum_{j,k} A_{ik}^* A_{kj}^* + \sum_{j,k,l} A_{ik}^* A_{kl}^* A_{lj}^* + ... \tag{17}$$

Now, the abundance of species $i$ is not only a function of direct interactions with all interaction *neighbors* (species connected by direct pairwise interactions, not neighbors in space). Rather, the abundance of species $i$ is also affected by how other species interact with its direct neighbors, and so on and so forth. This also includes closed chains, such as the effect on species $i$ on others, these on others, and these others back on species $i$, which will appear in the diagonal elements of $(A^*)^k$. The $(I - A^*)^{-1}_{ij}$ element of the matrix of net effects accounts for how $j$ affects $i$ not only by the direct, immediate interaction $A^*_{ij}$, but also through all possible chains of interactions across the community. This notion of indirect effects through the community is a fundamental property of complex systems and is at the core of our work.

Given an equilibrium state, characterized by a set of surviving species with abundances $x_i^*$ and interactions $A^*$, it is easy to show that the Jacobian matrix has elements $J_{ij} = x_i^* A_{ij}$ for $i \neq j$ and $J_{ii} = -x_i^*$, or, grouping the two matrices, $J_{ij} = x_i^*(-I + A^*)_{ij}$. From this, one can evaluate if the equilibrium state is linearly stable by finding the eigenvalues of this Jacobian matrix and evaluating if they are all negative, a topic with a long history in theoretical ecology (Allesina and Tang, 2012; May, 1972). Linear stability and the sign and value of eigenvalues provide a measure of the response of a fixed state to infinitesimal perturbations, with negative eigenvalues indicating that the system will recover its original state. We do not delve into different types of perturbations or other topics around ecological stability or resilience in our work, and we derive the reader to deeper discussions on the multidimensional meaning of ecological stability beyond linearization/eigenvalue methods in e.g. (Domínguez-García *et al.*, 2019; Donohue *et al.*, 2016; Kéfi *et al.*, 2019; Lajaaiti *et al.*, 2024).

#### C. Species interactions from random distributions

Given the parametrization we propose, the dynamics of the GLV model under study are fully governed by the off-diagonal values of the interaction matrix $A$. Yet, how can we parameterize this model? As discussed throughout this work, inferring all $A_{ij}$ parameters from time series analysis can represent an extremely complex task, even in systems with a handful of species, where the number of parameters to fit (the dimension of the interaction matrix without diagonal elements) scales as $S(S-1)$ if we are able to access $A_{ij} = a_{ij}/a_{ii}$ directly (Picot *et al.*, 2023; Rosenbaum and Fronhofer, 2023). Even more so, recent research discusses how this process could even be problematic by definition, with multiple combinations of parameters providing a good fit for the same time series (Lubiana Botelho *et al.*, 2025).

Instead, accumulated work has proposed that, even when quantitative predictions might be hard or impossible, we can gain qualitative predictions of species-rich dynamics by assuming that interactions are drawn from a random probability distribution. That is, assuming that we do not know all the details of $A$, but only its aggregate properties. Here we build on previous work that has already discussed how using random interactions can provide a good starting point or null assumption to start exploring the dynamical properties of a community model (Aguadé-Gorgorió *et al.*, 2024b; Barbier *et al.*, 2018,

2021; Bunin, 2017; May, 1972).

At its core, the disordered systems approach assumes that, due to the complexity of ecological communities, individual species and their interactions cannot be assigned specific roles (Barbier *et al.*, 2021). Instead, it focuses on broad characteristics of the interaction matrix, such as the average strength of interspecies interactions and their variability (Barbier *et al.*, 2018). This lack of imposed structure helps identify fundamental patterns that emerge in high-dimensional systems and distinguishes them from behaviors that require additional complexity, which is in fact the aim of this work.

Because most of the EC experimental observations do not harbor specific estimates of species interaction strengths (Chang *et al.*, 2023; Friedman *et al.*, 2017), the disordered approach becomes a useful starting point for our research: rather than exploring specific interaction strengths inferred from experiments, together with empirical ecological network structures like food-web architecture (Dunne, 2006) or mutualistic interactions (Bascompte, 2009; Payrató-Borras *et al.*, 2019; Suweis *et al.*, 2013), we can first ask, *Can the simplest community model predict emergent coexistence in the absence of knowledge of specific interaction structures? Or else, do we need to impose specific properties in A to observe EC?* This question can be followed by questions in the structure of the emerging network of interactions between surviving species, as recently studied in (Barbier *et al.*, 2021; Kessler and Shnerb, 2025) *Will we observe specific structures (e.g. Rock-Paper-Scissors) emerging naturally from the interaction dynamics? Or else, do we need to invoke additional non-random components to recover empirical observations of transitivity or EC?*

A key foundation for this perspective comes from Robert May's seminal work (May, 1972), which introduced the idea of treating interaction strengths in species-rich models as random parameters. Using Random Matrix Theory, this approach has provided a mathematical framework for studying ecological community dynamics using species-rich models, with particular emphasis on understanding stability (Allesina and Tang, 2012; May, 1972) or species coexistence (Barbier *et al.*, 2021; Bunin, 2017; Grilli *et al.*, 2017a). Despite the approximation of random interactions is obviously very strong and must be used with caution, we consider it a valuable tool, at least (but not only) to overcome the issue of parametrization.

Furthermore, as we have discussed above and along the work, what we are sampling at random is not the interactions within a stable community, but rather the interactions between a much larger pool of e.g. a hundred species from which stable states can possibly emerge. This is also useful because it allows us to uncover patterns that result from the ecological process (Barbier *et al.*, 2021; Kessler and Shnerb, 2025): for example, if triplets of competitors sampled from random distributions always hold a non-random Rock-Paper-Scissors architecture, it points towards the ecology benefiting such given structure. Such a comparison between emerging properties and a basal or random null model is at the core of the ecological thinking discussed earlier in (Hutchinson, 1953).

In our work, the direct effect of the abundance of species $j$ on the growth of species $i$ is mediated by $A_{ij}$, which are sampled from a Gaussian distribution with mean $\mu$ and standard deviation $\sigma$ following the classical approach of recent works (Altieri *et al.*, 2021; Bunin, 2017) but in the case where interactions are not rescaled by species diversity and so what we believe is the case with minimal possible assumptions (Aguadé-Gorgorió *et al.*, 2024b; Aguadé-Gorgorió and Kefi, 2024; Mallmin *et al.*, 2024). One of the advantages of this methodology is that one can draw a two-dimensional phase plane of the GLV model solely by modifying $\mu$ and $\sigma$ and find that four different and interesting dynamical phases exist (Aguadé-Gorgorió and Kefi, 2024). In figure 2A of the main text, we take advantage of recent work studying this phase space with weak (Bunin, 2017) and strong (Aguadé-Gorgorió and Kefi, 2024; Kessler and Shnerb, 2015; Mallmin *et al.*, 2024) interactions. In particular, we set $\mu \in [-2.00, 0.5]$ for a range of interaction strengths between strong competition and weak cooperation, and $\sigma \in [0, 1]$. These already allow us to visualize the whole domain of different possible dynamical behaviors (Fig. 2A in the main text). Again, self-regulation is considered outside of the matrix and implemented as an identity matrix, which has been used in (Zelnik *et al.*, 2024) as a means to rewrite species abundances and hence net effects in terms of a sum of indirect effects via the Neumann series (see section above). The random components therefore are present in interspecies interactions only.

Together with the null assumption of completely random and uncorrelated interactions discussed in the main text, we study the presence of EC in other matrix randomizations, inspired in the works of e.g. (Allesina and Tang, 2012; Bunin, 2017; Grilli *et al.*, 2017a, 2016) where different structures of random Jacobian and interaction matrices are studied. In particular we study if the GLV system predicts the presence of communities with emergent coexistence under fully symmetric ($A_{ij} = A_{ji}$) and anti-symmetric predator-prey-like ($A_{ij} = -A_{ji}$) random matrices. To build them, we generate an upper triangle from a normal distribution as discussed above and then generate the lower triangle according to these rules and the values of the upper triangle.

We also study the effects of matrix triangularity (or nestedness) and Connectivity (see below). The first is generated fixing the probability $p$ that a given value in the lower triangle $A_{ij}$, $i > j$ equals zero, so that a large $p$ value implies a high degree of nestedness. The second is generating by fixing the connectivity $C$, so that there is a probability $1 - C$ that whatever $A_{ij}$ value equals zero.

We also test the effects of a more nuanced interaction matrix involving single-resource competition, a weak random competition term and cross-feeding, as described in the main text and in Section I.M below. We finally test the effects of a more skewed distribution of competitive interactions following the recent observations described in (Koch *et al.*, 2024). This is done using a Gamma distribution using numpy's built in `numpy.random.gamma` function, for which we can control the shape and scale parameters and has a longer negative tail, so that there are many weak competitors but also a few much stronger ones. The results regarding EC in these alternative matrices are discussed in Section II.A.4.

Additionally, one could consider that the regularization technique proposed in the beginning of this work, by which interactions are modeled relative to self-regulation ($A_{ij} \equiv a_{ij}/a_{jj} = a_{ij}K_j$), could induce certain correlations betwen interaction strength and species traits or *fitness* captured by the carrying capacity $K_i$. The importance of these correlations in theoretical and experimental systems has been very recently discussed in (Pearl Mizrahi *et al.*, 2025). In this work, the authors propose that correlations between interaction strength and species carrying capacities can impact the response of a community to secondary extinctions. Moreover, these correlations increase competitive hierarchy (interaction strengths are ordered by carrying capacities), leading to equivalent results as those discussed in the sections below regarding hierarchy, transitivity and emergent coexistence.

Finally, it is noteworthy to mention that we do not aim at using more detailed or empirical interaction matrices. This is because our work aims at providing a set of qualitative results that are general enough and not overly dependent on system particularities. Moreover, one of the conclusions of our work hints towards the possibility that interaction parameters are not only very hard to obtain (Lubiana Botelho *et al.*, 2025; Picot *et al.*, 2023; Rosenbaum and Fronhofer, 2023), but that the bottom-up or reductionist method might have limitations. In this regard, we leave for future analysis the application of our results on empirically obtained interaction matrices, and refer the reader to a large set of works that try to obtain $A_{ij}$ or $J_{ij}$ values (or that at least provide useful data towards this) for microbial ecosystems (Arya *et al.*, 2023; Camacho-Mateu *et al.*, 2024; Castledine *et al.*, 2024; Ortiz *et al.*, 2021; Pennekamp *et al.*, 2018; Rao *et al.*, 2021; Schmitz *et al.*, 2024; Venturelli *et al.*, 2018).

It is alos worth noting that a large body of literature has studied a very similar problem of pairwise-to-community patterns in the case of plant communities, although we cannot claim if our work applies in that context, with the low dynamical responses of plants compared to bacteria and the general lack of explicit competitive exclusion being potential limitations. It could well be that interactions in plant communities are considerably weaker than those of microbial ecosystems, leading therefore to lower collectivity values and weaker indirect effects. We refer the reader to some works within this large set of plant community experiments and discussions on the applicability of GLV-like models to describe them (Dormann, 2007; Dormann and Roxburgh, 2005; Engel and Weltzin, 2008; Roxburgh and Wilson, 2000).

#### D. Network structure and connectivity in microbial communities

In a certain way, one can decouple the strengths of interactions, which are sampled as random parameters as described above, from the structure of these interactions (Grilli *et al.*, 2016; Poley *et al.*, 2025). This means that we can build interactions as the so-called *adjacency* matrix of who interacts with who, $M$, multiplied by the matrix of the strength of these interactions, $A$. This multiplication is of course done element-wise with the Hadamard product. Throughout this work, $A \circ M$ is called $A$, mostly because $M$ plays a secondary role as discussed below and in section II.A.6.

Taking advantage from the theory of graphs and the revolution of complex network methods, there has been extensive work in the last decades regarding the structure of ecological networks (Bascompte, 2010; Guimaraes Jr, 2020). In particular, we now know that mutualistic networks –encapsulating for example positive interactions between plants and their pollinators– can be strongly nested, whereas food-webs might be highly modular (Fortuna *et al.*, 2010). Moreover, theory suggests that, at least in principle, these structures could play a fundamental role in the functioning of ecological networks, providing communities with more robustness or resilience than what one could expect at random (Landi *et al.*, 2018). Yet, other models propose that structural patterns might not be *selected* or *beneficial* in terms of ecological dynamics, but rather they might be a byproduct of the assembly rules (Valverde *et al.*, 2018) or natural heterogeneity (Payrató-Borras *et al.*, 2019) of ecological communities. In any case, if and how non-random network structures play a pivotal role in ecological dynamics remain debated questions (Barbier *et al.*, 2018).

Conversely, accumulated work has not found strong evidence for non-random structures in microbial interaction networks beyond sparsity (Arya *et al.*, 2023; Camacho-Mateu *et al.*, 2024; Venturelli *et al.*, 2018). This, of course, remains an open avenue for future work, and it could be that the presence of modules or nestedness in microbial community networks depends on the type of microbiome under study and the presence of specific functional structures related to resource consumption or cross-feeding (see Section I.M or e.g. (Dal Bello *et al.*, 2021; Estrela *et al.*, 2022) for recent works on consumption structures). As discussed in the main work, for example, assembly of microbial consortia under a single resource might induce a nested and hierarchical competition matrix, whereas positive cross-feeding in this context might result in strong facilitation of otherwise weak competitors. Other scenarios of ecological mechanisms could result in other network structures.

For the main part of our work, however, we stick to recent evidence proposing that ecological networks of microbial interactions might not hold strong structures, and that sparsity might be the only highlighted property (Arya *et al.*, 2023; Camacho-Mateu *et al.*, 2024). We believe this is also in line with the above assumptions of a lack of structure in the distribution of $A_{ij}$. Network sparsity here refers to the fact that a large fraction of possible interactions between species are not realized. In mathematical terms, the adjacency matrix of species connectivity $M$ is filled with many zero elements. The usefulness of this assumption is two-fold: if microbial interactions do hold a certain degree of apparent randomness, our results and the overall disordered approach literature provide a starting point to discuss ecological dynamics. Instead, if certain microbial interaction networks are found to hold significant structural properties, the random matrix approach should at least provide a null model for comparison. Again, both the assumptions for $A$ above and for $M$ in this section provides an elegant take on how to have *a priori criteria as to the amount of structure it is reasonable to expect* (Hutchinson, 1953): as with the case of Rock-Paper-Scissors loops, the model with random adjacency matrices provides a baseline to investigate what collective properties emerge from randomness per-se (such as EC), and what properties require additional non-random structures to be observed. In this context, recent work on the interplay between microscopic disorder and macroscopic structure could provide a next step in our capacity to model the microbiome (Giral Martínez *et al.*, 2024).

Mathematically, we model the adjacency matrix $M$ and hence the interaction matrix $A \circ M$ as an Erdős-Rényi graph, which provides a classical and common model to build random networks with controled connectivity (Newman, 2018; Van Der Hofstad, 2024). In short, an Erdős-Rényi graph is build by defining a probability $p$ that two species interact. This can be implemented in several ways. We first build a matrix of normally distributed interactions with the `random.normal` function in `numpy`. Then, for each element, we keep it with probability $p = C$, where $C$ is the connectivity parameter, or set it to zero with probability $1 - C$.

In the main text we present results for the case with $C = 1$, meaning that all possible interactions are realized. Evidence for empirical communities indicates that this value might be quite large in some cases ($C \approx 0.77$ in (Venturelli *et al.*, 2018), $C \approx 0.9$ in (Koch *et al.*, 2023)), whereas it could be extremely lower in other ecosystems (Arya *et al.*, 2023; Camacho-Mateu *et al.*, 2024). In fact, it could well be that connectivity is an extremely variable parameter depending on the system under study: for example, experimentally assembled communities like the ones discussed in the main paper appear to hold quite densely connected matrices, with evidence of high fractions of strongly competing pairs, possibly due to the fact that species are grown on a single or a few resources and hence there are few available niches leading to a relatively well-connected matrix (Chang *et al.*, 2023; Friedman *et al.*, 2017; Venturelli *et al.*, 2018).

In any case, we discuss in section II.A.6 the effects of lowering connectivity below $C = 1$. Simply put, lowering connectivity alters the amount of species that can coexist on a given $\mathcal{N}(\mu, \sigma)$ distribution, because it reduces the effective competition strength while modulating heterogeneity non-monotonically (see Section II.A.6 and supplementary material of (Aguadé-Gorgorió and Kefi, 2024)). One possible corollary is that natural microbial ecosystems, which hold much higher species richness than what is observed in laboratory communities, could do so because the niche partitioning of resource-rich and heterogeneous environments implies that most interactions are weak or null: sparsity provides a simple and almost trivial way of potentially allowing increased diversity (Section II.A.6). In any case, we find that the qualitative results of EC and the role of indirect effects are the same for different connectivity values (Section II.A.4).

#### E. Finding stable communities

Given the mathematical framework and the parametrization techniques described above, our work uses two different methods to find equilibrium states in GLV communities given the only two parameters of mean and heterogeneity of interaction strengths $(\mu, \sigma)$. These methods have been used and described in detail in our previous works (Aguadé-Gorgorió *et al.*, 2024b; Aguadé-Gorgorió and Kefi, 2024).

##### 1. Simulating ecological dynamics

Given the GLV model of equation (2), one way of solving the problem is to simulate the ecological dynamics numerically. One simulation is defined by setting $\mu$ and $\sigma$ within the range of values of figure 2A in the main text either gradually (for figure 2A) or randomly with uniform probability within the same ranges for all other figures. We then generate a matrix of interactions $A$, with $A_{ij} \in \mathcal{N}(\mu, \sigma)$ with probability $C$ and $A_{ij} = 0$ with probability $1 - C$. We define random initial conditions from a uniform distribution $x_i(t = 0) \in \mathcal{U}[0.0, 1.0)$. Following (Aguadé-Gorgorió *et al.*, 2024b; Aguadé-Gorgorió and Kefi, 2024), an integration time of $t = 3000$ is chosen after visual examination that this time window is high above the typical relaxation dynamics. This initial value problem is integrated using a Runge-Kutta method of order 5(4) (Dormand and Prince, 1980).

Negative abundances are an artifact of the model but do not carry any ecological meaning. There are two main options to overcome this issue, depending of the type of dynamics one is interested in modeling. In somewhat *open* ecosystems, one could consider that any species that goes extinct locally ($x_i = 0$) can still arrive from a nearby site. In this context, one can add a constant small migration rate or a threshold below which a species never goes extinct (see section II.E and (Aguadé-Gorgorió and Kefi, 2024; Roy *et al.*, 2020) for a deeper discussion on migration).

Instead, here we are interested in modeling ideally *closed* environments such as microbial communities in the lab, where the community starts with a given set of species but, if a species goes extinct, it remains extinct and cannot reinvade unless the experiment allows for it (Chang *et al.*, 2023; Friedman *et al.*, 2017). To include this option, we define a very small threshold of $10^{-20}$, below which a species is assumed to go extinct and its growth rate and abundance is fixed to zero (see (Aguadé-Gorgorió *et al.*, 2025) for a detailed account on the role of imposing minimal abundance thresholds across community models).

Because we are interested in finding stable (or, at least, stationary) states, we measure if the abundance of each species remains the same after an additional time window of $t = 100$. To do so, we set a difference threshold of $10^{-3}$ as in (Aguadé-Gorgorió *et al.*, 2024b; Aguadé-Gorgorió and Kefi, 2024), so that if for at least one species $|x_i(t = 3100) - x_i(t = 3000)| > 10^{-3}$, we deem the state unstable and do not measure its properties. Instead, if all species abundances remain within this treshold, we define the state as stationary and record species abundances $x_i^*$ and their interactions $A^*$. To do so, we find the species that have zero abundance and erase them from the vector of stable abundances, $\boldsymbol{x}^*$, and erase their rows and columns from $A$. The output of a given simulation is then simplified to a vector of stationary abundances without zeros $\boldsymbol{x}^*$ and a matrix of interactions $A^*$, and most questions of this project center on studying the properties of these matrices of interactions $A^*$ for stationary states of coexisting species.

#### 2. Sampling and testing random species subsets

The above method can be costly if we need to repeat it a large amount of times, as we will do for example when measuring the statistics of intransitivity patterns (Section I.L). In this work we use our previous developments in (Aguadé-Gorgorió and Kefi, 2024) to implement a complementary method to find the vector of stationary abundances without zeros $\boldsymbol{x}^*$ and a matrix of interactions $A^*$ described above by taking advantage of the simplicity of the GLV model.

Simply put, given the same problem as above, defined by an interaction matrix $A \in \mathcal{N}(\mu, \sigma)$, we set a number of species $S^*$ randomly or in a stepwise manner between 3 and $S$, depending on the figure or method we want to implement (see next section). Communities with 2 species cannot show emergent coexistence and are well characterized by the pairwise model of Section I.B.1, so we are only interested on what can happen for triplets of species and above. We then select $S^*$ species randomly from the community, and obtain their interactions by eliminating all other rows and columns from $A$. This would account for assuming that all other species have gone extinct during the dynamics and these $S^*$ are the remaining ones, or else that, of the complete pool of initial species $S$, we have chosen to start with only a subcombination involving these specific $S^*$ species as done e.g. in (Friedman *et al.*, 2017), which is equivalent to setting some of the initial abundances in the numerical method above to zero ($x_i(t=0)$ for $i \notin S^*$) and integrating the dynamics from there.

We obtain the final abundance of the selected subset of species by implementing $\boldsymbol{x^*} = (I - A)^{-1}1$. If all row sums of the matrix of net effects are positive, it means that the subset of selected species all survive at positive abundance in the presence of the others. We compute the Jacobian matrix of this state as $J_{ij} = x_i^* A_{ij}^*$ for $i \neq j$ and $J_{ii} = -x_i^*$, obtain the eigenvalues using numpy's method `numpy.linalg.eigvals` and ask if all eigenvalues have negative real part to infer linear stability. If the conditions of positive abundances and linear stability are both met, we store the vector of stationary abundances of the surviving species only $\boldsymbol{x}^*$ and a matrix of their interactions $A^*$.

It is interesting to note here that the method of sampling subsets using mathematical results (this section) and the method of integrating the dynamics starting from all-positive initial conditions (section above) provide two complementary approaches; equivalent in mathematical terms but conceptually different. Starting from all species present and integrating the dynamics throughout time, during which some species can go extinct and a state with $S^* \lesssim S$ can be reached, is akin to the "random zoo" metaphor (Chang *et al.*, 2023; Serván *et al.*, 2018). Instead, selecting which subset of species we test from scratch is more similar to other type of experiments described below (Friedman *et al.*, 2017; Venturelli *et al.*, 2018), in which a handful of species –or different combinations involving only a subset of these– is grown together. Again, the two mechanisms become equivalent if, for the mechanism described in the section above, we do not start with all species present, but rather only a small subset of species being present and all other extinct. In this context both scenarios would be equivalent and would allow us to explore the properties of smaller communities with controllable size as done for Figure 3 in the main text.

As a final comment, it is worth noting that both approaches can be parallelized, as they are based on running a large amount of independent replicas of the same problem. This can be seen in the codes to generate for example figure 3 of the main text.

#### F. Evaluating emergent coexistence and the fraction of excluding pairs (Figure 2)

Once we reach a stable state with positive abundances encoded in $\boldsymbol{x}^*$ and pairwise interactions encoded in $A^*$, we ask if this state harbors emergent coexistence. To do so, we search for $A^*_{ij}$ elements that are smaller than -1, indicating that species $j$ in this community would exclude species $i$ if we grew them in co-culture, isolated from all other species of the community (see section I.B.1 above on pairwise exclusion). As a first approach, we only ask if $A^*_{ij}$ *or* $A^*_{ji}$ are smaller than -1: this means that a state is considered to harbor emergent coexistence if, for any pair of species $i \neq j$, $j$ excludes $i$, $i$ excludes $j$ or both (bistable competitive exclusion, see (Aguadé-Gorgorió *et al.*, 2024b) for a mathematical discussion and (Wright and Vetsigian, 2016) for an experimental analysis of bistable competition). We don't distinguish these three cases for now and are only interested if the species pair does not coexist in co-culture.

This highlights the EC property, whatever the specific outcome of one-way or bistable competitive exclusion. Once a state has been found to have at least one excluding pair and hence EC, we can plot figure 2A in the main text: we generate 100 random initial conditions and 100 species interaction matrices for each set of $\mu, \sigma$ values. For these 100 initial value problems, we count the number of states that are stable, $n$, and count how many of these states contains at least one excluding element, $n_{EC}$. In figure 2A we plot $n_{EC}/n$ for a large range of $\mu$ and $\sigma$ values. We find that EC is completely absent in a given domain, while it is present and in fact very common in a so-called *Emergent Coexistence Regime* (Fig. 2A main text), where on average about 80% of states carry at least one excluding pair (Fig. 2B in the main text). In section II.A of this Supplementary we discuss the mathematical developments to predict the boundaries of this EC domain.

In (Chang *et al.*, 2023), researchers not only measure if a state harbors or not excluding pairs, but also the fraction of these excluding interactions among the total species pairs in the community. We label this metric as the *fraction of excluding pairs*. Given the description above, it is rather straightforward to measure: a community with $S^*$ surviving species harbors $S^*(S^*-1)$ interactions and hence $S^*(S^*-1)/2$ species pairs. We count, out of these pairs, how many are excluding, meaning that, once again, $j$ excludes $i$, $i$ excludes $j$ or both.

In figure 2C of the main text, we simulate $10^5$ communities with random initial conditions and $\mu, \sigma$ again sampled randomly from uniform distributions within the domain of figure 2A. We measure the fraction of excluding pairs, for those communities that are stable and harbor at least one excluding pair. In communities without EC this value is of course zero and we do not plot it. In communities with $S^* = 3$, for example, we could find a Rock-Paper-Scissors triplet (3 pairs, 3 excluding pairs), two excluding pairs rescued by the other interactions, or one excluding pair and all other interactions non-excluding (see section I.B.2 above, for a discussion on EC in communities with 3 species). This results in a fraction of excluding pairs of 1, 0.66 and 0.33 respectively. The minimal fraction of excluding pairs is therefore $2/S^*(S^*-1)$ (a dark dashed line in figure 2C of the main text), and we discuss how to obtain the maximum fraction of excluding pairs in section II.A.3. The different experimental datasets are discussed in Section I.G and their values are incorporated after the simulations.

#### G. Gathering data on the fraction of excluding pairs (Figure 2C)

As described above, we are interested in finding empirical values for the presence of emergent coexistence and, more specifically, for the fraction of species pairs that coexist in a stable community but exclude one another in co-culture. We want to test our theoretical limit on the maximum number of exclusive pairs as a function of diversity in the random interaction assumption (see Section II.A.3 and figure 2C in the main text). Despite there is likely a lot of available data on estimated interaction matrices, from which one could count the presence of $A_{ij}$ elements smaller than the diagonal, we restrict our use of data to better-characterized empirical cases, where pairwise interactions are analysed from the outcomes of species co-cultures and not only estimated from time series data of the complete community.

Moreover, we also constrain the data to observations that consider certain degree of stability or stationarity of the species-rich community, meaning that species in a given community are found to coexist together (at least during a long transient). There exist, for example, very interesting datasets that sample species from a natural environment and measure the outcomes of their pairwise interactions (Higgins *et al.*, 2017), yet do not assemble such species to observe if they can coexist in a controlled environment. Even if these sort of observations are very relevant (pairwise co-cultures of species sampled from naturally co-occuring communities), we prefer to stick to well-controlled scenarios where the communities have been not only observed in their ecosystem of origin but also assembled and observed to coexist in the lab. Of course, these type of datasets are hard to obtain, as they require complex and multiple experiments involving many different species co-cultures, which is consistent with the attention gathered by works such as (Friedman *et al.*, 2017) and (Chang *et al.*, 2023). We believe that similar datasets will be obtained soon, and our work is open to confrontation with additional data: as discussed regarding the role of interaction structure, we can expect that particular datasets involving structured, far-from-random interactions might lead to different behaviors that what we have found (e.g. some structures could sustain more excluding pairs than what the random case proposes).

The work by (Chang *et al.*, 2023) sparked the inspiration behind our work by explicitly proposing that communities can hold many species pairs that do not coexist in isolation. Briefly put and recapitulating the explanation of the experimental setup in (Chang *et al.*, 2023), the team is able to assemble different communities in controlled environments with glucose as the sole limiting nutrient (Goldford *et al.*, 2018). Microbiomes from 12 soil and plant sources were suspended in M9 minimal medium, generating diverse bacterial pools with 110 to 1,290 exact sequence variants. The large species richness of these samples resembles the so-called *random zoo* model in theoretical ecology, where communities are thought to be assembled not by gradual invasions of one species at a time, but rather by starting from a large pool and allowing for species to become extinct during the dynamics (Serván *et al.*, 2018). The cultures underwent 12 sequential transfers, spanning approximately 84 bacterial generations. Community composition was tracked through 16S rRNA sequencing at various stages. Across all replicates, fewer than 25 coexisting ESV's were consistently detected, primarily from the Enterobacteriaceae and Pseudomonadaceae families (Chang *et al.*, 2023). Stationarity is measured by looking at the evolution of invasion fitness and estimating that ESV's present at the last passage would able to reinvade the community if starting at low frequency.

These 12 final communities contained between 5 and 13 ESV's, out of which researchers were able to isolate 62 species and grow them by pairs starting from different initial abundances. 144 pairs yielded a final conclusive outcome after co-culture growth, hereby explaining that, out of all species, not all pairs could be tested (in Figure 2C of the main text, we observe diversities ranging between 3 and 10, not 5 and 13, because only tested pairs are considered when building the tournament network). To measure for each of their assembled communities the fraction of excluding pairs, we used their figure 3A, in which the tournament outcomes across these 12 communities are presented, allowing us to count the number of species $S^*$ and number of excluding pairs, which we then divide by the total number of pairs $S^*(S^*-1)/2$ to obtain the fraction. Interestingly, researchers separate the outcomes of co-cultures into pairs in which one of the two species consistently goes extinct (competitive exclusion) and pairs in which the abundance of a species decreases, yet it is unclear if it would go extinct (on the path to competitive exclusion). We understand that one cannot truly discern if the dynamics of this later scenario would yield species extinctions and hence a clear exclusionary scenario, or else are simply a strong asymmetry in competition strength so that a species is simply kept at much lower abundance than the other but can still survive, which would imply a competition strength slightly above -1 (close to exclusion). To avoid potential bias, we have decided to not include them in the exclusionary counts and keep only the data of pairs that are observed to yield extinctions. If all observations of "towards the path of extinction" where considered as excluding pairs, one would find that some communities harbor more excluding pairs than what our approximation discussed in Section II.A.3 proposes, which is also possible in simulations: the analytical limit is only a statistical or aggregate approach, and for example some limit case communities fall above it (purple dot in Figure 2C). The combination of the data presented in Figure 3A in (Chang *et al.*, 2023) allows us to estimate first how many of the observed communities harbor at least one excluding pair: 66% of the communities harbor a clear example of exclusion, whereas 100% of communities harbor either a clear exclusion or at least a pair of species that are "towards the path of exclusion". From their figure 3A we can also infer the fractions of excluding pairs that can coexist in communities of 3, 4, 5, 7, 9 and 10 species respectively (Figure 2C of our main text): communities might hold more species, but their pairs are not sampled or results are not presented in Figure 3A and so we maintain only this values in place. Repeated values (communities with equivalent diversity and equivalent number of excluding pairs) were found, but are not marked in any particular way in figure 2C. The same work also takes advantage of the computed tournament matrix to measure the presence of low rank exclusions and rock-paper-scissors triplets, for which they observe low values and therefore highly transitive competition (see Sections I.L and II.D).

Some years before this work, (Friedman *et al.*, 2017) approached a similar problem and experimental exercise, even if the work proposed an apparently opposite conclusion to (Chang *et al.*, 2023), by which species composition in stable communities with a lower number of species could be predicted by pairwise outcomes. As we discuss below in Section II.B when discussing about predictability and the spectral radius, these two observations (Chang *et al.*, 2023; Friedman *et al.*, 2017) are in fact not contradictory: simple communities of 3 or 4 species can have low spectral radius and therefore composition can be predictable from pairwise outcomes (Friedman *et al.*, 2017). Larger communities with strong interactions are likely to have a larger spectral radius above 1, meaning that community composition will not correlate with pairwise observations and for example excluding pairs can coexist. In any case, researchers in (Friedman *et al.*, 2017) propose to obtain qualitative information from co-culture outcomes to predict community composition without the need to measure $A_{ij}$ elements.

To explore this, they used a set of 8 heterotrophic soil-dwelling bacterial species as their species pool. They grew different combinations of species in M9 minimal media for 5 growth-dilution cycles, spanning approximately 53 bacterial divisions. They measured the outcomes of the 28 pairs, 56 three-species combinations, a community starting with all the 8 species present, and each combination starting with 7 present and one absent. By looking at figure 2d of their main text, we can easily derive the tournament matrix of their species pairs. Using this tournament matrix, we study if there is evidence of EC in the triplets, meaning that a community with three coexisting species harbors some exclusionary pairwise interaction (Figure 3 of their main text), as well as the outcomes of larger communities involving 4 surviving species (Figure 5a of their main text).

The authors find that only 8.7% of trios build starting from 3 species harbor emergent coexistence (figure 3 of their main text). Yet, when they assemble communities starting from 8 or 7 species present, they find that 2 communities harbor EC, while one does not, leading to a 66% observation of EC albeit with very limited observations. The combination of these observations allows us to estimate the fractions of excluding pairs that can coexist in communities of 3 and 4 species, with trios observed to harbor either 1 or 2 exclusionary pairs, and the community with 4 species observed to harbor 1 exclusionary pair (Figure 2C of our main text). Repeated values (communities with equivalent diversity and equivalent number of excluding pairs) were found, but are not marked in any particular way in figure 2C.

In (Venturelli *et al.*, 2018), the authors study a similar problem of assembling synthetic communities, this time using bacterial strains from the human microbiome spanning the major phyla of human-associated intestinal bacteria. Their pool contains 12 species that they grow in monoculture and pairwise co-cultures (66 combinations), allowing the estimation of species growth and pairwise interactions. When compared to the experimental work described above (Chang *et al.*, 2023; Friedman *et al.*, 2017), the authors in (Venturelli *et al.*, 2018) do aim at measuring specific values to $A_{ij}$ elements from time-resolved pairwise co-cultures, allowing later to successfully confront their data to a GLV community model. Once again, pairwise interactions can result in both coexistence or competitive exclusion of one of the two species, making the dataset relevant in the context of our work confronting pairwise to community outcomes. As with the experiments described above, pairwise exclusion is again a very common scenario, involving about 50% of the studied systems. Visual observation of the data contained in Figure 2A of their main text allowed us to reconstruct a matrix of pairwise tournaments between species. Similar to (Friedman *et al.*, 2017), once pairwise interactions are well-characterized, the authors build a community starting with all 12 species present (8 in (Friedman *et al.*, 2017)) and 12 communities starting with all combinations starting with 11 species present and one absent (7 present in (Friedman *et al.*, 2017)). We therefore want to measure, out of all the species that can coexist in these communities, how many pairs did not coexist in the pairwise tournaments. Their dataset EV3 in their Supplementary Material contains the temporal dynamics of all 12+1 communities. In these, we look at species relative abundances at the the last temporal point, for which most species abundances are of the order $10^{-1}$ to $10^{-2}$, while a few have abundance to the order $10^{-4}$, which, consistently with the use of a survival threshold in our numerical simulations, are extinct and hence only a contamination or sampling error, or are in the path to extinction. The combination of their pairwise outcomes (Figure 2A in their main text) and the temporal data of the complete communities (Supplementary Dataset EV3) allows us to estimate the fractions of excluding pairs that can coexist in synthetic communities of 6, 7, 8, 9 and 10 surviving species respectively (Figure 2C of our main text, there are no communities in which all the initial 11 species survive). It is interesting to note that the fraction of excluding pairs in these communities appears to be much higher than for the rest of three experimental datasets that we analyzed. We do not have a strong hypothesis that explains these divergence, that could be related to intrinsic properties of the biotic interactions in each of the systems as well as how the experimental setup (different resource conditions or different initial abundances) modifies the structural properties of the communities.

Of course, our work does not claim that all systems are going to harbor EC signatures, and there can be many scenarios where coexistence is predictable from pairwise observations. We have investigated additional datasets that perform similar experiments confronting pairwise and community dynamics. For example, (Lele *et al.*, 2024) finds 3 out of 36 pairwise exclusions in a pool of species isolated from sourdough. Yet, when assembling multispecies communities with average diversities of 6 species, they did not find any of these 3 pairs coexisting in the full communities. Another work assembles a community with 5 species of soil microbes and finds that any species is able to invade any other subcombination of species, indicating complete coexistence (Castledine *et al.*, 2024), a result that is also found for protist communities of varied sizes with no evidence of pairwise exclusions (Pennekamp *et al.*, 2018). Some of the communities studied in (Friedman *et al.*, 2017) also lead to cases where the full community (often made up of 3 species) does not contain any pairwise exclusion, meaning that the number of excluding pairs is again null and leading to the scenario where pairwise observations predict community coexistence. We have decided to not paint neither simulated nor experimental datasets where the number of excluding pairs is zero in Figure 2C as these do not provide any additional information to the figure.

Finally, we have also added in Figure 2C of the main text a classical example of a rock-paper-scissors loop in E. coli (Kerr *et al.*, 2002) (blue square). In this well-known work, three strains of Escherichia coli are grown together and seen to coexist under certain local interaction conditions: the Colicin-Producing Strain (C) produces a toxin colicin that kills sensitive strains. The Sensitive Strain (S) is susceptible to the colicin toxin. The resistant Strain (R) is resistant to colicin but incurs a metabolic cost for this resistance. In this system, the colicin-producing strain (C) kills the sensitive strain (S), the sensitive strain (S) outgrows the resistant strain (R) due to the absence of the metabolic cost, and the resistant strain (R) outcompetes the colicin-producing strain (C) by avoiding the cost of producing colicin. This cyclic dominance maintains coexistence among the strains, leading to a community with three strains ($S^* = 3$). Even if the discussion could be opened regarding if these can be considered different species, they should considered as such in the GLV model as they have different interaction strengths. Because each of the three pairs of species has one excluding interaction, the fraction equals 1.

#### H. Measuring predictability properties of the community $(\phi, \kappa)$ (Figure 3)

In the present section we describe the numerical methods to measure $\phi$ and $\kappa$ from stable communities in our model. In the Sections II.B.3 and II.B.4 of the results we describe in more detail the meaning of $\phi$ and $\kappa$ and the mathematical developments to infer them analytically.

Once a stable community is found in our simulations, it is characterized by an interaction matrix $A^*$ (once again, this matrix contains zeros in the diagonal, as self-regulation is written as a separate identity matrix, see Section I.B.3 above). Following (Zelnik *et al.*, 2024), it is easy to measure the collectivity parameter that dictates the degree of collective integration of the community (Zelnik *et al.*, 2024), or, in mathematical terms, the spectral radius of $A^*$ that captures the relative weight of longer and longer chains of indirect effects. If $\phi$ is close to 1, we might need to look at very long indirect effects in order to say meaningful things about net effects and species abundances at equilibrium. If $\phi$ is larger than 1, direct interactions and chains of any given length no longer correlate with net effects and species abundances: the community is so collective that all species interactions ripple through the network and pairwise observations no longer contain any meaningful information of net effects. Mathematically, the spectral radius is the modulus of the largest eigenvalue of the matrix (not the real part, which tells us information of stability, but the modulus considering both real and imaginary parts, see (Zelnik *et al.*, 2024) for a clarification). In that sense, we simply obtain $\phi^*$, which is called $\phi$ throughout the paper, using the `numpy` library from python and `max(abs(numpy.linalg.eigvals(`$A^*$`)))`.

The condition number $\kappa((I - A^*))$ informs us about the error propagation of inverting a matrix, as discussed in detail in Section II.B.4 and e.g. (Demmel, 1987; El Ghaoui, 2002; Gilpin, 2024). Measuring the condition number of $(I - A^*)$ is equally easy in python. It only requires calling the built-in function `numpy.linalg.cond` or, as discussed in Section II.B.4, obtaining the largest and smallest singular values from `numpy.linalg.svd` and dividing them. In Section II.B.4 we discuss the condition number in detail as well as the mathematical developments to estimate it.

It is also interesting to remember the reader here how figure 3 is built. For the purple dots, we could find them by simulating the GLV model with $(\mu, \sigma)$ sampled in the domain of figure 2A and selecting those final stable states that carry EC as done for figure 2C. We could do the same by sampling random subsets and evaluating their feasibility and stability as described above. Yet, it is interesting to confront the spectral radius and condition number of stable communities with EC with those of stable communities without EC. Now, the first method, integrating the GLV model with all initial conditions positive, will lead to very large communities without EC, because these communities are found in the unique fixed point regime where interactions are weak (Fig. 2A and (Aguadé-Gorgorió and Kefi, 2024; Bunin, 2017)). Yet, to find smaller communities without EC and small $\phi$ or $k$, we need to assume that some species have gone extinct or where not present at start, even if interactions are weak, so that we do not end with most of the pool of species surviving together. This can be done by assuming a subset of initial conditions is zero, or by using the sampling method in which we sample random subsets of $A$ and test numerically for feasibility and stability. In figures 3 and 4 of the main text (and later in the figures measuring stability described in the section below) we use this last method for computational efficiency.

#### I. Designing empirical tests on predictability

Throughout the main text of this work we study how several metrics of a community and its interactions (namely $S^*, \phi, \kappa$) limit our capacity to predict community composition and hence explain emergent coexistence. To illustrate this, we want to design numerical simulations that replicate potential experimental scenarios where our work could apply in the context of designing and assembling microbial communities. Here we present the numerical construction to build the simplest possible tests. We refer the reader to very recent works such as (Arya *et al.*, 2025) or (Solé *et al.*, 2024) that describe in detail many empirical scenarios and practical considerations regarding the assembly of microbial consortia and how theoretical knowledge plays a role in those settings.

The collectivity parameter has been recently linked to multiple ecological properties of a community in relation to predictability, with a special emphasis on temporal unpredictability in the response to a perturbation (Kawatsu, 2024; Zelnik *et al.*, 2024). Inspired by the experimental exercises of e.g. (Chang *et al.*, 2023; Friedman *et al.*, 2017), here we want to understand how the collectivity parameter can inform about the composition of a community from information of pairwise interactions. To do so, we design the following test. We generate a pool of $S = 100$ interacting species, with interactions sampled randomly within the parameter range proposed in figure 2A of the main text. Again, interaction strengths more negative than -1 wil be indicative of exclusion, whereas the rest are coexisting interactions. Now, from this pool, we select subsets of $S^*$ species, with $S^*$ randomly chosen between 2 and 15, ensuring that all species pairs in this subset can coexist. The question is therefore wether these subsets, where all species coexist by pairs, will coexist in a larger community, and how this is related to $\phi$. We measure coexistence in the community setting by asking if all the row sums of $(I - A^*)^{-1}$ are positive, so that we have a relation between collectivity $\phi$ and the likelihood of community coexistence by assembling coexisting pairs. To transform the dataset that contains all values of $\phi$ for

$10^5$ sampled communities and a binary "coexists (1)" or "fails (0)" metric into a smooth function for probability, we define a `rolling mean` function in python. The rolling mean function computes a rolling (moving) mean of the dataset while ensuring data is sorted and down-sampled for smoother visualization or analysis. It first sorts the input arrays $\phi$ and "coexists" based on the values of $\phi$ to maintain an ordered sequence. Then, using a sliding window approach of fixed size (100 data points) and a step size of 1, it iterates through the sorted data, computing the mean of both $\phi$ and "coexists" within each window. This results in a reduced and smoothed representation of the original dataset, capturing underlying trends while minimizing noise.

The condition number $\kappa$ is explained in detail in Section II.B.4 once we introduce its mathematical formulation and results. Interestingly, the condition number of an ecological interaction matrix has also been recently used to discuss temporal predictability of a community of interacting species (Gilpin, 2024). Here we want to test how the condition number of a given matrix of interactions informs us about the potential error in predictability of species abundances and community composition. This can be encapsulated in the following simple test. We sample once again the same subsets of $S^*$ species and their interactions, $A^*$. We also consider a certain numerical error in our measurements, which can be introduced for example by setting $A^*_{ij} + \epsilon$, where epsilon is a Gaussian distributed number with zero mean and a standard deviation of e.g. $\sigma = 0.1$ (we will later see how, as expected, the predictability depends on the strength of this error, that we fix to $\sigma = 0.1$ for illustrative purposes). Now, we can ask: does the subset of species and our imprecise measurements of their interactions yield coexistence? This means, does the matrix $(I - A^* + \epsilon)^{-1}$ contain all-positive row sums? If so, our numerical approach predicts that the subset of species will coexist. We then ask if the real community, with abundances measured from $(I - A^*)^{-1}$, does coexist. If both communities yield positive abundances, our prediction was correct despite the error. If not, we are predicting a coexisting community in a setting where the real community will not coexist, and so we set this result as a failure (coexists=0). In parallel, we measure the condition number of $A^*$, which provides a numerical metric for how the $\epsilon$ error propagates when inverting the matrix. To visualize the results in a smooth way, we use the same rolling mean function as above, by which we define windows within the dataset and compute the mean $\kappa$ and "coexists" values within those windows. The results of both tests are discussed in Section II.B.5 and Figure 9.

#### J. Measuring May's threshold for stability in smaller systems

In the sections above, we have found stationary or stable communities by using previous methodology (Aguadé-Gorgorió *et al.*, 2024b; Aguadé-Gorgorió and Kefi, 2024) to integrate the dynamics or sample stable subsets from the pool. Later in the work, we find it interesting to understand what aggregate properties of interactions allow some communities to be stable. In this regard, the seminal work by Robert May (May, 1972) provided a first understanding that there is a limit in the statistics of a random matrix for it to maintain negative eigenvalues and hence linear stability. In the GLV model, this criteria translates to the condition by which the interaction strength of a fully-connected community must be maintained within this lower bound and cannot become lower (more competitive) (Aguadé-Gorgorió and Kefi, 2024; Bunin, 2017; Mallmin *et al.*, 2024):

$$\mu^* > \min(\mu^*) = \sqrt{\frac{S^*}{2}}\sigma^* - 1, \tag{18}$$

which is valid for large $S^*$, where $S^* \geq 50$ has been found to be a good bound (Aguadé-Gorgorió and Kefi, 2024). For smaller diversity $S^*$, a correction proposed in (Aguadé-Gorgorió and Kefi, 2024) using numerical methods estimates the threshold to be found at

$$\mu^* > \min(\mu^*) \approx \frac{(S^*)^{1.08}}{14.118}\sigma^* - 1. \tag{19}$$

However, both of these methods are statistical boundaries. In Section II.C we will show that small communities can overcome this boundaries by plotting the distance to the critical point using absolute values, $d = |\min(\mu^*)| - |\mu^*|$. A positive value means that $\min(\mu^*)$ is more negative than $\mu^*$ (stronger competition in absolute terms) and so the state is within the stable domain predicted by the random matrix theory of the GLV model. Instead, a negative distance means that the state has stronger interaction strength than what the boundary predicts. We find that, at low to moderate diversity, there are many states that harbor stronger competition than what the Random Matrix Theory boundary would predict (see Figure 10 and Section II.B for a detailed discussion).

#### K. The Routh-Hurwitz criteria

We then ask if there are other explanations that help us understand ecologically how can states with $S^* \ll 50$ maintain stability under strong competition. In this context the Routh-Hurwitz (RH) criteria provide a valuable tool (Bodson, 2020; Routh, 1877; Toni, 2014). Very generally, the RH criteria in the context of the stability of a Jacobian matrix are met when positive long loops do not overcome the strength of negative shorter ones (Levins, 1974). Yet, this is an extreme simplification of the RH criteria, which get much more complex than that for species-rich systems. The purpose of this section is not to provide a detailed mathematical explanation of the RH criteria, but to provide a simplified rationale that allows to bring forward some intuitions and is consistent with previous work in ecology (Dambacher *et al.*, 2003; Levins, 1974; Neutel *et al.*, 2002). Here we focus on the approach of the RH criteria that studies the so-called Hurwitz determinants (Hurwitz, 1895) rather than the Routh test (Horn and Johnson, 1994).

Again, we know that the linear stability of a system is captured by the sign of the eigenvalues of its Jacobian evaluated at equilibrium. This captures if, under a small perturbation, the system will return to equilibrium or not. The elements of the Jacobian matrix $J$ are the direct effect between the abundance of a species and the growth of another evaluated at the equilibrium $x^*$, namely

$$J_{ij} = \frac{\partial}{\partial x_j}\left(\frac{dx_i}{dt}\right)_{x_i = x_i^*}. \tag{20}$$

In the specific case of the GLV model, the Jacobian is directly proportional to abundances and interaction strengths, and has off-diagonal elements $J_{ij} = x_i^* A_{ij}$. Thus, it contains interactions strengths rescaled by species abundances. The rationale of the RH criteria might not be constrained to the GLV model, as all models can be linearized around an equilibrium. The basic problem is to find the conditions by which the eigenvalues of $J$ have negative real part, without a priori knowing anything of $J$. The eigenvalues of $J$ are the solutions of the equation

$$\det(J - \lambda I) = 0. \tag{21}$$

In the case of $S^*$ species (dimensions), this can be written as the polynomial equation

$$(-1)^{S^*}\lambda^{S^*} + C_1\lambda^{S^*-1} + C_2\lambda^{S^*-2} + ... + C_{S^*} = 0. \tag{22}$$

The Routh-Hurwitz (RH) criteria relate the sign of all eigenvalues $\lambda_i$ to properties of the coefficients $C_k$ of the characteristic polynomial. An essential property is that all elements $C_k$ are composed by non-trivial closed loops of different length. In (Levins, 1974) and subsequent work (Dambacher *et al.*, 2003; Neutel *et al.*, 2002), these coefficients have been called "Feedbacks". In particular, $C_k$ was labeled as $F_k$ and defined as *the feedback of a matrix at level k*, and was later directly linked to ecological feedbacks between $k$ species (Neutel *et al.*, 2002). We believe this might be a misleading name, because $C_k$ contains nontrivial combinations of loops of different length and can easily be much more complicated than what one could understand are "the loops of length $k$". For example, in a 3-species GLV model, the three coefficients of the characteristic polynomial are:

$$C_1 = J_{11} + J_{22} + J_{33} \tag{23}$$

$$C_2 = J_{12}J_{21} + J_{13}J_{31} + J_{23}J_{32} - J_{11}J_{22} - J_{11}J_{33} - J_{22}J_{33} \tag{24}$$

$$C_3 = J_{12}J_{23}J_{31} + J_{13}J_{32}J_{21} + J_{11}J_{22}J_{33} - J_{11}J_{23}J_{32} - J_{22}J_{13}J_{31} - J_{33}J_{12}J_{21} \tag{25}$$

Already looking at this coefficients it seems clear that they are not only loops of a given length, as the elements of $A^2$ can take the form of $A_{ij}A_{jk}$ and be considered chains of length two. Instead, they appear to be combinations of products of closed loops of different lengths. What is clear is that the coefficients all involve *closed* loops, making this different to the elements of $A^k$ that appear in Neumann series which involve all chains of interactions, open and closed. To find a useful and simplified way to capture the content of these coefficients without loosing detail, (Levins, 1974) defined the contents of $C_k$ as

$$C_k = \sum(-1)^{m+1}L(m,k) \tag{26}$$

Each $L(m,k)$ is a product of $k$ elements that involves $m$ different loops. We can relate these $L(m,k)$ to the examples above. For example, $J_{12}J_{23}J_{31}$ has $k=3$ elements, and only $m=1$ closed loop. $J_{11}J_{23}J_{32}$ also has $k=3$ elements, but contains $m=2$ closed loops. This is why $L(1,3) = J_{12}J_{23}J_{31}$ has a positive sign in $C_3$, as $(-1)^{1+1} = +1$, while $L(2,3) = J_{11}J_{23}J_{32}$ comes with $(-1)^{2+1} = -1$.

From here we could start to make careful connections with ecological feedbacks. Suppose $J_{11}$ and $J_{23}J_{32}$ are two negative feedback loops. The first through self-regulation, as all diagonal elements of the GLV Jacobian will be negative ($J_{11} < 0$), the second through predation ($\text{sign}J_{23} = -\text{sign}J_{32}$). A system containing only these two loops, therefore, should be linked to stabilization. If we multiply these two loops, we do not want a positive sign to emerge: we expect the combination of this two loops to remain negative and hence be linkable to stability. We will later see the mathematical explanation of this.

So far, we have come to this: Each coefficient of the characteristic polynomial is a sum of terms involving feedback loops. More specifically, the terms of the coefficient $C_k$ are all products of $k$ elements of the Jacobian that are made up of loops. For example, if $k=3$, $J_{12}J_{23}J_{31}$ and $J_{11}J_{23}J_{32}$ are products with 3 elements containing loops, while $J_{11}J_{21}J_{13}$ will not appear in $C_3$ because it involves non-closed loops. Each of these loop combinations will have positive sign if they contain an odd number of loops, or negative sign if they have an even number of loops.

The RH criteria impose a sign for the non-trivial combinations of different $C_k$ elements that is related to a set of determinants. These so-called Hurwitz determinants are highly nontrivial and it is beyond the scope of this supplementary to explain this in detail. We refer the reader to very good works (Bodson, 2020; Clark, 1992; Toni, 2014) where this has been explained with solid mathematical rigour.

The first of the RH determinants are:

$$\lambda_1 = C_1 = \sum_j J_{jj} < 0 \quad \text{(negative trace, self-regulation)} \tag{27}$$

$$\lambda_2 = \begin{vmatrix} C_1 & C_3 \\ C_0 & C_2 \end{vmatrix} = C_1C_2 - C_0C_3 = C_1C_2 - (-1)^N C_3 > 0 \rightarrow C_1C_2 > -C_3 \tag{28}$$

$$\lambda_3 = \begin{vmatrix} C_1 & C_3 & C_5 \\ C_0 & C_2 & C_4 \\ 0 & C_1 & C_3 \end{vmatrix} = C_1C_2C_3 + C_3^2 - C_1C_5 - C_1^2C_4 < 0 \tag{29}$$

...

The second determinant for example is related with measuring the weight of loops of length 3 compared to length 1 and 2, as done for empirical food web data in (Koch *et al.*, 2023; Neutel *et al.*, 2002, 2007; Neutel and Thorne, 2014). However, as systems with diversity larger than 3 are considered, coefficients like $C_4$ and $C_5$ become more and more intrincate, involving all combinations of closed loops of length 5, 4, 3, 2 and self-loops. This is where the RH criteria are clearly beyond the "positive loops are destabilizing", which is only valid for small species motifs. In the multispecies RH criteria, the loop combinations leading to stable or unstable behavior become extremely complex.

Overall and very broadly, the RH criteria seem to tell us that stability is not simply mediated by $\mu$ and $\sigma$ as for large random matrices: when the elements of a matrix are random, but they are not positioned randomly across the matrix, the stability properties can change. Certain topologies could increase stability by disrupting long feedback loops, and certain interaction strength distributions could increase stability by ensuring that long loops are filled with weaker elements than expected at random (Neutel *et al.*, 2002).

We discuss in Section II.C and plot in Figure 10 the sign of the second determinant for communities of different size and again with (purple) and without (gray) EC. To measure the second determinant, we compute the Jacobian matrix of a system with abundance $x^*$ and interactions $A^*$ as discussed in Section I.B and make use of numpy's built-in `numpy.poly` function, that returns the characteristic polynomial of the Jacobian matrix. We compute the second determinant by calculating $C_1C_2 + C_3$. We also observed equivalent results for stability for the next Hurwitz determinant, $C_1C_2 + C_3 + (C_3)^2 - C_1C_5 - (C_1)^2C_4$ (not shown).

As discussed in Section II.C, we will see that all communities fulfill the RH criteria, at least for the first determinants. This should not come as a surprise, as these are known necessary conditions for the system to be stable. Yet, taken together, our results seem to indicate that small communities can sustain stronger competition than what the random matrix theory (Random Matrix Theory) approximation predicts by favouring specific interaction strength configurations that fulfill the RH criteria. As we later discuss, this does not mean that Random Matrix Theory predictions are not valid, but rather that they do not really operate for systems involving only a handful of species, where more nuanced feedback structures can play a dominant role. Despite claiming that *negative loops are stabilizing* might be an oversimplifcation of the RH criteria, at least for the second determinant we can gain some intuition that tells us that *non-trivial combinations of short, negative feedbacks need to counterbalance long positive feedbacks.* More importantly, these feedbacks are feedbacks in the Jacobian matrix, which is not exactly the matrix of interactions as it also contains the weights imposed by the abundances of species, themselves related to the inverse matrix of net effects. We consider that a mathematical analysis of the RH criteria and their implications for ecological stability complementing existing work (Dambacher *et al.*, 2003; Levins, 1974; Neutel *et al.*, 2002) with more recent developments would provide a valuable tool to network ecology.

#### L. Competitive hierarchies and intransitivity (Figure 4)

Competitive intransitivity, in very broad terms, refers to situations of multispecies competition in which one cannot define a clear hierarchy of best-to-worst competing species (Allesina and Levine, 2011; Gallien *et al.*, 2017; Laird and Schamp, 2006; Levine *et al.*, 2017; Soliveres *et al.*, 2015). The central importance of intransitivity stems from the early concept of competitive exclusion, itself tied to the survival of the fittest (Hardin, 1960). The history of competitive interactions as a central force in nature –opposed to the presence of other, often positive interaction types– could fill many pages, and we fully agree that competition is not the only ingredient at play and ecology even if it has centered most of our attention for decades (Bimler *et al.*, 2025; Kéfi *et al.*, 2024). In any case, if one assumed that competitive exclusion is abundant, as observed in EC experiments (Chang *et al.*, 2023), and that a system is defined by a perfect hierarchy of best, second-best, third-best competitor and so on, it is easy to arrive to the conclusion that such a system cannot hold coexistence, and that the best competitor will at the end win. This reasoning sparked the search for competitive structures that dismantled single-species dominance.

There exist many ways in which one could measure the degree of intransitivity of a given ecological community (Gallien *et al.*, 2017; Koch *et al.*, 2023), and we do not aim to review them here or apply them extensively in our model (see e.g. (Feng *et al.*, 2020) for a comparison of methods). Briefly put, we could separate metrics of intransitivity in those that use the binary outcomes of competition (pairwise coexistence or exclusion, a *tournament* matrix (Allesina and Levine, 2011)), from those that require knowledge of specific interaction strength values $A_{ij}$. For the first, it suffices to obtain a tournament network, in which we know whether species $A$ excludes species $B$, they coexist, or $B$ excludes $A$, for all species pairs, as done for example in figure 3A of (Chang *et al.*, 2023). This, of course, is particularly useful in the light of our work: available data of pairwise tournaments does not necessarily contain $A_{ij}$ estimates, pairwise outcomes alone are much easier to obtain and, as discussed throughout the work, are more robust (in fact, much less dependent to) to numerical error. Following (Chang *et al.*, 2023) and (Higgins *et al.*, 2017), we choose in the main text to measure the fraction of Low Rank Exclusions and the fraction of Rock-Paper-Scissors triplets as two key metrics of intransitivity that are independent to specific $A_{ij}$ values, and study an additional metric below (Laird and

Schamp, 2006)

#### 1. Competitive ranks and Low Rank Exclusion

One way of measuring the presence of intransitivity is the following. Assume we can order all species by a certain *rank*, a number that captures their competitive capacity or position in a hierarchy. Are there species with lower rank excluding species with a higher rank, or are all exclusions pointing "downwards" in this hierarchy? One framework to define a competitive rank, discussed for example in (Chang *et al.*, 2023; Feng *et al.*, 2020; Gallien *et al.*, 2017) is to define the competitive rank $R$ of a species $i$ as

$$R_i = \frac{\text{Number of wins - Number of losses}}{\text{Total number of interactions}}. \tag{30}$$

In this simple definition, a species that tends to win more often has a positive rank close to 1, a species that tends to be excluded more often has a negative rank close to $-1$, and species that fall in-between these two extremes or whose interactions are mostly coexistence (weak competition or cooperation) will have a rank close to zero. By looking at the tournament matrix (exclusions for elements smaller than -1, coexistence for the rest), we can easily attribute a rank to each species in a community.

Next, in a given community, we count the total number of excluding interactions, as done before to count the number of excluding pairs (the number of $A_{ij} < -1$ elements, see section I.B). However, we propose a correction to avoid counting inconclusive tournaments: cases of bistable exclusion, where both $A_{ij}$ and $A_{ji}$ are smaller than -1, are not counted. This is because, in these cases, different initial conditions would result in different winners (Aguadé-Gorgorió *et al.*, 2024b), so that we could not conclude the direction of the exclusionary effect, as empirically observed in (Wright and Vetsigian, 2016). Nevertheless, the fraction of this interactions in the random model and in the subsequent stable communities that emerge from it is much smaller than the fraction of one-way exclusions (not shown). We do not expect it to alter the qualitative results significantly, leaving this study for future work. Of the one-way excluding interactions, we count the fraction in which the excluder species has a lower rank than the excluding species. Importantly, this means that we are not measuring out of the total number of interactions ($pS^*(S^* - 1)$) how many are a low rank exclusion. We are only measuring out of the total number of *excluding* and *one-way* interactions, how many are a low rank exclusion. The first metric would give very low values, as many communities harbor a lot of interactions that are not exclusionary. In this work we show that even the second metric (the correct one) gives very low intransitive values, consistent with empirical observations of laboratory microbial communities (Chang *et al.*, 2023; Higgins *et al.*, 2017). Moreover, we do not count bistable exclusions nor low-rank bistable exclusions, meaning that they are erased from the numerator and denominator of the Low Rank exclusion division and we do not believe they would modify the result, together with the fact that they represent only a small fraction of all exclusions as discussed above.

As discussed in detail in section II.D.1, this method has a caveat: there is a minimum number of exclusions that are necessary for a Low Rank Exclusion to happen: enough exclusions for a species to have high rank, and for another to, having a lower rank, still have at least one *win* in its list (excluding the higher ranked species). In section II.D.1 we evaluate and correct this caveat, and discuss the results of the main text, that show that low rank exclusions are in fact very rare even in the random unstructured case.

#### 2. Competitive triplets and Rock-Paper-Scissors

A fundamental type of intransitive motifs concern Rock-Paper-Scissors (RPS) triplets (Allesina and Levine, 2011; Kerr *et al.*, 2002; Sinervo and Lively, 1996). Their importance can be understood because they provide the simplest form of competitive coexistence in a community of three excluding competitors. As discussed in the main text, three species can coexist even wehn all pairs result in competitive exclusion, provided that the structure of exclusions is RPS: $A$ excludes $B$, $B$ excludes $C$ and $C$ excludes $A$ (Allesina and Levine, 2011; Gilpin, 1975; Levine *et al.*, 2017). This crucial and easy-to-understand example provides a first picture of how intransitive competition can allow for species coexistence beyond two-species systems. Yet, it could well be that the RPS motif, which is a particular but intuitive three-species example, has permeated our understanding of indirect effects towards thinking that multispecies coexistence *requires* closed RPS loops (Levine *et al.*, 2017). Yet, the presence and abundance of these structures in species-rich models and in natural ecosystems beyond the three-species case study remains an open question.

To measure the presence of RPS triplets, a first point is of course to define a competitive triplet. Species-rich communities harbor many triplets of interacting species: the point is to focus on those triplets where the three species are connected by competitive exclusion. As in (Chang *et al.*, 2023; Higgins *et al.*, 2017) we count, given a matrix $A^*$ of species interactions resulting in a stable community, the number of different instances in which a trio of species is connected by competitive exclusion. This implies $A_{ij}$ or $A_{ji} < -1$, and the same for a $jk$ and a $ki$ pair. Then, we count the cases in which such a triplet has a RPS motif, in which there is no specific rank of best-to-worst species, but rather each species excludes one species and is excluded by the other.

It is interesting to notice, as discussed in the main text and later in section II.D.2, what is a null expectation for the fraction of triplets that are RPS. First, we find from simulations that competitive triplets can only yield stable coexistence if exclusions are one way, so that $A_{ij} > 1$ *and* $A_{ji} < -1$ or vice-versa for all pairs (not shown). Now, we know that the competition matrix for a stable competitive triplet has 3 exclusionary elements and 3 non-exclusionary elements. We can order these elements in two RPS motifs (clockwise and counter-clockwise orders if we were to draw them) and 6 non-RPS motifs (species 1 wins over 2 and 3, 2 wins over three; species 1 wins over 2 and 3, 3 wins over 2; and so on...). The null expectation, therefore, would imply that every 2 out of 8 triplets are RPS, a 25%. Any community where a much larger fraction of triplets are RPS might imply that these overabundant RPS motifs are required for stable coexistence. In section II.D we discuss in detail the results shown in the main text.

3. Other metrics of intransitivity

In a seminal work with rather straightforward title (*Competitive Intransitivity Promotes Species Coexistence*), Laird and Schamp studied the outcome of tournament interactions in a spatially embedded (cellular automaton) community (Laird and Schamp, 2006). Their work brought forward very relevant discussions and a clear goal of modeling species-rich systems. As opposed to our work and many ecological modeling efforts where interactions are weighted, their interaction matrices are explicit binary tournaments, with a 1 implying exclusion and a 0 implying coexistence. Because of this configuration, an interaction between two neighboring individuals would only result in coexistence with no effect, or extinction of one of the two species. Following our work and the clear expression of indirect effects as $A_{ij}A_{jk}$... elements, it is clear that pure (binary) tournaments might not be able to produce interesting outcomes in terms of collective behavior, as all coexisting interactions are considered null instead of simply weak ($A_{ij} = 0$) and they just lead to no effects at all, whereas all exclusionary effects have the same specific strength ($A_{ij} = 1$) and cannot yield heterogeneous competition.

Nevertheless, the authors provide multiple interesting metrics of intransitivity in competition tournaments, which are relevant to us as we look at datasets for which $A_{ij}$ is not always or not necessarily measured. The central metric that the authors propose looks at the distribution of *wins* across species. In our model, this accounts for counting the number of elements smaller than -1 that one can find for each column of the interaction matrix $A^*$ of a surviving community. The authors then measure the standard deviation of this vector of wins, which provides information on how evenly is competition distributed: a purely homogeneous vector of wins would mean there is no signature of a clear competitive dominance. To obtain their metric, the authors also measure the standard deviation of the most homogeneous vector containing the same number of wins as well as the most heterogeneous vector containing the same number of wins. We constructed two reference vectors: a homogeneous and a heterogeneous distribution of wins. The homogeneous vector evenly distributes the total count of wins in the original system across all positions. Any remainder is distributed incrementally to ensure an exact sum. The heterogeneous vector follows a decreasing sequence, assigning higher values to the first positions and tapering down to zero. This sequence is then scaled to match the total count, with minor adjustments for rounding errors. These distributions provide contrasting benchmarks for evaluating the observed competition hierarchy. In (Laird and Schamp, 2006), the authors propose a metric of relative transitivity: $\sigma_{obs} - \sigma_{min}/\sigma_{max} - \sigma_{min}$, which approaches 0 in highly intransitive systems without hierarchy, and 1 in highly transitive systems with a clear hierarchy. This metric has a caveat in our model where there might be many systems with a small number of wins: in communities with only one exclusionary pair, all vectors would be the same (1 species has 1 win, all other species have zero wins). To avoid this issue but maintaining the notion of the original work, we also measure transitivity as $\sigma_{obs}/\sigma_{max}$. This measure does not have a bottom, basal expectation (we would not know if any value much lower than $\sigma_{max}$ implies more or less intransitivity). A value of 1, however, indicates that the observed vector of wins is as hierarchical as it can be. In Section II.D.3 we describe the results observed for the two metrics, which are consistent with the observations of the main text by which competition tournaments in our system are not necessarily intransitive once many species interact.

There is a large body of interesting theoretical work analysing other metrics of intransitivity (see e.g. (Feng *et al.*, 2020)) and the potential role it can play in ecological dynamics (see e.g. (Allesina and Levine, 2011; Gallien *et al.*, 2017)). It is not at the core of our work to analyse all these details: we do not claim that transitive or intransitive interactions favour or impair species coexistence. We only claim that species coexistence of strong competing pairs (emergent coexistence) is possible even under highly transitive systems provided that many species interact. This, at the very least, seems to imply that strong forms of intransitivity such as rock-paper-scissors loops or low rank exclusions are not *required* for competitive coexistence. We have also left aside many interesting metrics of intransitivity that consider not only tournament information, but specific species interaction strengths (Feng *et al.*, 2020). Because some of the most valuable data used throughout our work provides information of co-culture outcomes but not measurements of interaction strengths, we would not be able to test additional $A_{ij}$-specific metrics in such datasets.

### 4. Building Figure 4

The sections above propose a simple method to measure the fraction of exclusions that are low rank exclusions and the fraction of triplets that are RPS triplets in a stable community, given its matrix of species interactions $A^*$. Yet, how can we obtain statistics of these fractions across communities?

To build panels A and C of figure 4 in the main text taking this into account, we first obtain a very large number of stable states from the species pool using the methods described above and $10^7$ random initial conditions and interaction statistics $\mu, \sigma$ in the domain of figure 2A. For each stable state that we find to harbor EC, we record its fraction of RPS triplets and low rank exclusions as described above, as well as the number of species that survive in this stable state, $S^*$. In panels A and C we plot, for each value of species diversity $S^*$, the mean and $\pm 0.25\sigma$ of the standard deviation of all RPS and LRE fractions. Because there are in general few states with many species that contain a considerable number of exclusions (see Figure 2C in the main text, few states close to the maximum fraction in a red dashed line), we require a lot of simulations to make these statistics informative at high-enough species diversity. Also, to make the visualization easier and less confounding, we choose to paint only 0.25 standard deviations instead of $\pm 1\sigma$ so that each shaded area is distant enough. Even when painting $\pm 0.25\sigma$ we can observe a large variation of intransitivity metrics except for the $S^* = 3$, again pointing towards a high randomness and lack of selection of intransitive motifs.

To build panels B and D of figure 4 in the main text, we aim at conveying the probability distribution of the RPS and LRE fractions we would expect for a given diversity value. These probability distributions, in turn, highlight a basal expectation for the experimental tests proposed in (Chang *et al.*, 2023; Higgins *et al.*, 2017): If we observe many competitive triplets and exclusions, what are the fractions of RPS and LRE motifs that we would expect? A fundamental issue here is that many communities might harbor, for example, a single competitive triplet, or maybe two, or a small number of overall exclusions (Fig. 2C of the main text. The statistical distribution of a triplet being RPS, therefore, appears almost binary: if we painted these probability distributions across communities, because each community harbors few triplets, the average distribution would in turn look binary. Yet, we are interested in the overall expected probability of a RPS motif in the case of seeing many triplets: for example, in both (Chang *et al.*, 2023) and (Higgins *et al.*, 2017), the authors count how many triplets are RPS out of a large number of competitive triplets. Averaging the RPS and LRE fractions across communities is therefore not a solid statistical approach.

To overcome this issue, we propose exploring the statistics of RPS and LRE not for a given community, but for a large observation of many communities. Once we have the list of all communities, their given $S^* = 8$ value, and the RPS and LRE fractions for each community, we define a single experimental observation as the mean RPS and LRE fractions found across a random subset of 100 communities. Another option would be e.g. to directly recapitulate (Chang *et al.*, 2023) by asking for the mean fraction of RPS triplets found after observing 77 triplets, which is the data provided in their work (0/77). We generate a list of 100 experiments, in which each experiment is in turn the mean RPS and LRE fraction of 100 randomly selected communities. This coarse-grains RPS and LRE measures across many communities into a single pair of values. We then plot the estimated probability distribution for RPS and LRE fractions in these 100 sets of 100 communities each. To do so, we employ a kernel density estimate (KDE), a non-parametric method for estimating the probability density function of a continuous variable (Chen, 2017). KDE works by placing a smooth kernel (typically a Gaussian) at each data point and summing these kernels to produce a continuous density curve. This provides an intuitive representation of the underlying distribution (the probabilities of a triplet being RPS or an exclusion being LRE) without assuming a specific parametric form. We use the `kdeplot` function from the `Seaborn` library, a Python data visualization tool built on Matplotlib.

#### M. Nested interactions, single-resource competition and cross-feeding

The assumption that interactions are random and sampled from a normal distribution provides a null model or a starting point to describe a system for which we do not have any information of the possible interaction structure. Once again, we do not claim that interactions in realistic settings are fully random, but rather that the random model provides a basal expectation over which to test the presence of patterns or the role played by additional structures. For example, we refer the reader to recent works where random interactions at the species-level are coupled to more general non-random structures at the functional group level (Barbier *et al.*, 2018; Giral Martínez *et al.*, 2024; Martínez *et al.*, 2024)

When investigating the role of intransitive structures as described above, it becomes interesting to test additional structures in the network of interactions that can modify intransitivity. A limit-case scenario would be for example that of a purely triangular matrix, where the lower triangle is filled with zeros so that $A_{ij} = 0$ if $i > j$. In such a configuration, species 1 perceives interactions by all other species, species 2 by all minus the first, and so on, until species $S$ does not perceive any effect and only undergoes self-regulation. Albeit there is strong evidence that certain degree of nestedness –a hierarchy from generalist to specialist species– exists for example in plant-pollinator communities dominated by mutualism (Bascompte, 2009), assuming that all interactions involving facilitation $(+,+)$ but also predation $(+,-)$ or competition $(-,-)$ are perfectly nested is of course an extreme assumption. However, the fully nested matrix provides an interesting way to disentangle emergent coexistence and indirect effects from intransitive loops. As discussed in the results section, a nested matrix cannot have closed loops such as a RPS architecture nor any kind of low-rank exclusion. However, indirect effects are still possible and higher-ranked species can rescue lower-ranked species by keeping intermediate competitors at bay. To build such a matrix, we simply generate interaction matrices in the same way as above, with all off-diagonal elements sampled from a Gaussian distribution with $\mu, \sigma$ parameters. We then impose zeros in the lower triangle by setting $A_{ij} = 0$ for all $i > j$. Of course, the new matrix will not have the same mean and standard deviation as the original one, and because of that it would not be valid to compare quantitative properties (e.g. the fraction of excluding pairs) between these matrices. However, we can still measure intransitivity properties in this matrix and ask whether emergent coexistence is possible in the absence of e.g. RPS loops (Section I.L).

Could it be that slightly nested architectures exist due to ecological properties of a given system, even if not in the form of a perfectly triangular setting? In this context it is interesting to reread (Chang *et al.*, 2023) and previous and very relevant work by the same team (Estrela *et al.*, 2022; Goldford *et al.*, 2018). In particular, the authors propose that their experimental setup leads to a specific interaction setting, in which species are grown on a single limiting resource and can also grow due to cross-feeding of necessary metabolites. Here, we propose a modification of the fully random matrix that tries to account for these particularities of the experimental datasets under study but in the most general way possible, and ask whether the new, modified matrix predicts better the patterns of the model, mostly in terms of the absolute absence of intransitive RPS motifs that is found in (Chang *et al.*, 2023). If one remembers the reasoning behind the random matrix approach, it was proposed that randomness and the lack of internal structure nor details emerges from scenarios where interactions are mediated by many different traits (we refer the reader to (Barbier *et al.*, 2021) for a very clear discussion of this). If species are competing for different resources, competing for space in different ways, etc. (that is, if interactions are determined by a multiplicity of traits), it is expectable that interactions lack somehow a structure and become heterogeneous.

Instead, what happens if competition is mostly mediated by a single trait, which is the capacity to absorb, uptake or consume a single limiting resource as in (Chang *et al.*, 2023)? In this context, it is reasonable to assume that each species has a given capacity to interact with that resource. In the simplest terms, one can consider that this capacity is mediated by a single trait, $\gamma_i$, that captures the speed at which the species consumes the resource and hence makes it unavailable to a second species (Lee *et al.*, 2023). Of course, one could consider cases in which the rate of consumption is different from, and even imposes a trade-off to, the efficiency in transforming that resource to energy (Vincent *et al.*, 1996). This could in fact explain certain degree of heterogeneity in the effective interaction matrix that the GLV model cannot capture: in consumer-resource models, an interaction between two species $A_{ij}$ is in fact a phenomenological description of a more nuanced process related to the indirect competition for a shared resource. Nevertheless, for a first assumption that is consistent with the rest of our work, we consider that we can consider the resource layer implicitly and focus on direct pairwise competitive interactions. We refer the reader to (van den Berg *et al.*, 2022) for a great discussion linking the phenomenological approach of GLV-like models with the more mechanistic approach of consumer-resource models.

Now, if $\gamma_i$ is the resource consumption of species $i$, we could for example assume that competition between species $i$ and $j$ is linearly proportional to the difference in consumption rates. Alternatively, we could for example assume that, in a competition between two species, each species perceives a competition that is proportional to how much better the other species is at uptaking the resource. Once again, because our main focus is on gaining a qualitative understanding on properties such as emergent coexistence, indirect effects or intransitive architectures, it is out of the scope of this work to analyze the quantitative differences that emerge from using variations of this single-trait competition. We refer the reader to e.g. (van den Berg *et al.*, 2022; Chesson, 1990; Laigle *et al.*, 2018; Vincent *et al.*, 1996) for more in-depth discussions on resource competition and species traits that are beyond the scope of the present work.

One way of writing the interaction mediated by the single resource consumption trait is then

$$\tilde{A}_{ij} = -\frac{2\gamma_j}{\gamma_i + \gamma_j}. \tag{31}$$

In this setting, both species $i$ and $j$ perceive competition in the presence of the other. If species have the same or very similar trait $\gamma$, $\tilde{A}_{ij} \approx -1$ and competition is equivalent to self-regulation, hereby accounting for a neutral scenario. Instead, if species $j$ is a much better consumer than species $i$ ($\gamma_j \gg \gamma_i$), species $i$ perceives $\tilde{A}_{ij} \approx -2$, whereas species $j$ perceives $\tilde{A}_{ji} \approx 0$. Writing the single-trait competition in this way bounds the strength of competition between 0 and -2, which helps us understand the properties of the interaction matrix in a trivial way as we describe here. We start by ordering the $\gamma_i$ values, which are themselves generated a random uniform distribution $\mathcal{U}[0.3, 0.7]$. This is only a guess and we do not claim these values are necessarily rooted in empirical observations, so that any prediction from the model can only be qualitative. One useful aspect of defining competition as in the equation above is that competition strengths will always be bounded between 0 and -2 irrespective of the choice of $\gamma$ values, meaning that the lack of empirical parametrization of $\gamma$ only affects the model up to a certain extent.

The single-trait competition matrix $\tilde{A}$, once species are sorted by ascending $\gamma$ value, resembles the nested matrix discussed above: all $\tilde{A}_{ij} < -1$ for $i < j$. The upper triangle of the matrix is therefore filled with exclusionary interactions: species 1 will loose any competition with all other species, species 2 will loose except against species 1, and so on. Interactions in the lower triangle are not zero like in the fully nested model, but between 0 and -1, indicating a weak negative effect that does not lead to exclusion.

Yet, even in the case of single-nutrient competition discussed in (Chang *et al.*, 2023), one could consider that such perfect triangular architecture might be unrealistic, as we know that many other mechanisms could still be in place for example related to crowding or secretion of additional metabolites (Ghoul and Mitri, 2016). Given that the authors indicate a strong relevance of single-resource competition, we can consider that these additional mechanisms play a much weaker role. To do so, we consider a small random component in the competition matrix that is no longer mediated by the resource trait $\gamma$:

$$\tilde{A}_{ij} = -\frac{2\gamma_j}{\gamma_i + \gamma_j} - a_{ij}. \tag{32}$$

The small and random components of competitive interactions $a_{ij}$ are sampled from a normal distribution $\mathcal{N}(0.1, 0.01)$, giving small positive numbers that result in competition due to the addition of a minus sign in front of $a_{ij}$. In section II.A.4 we review the effects related to the presence or absence of this small competitive component: as expected, if $a_{ij} = 0$, the matrix is perfectly hierarchical, and there will not be any Rock-Paper-Scissors loops nor low-rank exclusions. It is therefore interesting to see whether high intransitivity is maintained under this additional random component and the positive cross-feeding term described below.

A last component modulating the strength of interactions described in (Chang *et al.*, 2023) is the presence of metabolic cross-feeding. In particular, the authors of (Estrela *et al.*, 2022; Goldford *et al.*, 2018) observe that lower-ranked competitors are able to survive likely due to better consumers secreting metabolic compounds that lower-ranked consumers can feed on to grow. This would indicate the presence of a second, and potentially independent layer of interactions beyond single-nutrient competition by which species facilitate the growth of one another. We refer the reader to e.g. (Pfeiffer and Bonhoeffer, 2004) for an in-depth review of this fascinating phenomenon. It could be that there appears to be a negative correlation between the consumption rank $\gamma_i$ and the feeding towards other species, so that stronger competitors are also those that secrete most metabolites to weaker ones (Estrela *et al.*, 2022; Goldford *et al.*, 2018). Yet, this could very well be an emerging property of the dynamics, and we decide to not introduce any a priori structure nor correlation in the properties of a cross-feeding matrix. In this context, we define the complete interaction matrix as

$$\tilde{A}_{ij} = C_{ij} - \frac{2\gamma_j}{\gamma_i + \gamma_j} - a_{ij}. \tag{33}$$

Now $C_{ij}$ are strictly positive values, given that all negative interactions can be captured in the last two terms of the right hand side described above. In the absence of additional information, we can parameterize the positive cooperation terms $C_{ij}$ by sampling them from a uniform distribution $\mathcal{U}[0.0, 1.0]$, meaning that their strength is approximately half the strenght of single-nutrient competition. We discuss in Section II.A.4 the implications of this assumption: because the GLV model does not have saturating interactions, it might be necessary to maintain positive interactions relatively lower to competition to allow the system to reach stable states instead of unbounded growth (See e.g. Figure 2A in the main text). Once again, we admit that we do not have empirical support for these parameters and use them only as a proof of concept to explain how hierarchical competition and random cross-feeding could lead to transitive EC. More detailed models of cross-feeding should incorporate explicit resource dynamics (van den Berg *et al.*, 2022; Mehta and Marsland III, 2021) or saturating responses (Aguadé-Gorgorió *et al.*, 2024b; Qian and Akçay, 2020) in order to describe more realistic scenarios with the goal of quantitative modeling. We discuss in Sections II.A.4, II.D the outcomes of this model in relation to the empirical observations of EC under strongly transitive architectures.

#### N. Other dynamical models

One of the central objectives of the present work is to show that EC, as a specific case study of an emergent collective or community-level property, can happen in simple models of pairwise interactions. The overall focus on the GLV model in our study is not because we believe that EC is a particularity of such a model; instead, we hypothesize that EC will happen in any system with many interacting species and strong enough competition, leading to a dense network of indirect “enemy of my enemy...” effects. The use of the GLV model in the main text and throughout this supplementary can be explained because it is a very simple (or the simplest) possible species-rich model and a widely-used method to describe interacting microbial species (Altieri *et al.*, 2021; Bunin, 2017; Galla, 2018; Hu *et al.*, 2022; Kessler and Shnerb, 2015; Mallmin *et al.*, 2024; Pasqualini *et al.*, 2024). More importantly, its simplicity provides interesting tools to analyse the weight of indirect effects via the collectivity metric (Zelnik *et al.*, 2024), the presence of exclusionary interactions via $A_{ij} < -1$ elements (Section I.B), and so on. Any other high-dimensional model will be much harder to treat analytically and the results much less appealing if we aim of conveying a simple and clear message.

In this section we present other dynamical models that we can at least simulate to search for EC in non-GLV models but still with strictly pairwise interactions. One of the important considerations here is that finding EC once we have reached a stable state is not as simple as before. In the GLV model, exclusionary interactions between two species will happen as soon as $A_{ij} < -1$, provided that $A_{ij}$ is the relative interaction strength, which is equivalent to saying $a_{ij} < a_{ii}$ where both elements are negative. Yet, other dynamical models need not have such a simple condition, and in fact obtaining a simple analytical expression for which we can look at a stable state and infer if some of its constituent species would not coexist in isolation might not be possible. In that case, what we propose in this section is a numerical exercise that is even closer to the original experiments proposed in (Chang *et al.*, 2023).

For each of the dynamical models proposed below, we will obtain numerically a tournament matrix for the outcomes of growing species by pairs. Similar to what is discussed in section I.E.1, we start from a large pool of $S$ interacting species and random interactions, and see if the dynamics lead to a stable state. Once a stable state is reached, and because we cannot simply search for $A^*_{ij} < -1$ elements, we simulate the dynamics for each of the $S^*(S^*-1)/2$ pairs that have survived together, again following the methods described in Section I.E.1 but with only two species present at $t = 0$, and evaluate if at the end of the dynamics there is any species in the pair that has gone extinct, indicative of competitive exclusion. This is a weaker method compared to the analytical test, and might be prone to errors in which a species has gone extinct due to fluctuations, yet we believe this might be a rare result and one should not expect fluctuating abundances in systems where we only simulate the dynamics of two species at a time. To further avoid this scenario, we do not start species pairs at random initial conditions, but rather at $x_i = 1$ to ensure abundances are not arbitrarily low at start. If any pair of species in the stable state shows a signature of competitive exclusion, we consider that the stable state contains an excluding pair, and so that Emergent Coexistence is present.

The numerical cost of performing pairwise tests of all possible pairs in a stable state, and repeating this process for many different interaction matrix randomizations, implies that the analysis of the models below is particularly slow. We have performed the simulations with $S = 30$ instead of $S = 80$, which we believe does not change the outcomes of the model beyond qualitative differences on the final diversity of stable states $S^*$. Because of cost of these simulations, we have focused on showing that the models harbor EC, meaning that there exist stable states where at least one pair of species would not coexist if isolated from the community. This analysis is consistent with the analysis for the GLV model of figure 2A of the main text. We leave for future work repeating the complete study of Figure 2C, Figure 3 and Figure 4 for the models detailed below. In any case, the results discussed in the Results section fulfill our goal, by which we see that EC is possible and again common in these models provided that similar conditions (which we are not able to obtain analytically) on interaction strength and heterogeneity are met.

#### 1. Saturating interactions with a Holling response

A particular property of the GLV model, easy to see when looking at its phase space (Figure 2A in the main text), is that cooperative interactions easily lead to unbounded growth (the right side of the phase space when $\mu > 0$). A natural method to obtain a more realistic dynamical system, where growth cannot be infinite, is to assume that interactions saturate at high population abundance (Yu *et al.*, 2019). This can be for example implemented by a Holling type-2 response (Castillo-Alvino and Marvá, 2020; Holling, 1959; Kvrivan and Eisner, 2006), so that the dynamical system under study would become

$$\frac{dx_i}{dt} = x_i \left( 1 + \sum_{j=1}^{S} A_{ij} \frac{x_j}{1 + x_j} \right). \tag{34}$$

In this model, the $A_{ij}$ can be sorted from an equivalent matrix as that proposed in the GLV model, where we control again $\mu$ and $\sigma$ and follow the same procedure of Section I.E.1 to draw a phase space of EC. Note here that we have included self-regulation inside of the sum, as opposed to the main text, where interactions and self-regulation are separated into two different matrices $(-I + A)$. This was useful for the mathematical analysis of indirect effects but does not have relevant implications for the model presented here. Another interesting comment is that, while saturating interactions lead to a more realistic setting of cooperation, where positive effects cannot lead to infinite growth, saturation of competition opens up questions regarding emergent coexistence. The strength of competition exerted by a dominant species will eventually saturate, imposing a potential limit to species dominance. In this setting, in which run-off effects like single-species dominance or overly strong indirect effects are not possible, we ask if emergent coexistence is still possible.

In section II.A.5 we present the results of this model again in terms of the fraction of communities with EC as a function of $\mu$ and $\sigma$, the mean and standard deviation of elements of the interaction matrix.

#### 2. Multilayer interactions with Allee Effects

The model discussed above is a direct modification of the GLV model to include more realism via the non-linearity of interactions. Yet, considering that all interactions follow the same functional response might be unrealistic in some cases, and saturating competition could blur out the capacity of the model to represent strong exclusionary interactions such as those seen in (Chang *et al.*, 2023). Furthermore, under weak interactions, the model predicts that all species will grow, meaning they can all invade from a very small $(x_i = 0 + \epsilon)$ abundance.

Two variations to the above model could be introduced in order to account for more realism. Following the model presented and studied in (Aguadé-Gorgorió *et al.*, 2024b), the first variation considers a multilayer architecture of interactions, where cooperation and competition are introduced in two different matrices $A$ and $B$ respectively (Pilosof *et al.*, 2017). This formalisms allows us to consider a richer dynamical configuration, by which species can compete and cooperate following different statistics ($\mu_A, \sigma_A$ and $\mu_B, \sigma_B$) and different functional forms as shown below. Additionally, the second variation that we can consider is a more realistic Allee Effect, by which a species requires a minimal abundance to grow, and below which it goes extinct (Courchamp *et al.*, 2008). This provides a rich set of dynamical scenarios, as each species can now grow or become extinct on its own even in the absence of interactions. We refer the reader to (Aguadé-Gorgorió *et al.*, 2024b) and (Altieri and Biroli, 2022) for a detailed analysis of the properties of species-rich models with Allee Effects.

Given these two considerations to include realism and dynamical richness in a community model, the dynamics of a given species in this system could be governed by

$$\frac{dx_i}{dt} = x_i \left( \sum_{j=1}^{S} A_{ij} \frac{x_j}{1+x_j} - d_i - \sum_{j=1}^{S} B_{ij} x_j \right). \tag{35}$$

Now, as opposed to the models above, the elements of matrix $A$ and $B$ are positive by definition. This implies that $A$ captures a saturating facilitation term, and $B$ describes linear competition as in the GLV model. The additional term $d_i$ is also positive-defined and includes a linear death rate term to the system. In the absence of interactions, this linear death term generates an Allee Effect as described in (Aguadé-Gorgorió *et al.*, 2024b).

To build and analyse this model, we generate positive-defined matrices $A$ and $B$ and vector $d$ from log-normal distributions using `numpy.random.lognormal` function. Because we want to focus on the role played by interactions, we fix the statistics of $d$ to $\mu_d = 0.1$ and $\sigma_d = log(1.1)$ following (Aguadé-Gorgorió *et al.*, 2024b) and focus on $A$ and $B$. To generate a phase space with two dimensions, we can for example fix the standard deviations of the cooperation and competition matrices to $\sigma_A = \sigma_B = log(1.1) \approx 0.09$ and modulate the mean interaction strengths of cooperation and competition as the x- and y-axis respectively. We refer the reader to (Aguadé-Gorgorió *et al.*, 2024b) where a detailed analysis of this model and its phase space is performed, and focus here only on studying if emergent coexistence is present.

#### 3. Sublinear growth dynamics coupled to linear interactions

Throughout this work we focus in the role of species interactions, consistent with the overall question of whether pairwise interactions are enough to explain community properties. Yet, the functional shape of growth rates can also modulate the dynamical properties of a system.

A traditional concept in the evolution of prebiotic replicators is to consider that replication rates could be sublinear with the abundance of replicating entities (Czárán and Szathmáry, 2000; Piñero and Solé, 2018; Szathmáry and Smith, 1997). Recently, research has proposed that these sublinear replication rates could govern the growth of ecological species in natural ecosystems, which could lead to particular community-level properties regarding increased ecosystem stability (Hatton *et al.*, 2024). The possibility that population dynamics are not mediated by linear growth terms could also play important roles in the dynamics of heterogeneous solid tumors (Aguadé-Gorgorió *et al.*, 2024a; Rodriguez-Brenes *et al.*, 2013). One minimal way of writing a sublinear growth model of

$S$ interacting species is

$$\frac{dN_i}{dt} = r_i N_i^k - d_i N_i - N_i \sum_{j \neq i}^{S} A_{ij} N_j. \tag{36}$$

Following the notation proposed in (Aguadé-Gorgorió *et al.*, 2025; Hatton *et al.*, 2024), here we are studying not the species abundances relative to their carrying capacity, $x_i = N_i/K_i$ as used for the GLV model, but simply the total abundance $N_i$ of each species in the original pool. Interestingly, this apparently subtle variation in the growth function has profound implications in the stability properties of the model, which are described in more detail in (Mazzarisi and Smerlak, 2024).

Yet, the model as it stands has a very particular property which was originally found in (Szathmáry and Gladkih, 1989), by which species never go extinct, leading to a *survival of all* even under strong competition, as opposed to the more classical *survival of the fittest* of the linear growth model. This result had profound implications in the original studies of Darwinian selection (which is not possible if all replicators always coexist) and the origins of life (von Kiedrowski, 1993), and is also relevant when studying scenarios where competition can lead to the extinction of certain species as is the case of microbial communities (Chang *et al.*, 2023; Friedman *et al.*, 2017; Venturelli *et al.*, 2018). When considering ecological communities of interacting species, and as described in detail in (Aguadé-Gorgorió *et al.*, 2025), in the sublinear replication model species cannot go extinct even if competition is infinitely strong. This mathematical artifact emerges because, at low abundance, competition decays linearly with abundance, but per-capita growth rates become extremely large and species are always able to survive (Aguadé-Gorgorió *et al.*, 2025). To solve this issue, a natural mechanism is to introduce a minimal abundance below which a species growth rate is not sublinear. In its simplest form, we assume that a species will go extinct below if the total population is below one individual, $N_i < 1$ (Aguadé-Gorgorió *et al.*, 2025). In the GLV model, we introduced a minimal abundance threshold of $10^{-20}$ below which a species goes extinct. This was not to ensure that species go extinct, which is a natural possibility in the GLV model, but rather to avoid that species have a negative abundance. The natural threshold in GLV could simply be to assume that a species goes extinct if $x_i \leq 0$. In the sublinear growth model, such a natural threshold of null density would not work, because species never reach it, and the threshold has to be explicitly higher.

To simulate the sublinear growth model, we generate $A_{ij}$ following the same dynamical properties of the model above and set $r_i = 1$ and $d_i = 0.05$. Following the simulations proposed in (Aguadé-Gorgorió *et al.*, 2025), we set a smaller extinction threshold of 0.03 which can be indicative of the biomass of a single individual in case the model is written in biomass terms instead of abundances (see the supplementary code `13_FIGS3D.py` as well as (Aguadé-Gorgorió *et al.*, 2025) for the full characterization). We describe in section II.A.5 the findings in terms of the presence of EC in the $(\mu, \sigma)$ phase space of the sublinear growth model.

### II. MAIN RESULTS AND ADDITIONAL OUTCOMES

#### A. Mathematical results for emergent coexistence in the Generalized Lotka-Volterra phase space

The first set of results exposed in the main text are contained in Figure 2, and can be summarized as "EC is a common and predictable outcome in the GLV model". In this section we explain the analytical developments to find the boundaries of the Emergent Coexistence domain (Figure 2A,B in the main text), that allow us to understand what are the minimal conditions for which EC can happen in the GLV model with random interactions. The two transition lines $\sigma_1^c$ and $\sigma_2^c$ are explained in Sections II.A.1-2, are calculated in the code for Figure 2B and are implemented in figure 2A following the results of figure 2B. In Section II.A.3, we explain the analytical and semi-analytical methods that allow us to estimate the maximum fraction of excluding pairs that a stable community with diversity $S^*$ can sustain, as shown in the red dashed line of figure 2C. Additional results considering other interaction matrices, other dynamical models and other connectivity values are also discussed. The central message in those is that EC happens across different assumptions and is not a particularity of the GLV model or the random interactions or high connectivity assumptions of the main text.

##### 1. The minimal competition necessary for emergent coexistence

Emergent Coexistence requires by definition that at least one element in the interactions between surviving species is exclusionary, meaning that $A^*_{ij} < -1$ for $i \neq j$. For this to happen, we need *at least* that the same condition applies to the complete pool of interacting species, so that $A_{ij} < -1$ for $i \neq j$. A necessary condition for EC is therefore that at least one element of the complete pool is exclusionary, or else EC will be completely impossible. The mathematics behind this rule are simple to obtain from basic probability. In a pool of $S$ species there are $S(S-1)$ interspecies interactions, each sampled independently from a Gaussian (Normal) distribution with mean $\mu$ and standard deviation $\sigma$,

$$A_{ij} \sim \mathcal{N}(\mu, \sigma^2), \quad \text{for } i = 1, 2, \ldots, S(S-1). \tag{37}$$

We want to compute the probability that at least one of these elements is smaller than $-1$:

$$P\left(\text{At least one } A_{ij} < -1\right). \tag{38}$$

The key intuition is that

$$P\left(\text{At least one } A_{ij} < -1\right) = 1 - P\left(\text{All elements } A_{ij} > -1\right), \tag{39}$$

which, furthermore, can be developed into

$$P\left(\text{At least one } A_{ij} < -1\right) = 1 - P\left(\text{All } A_{ij} > -1\right) = 1 - \left(1 - P\left(\text{One } A_{ij} < -1\right)\right)^{S(S-1)} \tag{40}$$

For a single normally distributed random variable $A_{ij}$, the probability that it is less than $-1$ is given by the cumulative distribution function (CDF) of the normal distribution:

$$P(A_{ij} < -1) = \Phi\left(\frac{-1-\mu}{\sigma}\right), \tag{41}$$

where $\Phi(z)$ is the CDF of the standard normal distribution:

$$\Phi(z) = \frac{1}{\sqrt{2\pi}} \int_{-\infty}^{z} e^{-t^2/2} dt. \tag{42}$$

Since the elements are independently sampled, the probability that all $S(S-1)$ elements are greater than or equal to $-1$ is:

$$P\left(A_{1,2} \geq -1, A_{1,3} \geq -1, \ldots, A_{S,(S-1)} \geq -1\right) = \left(1 - P(A_{ij} < -1)\right)^{S(S-1)}. \tag{43}$$

Thus, the probability that at least one element is smaller than $-1$ is:

$$P\left(\text{At least one } A_{ij} < -1\right) = 1 - \left(1 - \Phi\left(\frac{-1-\mu}{\sigma}\right)\right)^{S(S-1)}. \tag{44}$$

Interestingly, when $S$ is sufficiently large, the shape of this function approaches a step function. For example, given $S = 100$, $S(S-1) = 9900$, and a fixed $\mu$ value, there is a critical value for $\sigma_1^c$, below which the probability of finding exclusions approaches zero, and beyond which the probability of finding exclusions approaches one. This explains why EC is confined to a specific domain in the $(\mu, \sigma)$ phase space: there is a sharp boundary below which all interactions will result in pairwise coexistence. However, once this boundary is crossed and there is at least one exclusionary interaction in $A$, it does not mean that all stable communities will contain it. This explains why the fraction of EC states does not reach 1 sharply after $\sigma_1^c$ in figure 2B of the main text: there will be some stable states where $A^*$ contains the exclusionary values, and some will not. Once many $A_{ij} < -1$ values are present in $A$, most stable states will contain at least one $A^*_{ij} < -1$. But, by definition, EC is impossible when $P\left(\text{At least one } A_{ij} < -1\right) \to 0$.

To estimate the threshold value $\sigma_1^c$ for each given value of $\mu$ in Figure 2A and 2B of the main text, we equate the probability of finding at least one $A_{ij} < -1$ to 0.5, which marks the intermediate value of the transition between 0 and 1, and use the bisection method, a numerical root-finding technique. The bisection method is a bracketing method that iteratively reduces an interval $[a, b]$ containing the root by evaluating the function at the midpoint. If the function changes sign across an interval, the root must lie within that interval. The process is repeated until the interval is sufficiently small, guaranteeing convergence. In our implementation, the function `bisect` from the SciPy `optimize` library is used, which efficiently finds $\sigma_1^c$ within the predefined search range of [0.01, 1], ensuring numerical stability and accuracy.

It is noteworthy to observe that this line is not the same line defining the transition from the unique fixed point to the multistability regimes, originally found in (May, 1972) and accurately described in (Bunin, 2017). Our critical line $\sigma_1^c$ describes the probability that there is an exclusionary value in the pool $A$, and hence imposes a necessary condition for EC. The critical line separating mono- to multi-stability, marked as a dashed line in Figure 2A of the main text, is found using more advanced techniques from dynamical mean field theory (see e.g. (Bunin, 2017) or the many references discussed in the supplementary material of (Aguadé-Gorgorió *et al.*, 2024b)). To paint the dashed line in Figure 2A of the main text we have used the equations described in our previous work (Aguadé-Gorgorió and Kefi, 2024) and in (Mallmin *et al.*, 2024) and Section II.C.

In this context, and as discussed in the main text, it is also interesting to realize that the EC domain fall inside the multistability domain of the GLV model: there can be alternative stable states that do not have EC, but all states with EC can be considered alternative stable states. A thorough description of the properties of these alternative stable states in the GLV model can be found in (Aguadé-Gorgorió and Kefi, 2024; Biroli *et al.*, 2018). Despite this is only preliminary evidence and much work should still be done in this regard, it is interesting to pinpoint that the original EC experiments, described in (Goldford *et al.*, 2018), had signatures of multistability: different stable community compositions could be found by assembling the same species starting from different initial conditions. This could be another match between what the GLV model with random interactions predicts (EC happens in the regime where different initial conditions lead to alternative stable states) and what empirical observations show.

#### 2. The maximum competition allowed for emergent coexistence

The EC regime is also bound at the other extreme, in which, if interactions are too strong or too homogeneous, only a single species will survive and no stable multispecies communities can be found (the *competitive exclusion* regime, (Aguadé-Gorgorió *et al.*, 2024b)). We are interested in understanding this transition.

For a stable community with 2 species to emerge so that competitive exclusion is avoided, we need a pair of elements ($A_{ij}$,$A_{ji}$) to be larger than -1. This means, we need two species that do not exclude one another. The simple probabilistic rules determining this transition allow us to infer the transition line separating exclusion from multistability in the GLV map for the first time to our knowledge. We want to calculate the probability that at least one pair of elements, $A_{ij}$ and $A_{ji}$, from the matrix are both larger than $-1$. The key aspect of this problem is that the pair $(A_{ij}, A_{ji})$ should both exceed $-1$, not just each element independently. We need to calculate the probability that such a pair exists in the matrix.

We first need to compute the probability that both $A_{ij}$ and $A_{ji}$ exceed $-1$. Since $A_{ij}$ and $A_{ji}$ are independently sampled from the Gaussian distribution, the probability that both elements are greater than $-1$ is simply the product of the individual probabilities:

$$P(A_{ij} > -1) = 1 - \Phi\left(\frac{-1-\mu}{\sigma}\right)$$

where $\Phi(z)$ is the cumulative distribution function (CDF) of the standard normal distribution. Similarly, the probability that $A_{ji} > -1$ is:

$$P(A_{ji} > -1) = 1 - \Phi\left(\frac{-1-\mu}{\sigma}\right)$$

Thus, the probability that both $A_{ij}$ and $A_{ji}$ are greater than $-1$ is:

$$P(A_{ij} > -1 \text{ and } A_{ji} > -1) = \left(1 - \Phi\left(\frac{-1-\mu}{\sigma}\right)\right)^2$$

For simplicity, let us name this pair event as $C = A_{ij} > -1$ and $A_{ji} > -1$. This means, $P(C)$ is the probability that there is one coexisting pair

We now want to find the probability that *at least* one pair $(A_{ij}, A_{ji})$ satisfies the condition $A_{ij} > -1$ and $A_{ji} > -1$, which resembles the mathematical development we discussed above for *at least* one pair $A_{ij} < -1$. The probability that at least one pair satisfies the condition is:

$$P\left(\text{At least one } C\right) = 1 - P\left(\text{No } C\right), \tag{45}$$

The probability of $P\left(\text{No } C\right)$, i.e., none of the $S(S-1)/2$ pairs satisfying $C = A_{ij} > -1$ and $A_{ji} > -1$, is given by:

$$P\left(\text{No } C\right) = 1 - (1 - P(C))^{S(S-1)/2} \tag{46}$$

Substituting for $P(C)$, we obtain the final expression:

$$P\left(\text{At least one } C\right) = 1 - \left(1 - \left(1 - \Phi\left(\frac{-1-\mu}{\sigma}\right)\right)^2\right)^{S(S-1)/2} \quad (47)$$

This provides the probability that at least a pair of two elements in $A$ are greater than $-1$, given that each element follows an independent Gaussian distribution with mean $\mu$ and standard deviation $\sigma$. This equation also defines for the first time to our knowledge an estimate for the line separating the exclusion and the multistability domains of the GLV map (Aguadé-Gorgorió and Kefi, 2024; Mallmin *et al.*, 2024), marked again with a bold line in Figure 2A. The rationale is the same one as described in the section above: when $S$ is sufficiently large, this function has a very sharp transition from 0 to 1 (see Fig. 1), which we can use to settle $\mu$ and find $\sigma_2^c$ with the same numerical method as above.

Yet, a community with two coexisting species will not necessarily have EC: we need a third species and one exclusionary element. One possibility for this is that there is a coexisting pair, as derived here, and that this coexisting pair maintains a third species in which one of the coexistors excludes it, the other one rescues it, something similar to what we have discussed in Section I.B.2. Another possible structure, as we know from Rock-Paper-Scissors triplets, discussed originally in e.g. (Gilpin, 1975) and in Section I.L.2, a community with 3 species can be stable and show EC if at least three elements of the interaction matrix $A^*$ are larger than -1. We want to find the probability that there are 3 exclusionary elements forming a rock-paper-scissors loop while the other three elements are non-exclusionary.

The probability for finding these structures is harder to infer analytically. Yet, we can easily predict the shape of the function by sampling a large set of matrices at random with elements sampled from a normal distribution with fixed $\mu$ (e.g. $\mu = -1.5$ for the second panel of Fig. 2B) and increasing $\sigma$, and plot the fraction of matrices that have a coexisting pair and a potentially coexisting RPS triplet. This means a triplet for which all three elements are larger than 1, although not all triplets that have this structure will lead to stable coexistence. Interestingly, we find that the probability of finding a coexisting pair, $\{A_{ij} > -1 \text{ and } A_{ji} > -1\}$, is very similar to the probability of finding a RPS triplet of elements larger than 1 (Fig. 1). This indicates that the above analytical estimate for a coexisting pair might be good enough as an estimate for the boundary beyond which stable rock-paper-scissors triplets can appear.

#### 3. The maximum fraction of excluding pairs that a community can sustain

In Figure 2C of the main text we plot the fraction of excluding pairs in stable communities with different diversity emerging from different realizations of the GLV model (that is, different initial conditions and $A \sim \mathcal{N}(\mu, \sigma)$ in the range of Figure 2A). Our aim here is to describe the mathematical developments to obtain an estimate for the maximum number of excluding pairs that we expect to see in communities of size $S^*$, corresponding to the communities that can sustain more competition.

To do so, we bring together the following mathematical developments:

- Given $(\mu, \sigma)$, we predict the most diverse state $\max(S^*)$ that the model will generate.
- Given $(\mu, \sigma)$, we predict the maximum fraction of exclusionary elements that can be found in $A^*$.

For the first point, as we previously showed in (Aguadé-Gorgorió and Kefi, 2024), we can estimate some properties of the alternative stable states that emerge from the GLV model given a $(\mu, \sigma)$ pair inside the multistability domain of Figure 2A in the main text. This comes from the question: given a large pool characterized by an interaction matrix $A(\mu, \sigma)$, what are the diversities $S^*$ and interaction statistics $(\mu^*, \sigma^*)$ of the stably coexisting subsets of species from the pool?

Given $(\mu, \sigma)$, one can show (Aguadé-Gorgorió and Kefi, 2024) that the most cooperative stable community that can emerge will have

$$\max(\mu^*) \approx \mu + 4\frac{\sigma}{\sqrt{S^*(S^*-1)}}. \tag{48}$$

That means, a state can be more cooperative than the original pool (by selection of the less-competitive species). Yet, as we increase the size of the subset, its statistics will resemble those of the original pool. This result is a well-known property in the statistics of the exercise of sampling a small subset of $N = S^*(S^*-1)$ elements from a large set of independent randomly distributed variables (Johnson *et al.*, 2002).

The opposite could be true regarding the most competitive subset. Yet, the strength of competition in stable states in the GLV model is bounded. From (Bunin, 2017) or (Mallmin *et al.*, 2024), one can show that the most competitive clique will have mean interaction strength following

$$\min(\mu^*) \approx \sqrt{\frac{S^*}{2}}\sigma^* - 1, \tag{49}$$

which, in the correction for small subsets of $S^* \lesssim 50$ found by (Aguadé-Gorgorió and Kefi, 2024), writes

$$\min(\mu^*) \approx \frac{(S^*)^{1.08}}{14.118}\sigma^* - 1. \tag{50}$$

This equation indicates that a larger state $(S^*)$, or one in which interactions are more heterogeneous $(S^*)$ will be able to sustain less competition (a more positive $\mu^*$ value). Yet this equation depends on the standard deviation of interactions in the stable state, $\sigma^*$. As it will later be obvious, our aim is to obtain the statistics of the state as a function of the original pool statistics, $\mu, \sigma$. Now, there is also a closed expression that links the standard deviation of a subset of size $S^*(S^*-1)$ with the statistics of the original distribution. As seen in (Aguadé-Gorgorió and Kefi, 2024) (Fig. 6 of the supplementary material), the smallest standard deviation of a stable community will be

$$\min(\sigma^*) = 1 - \sqrt{1 - \frac{2}{S^*(S^*-1)-1}\left(\frac{\Gamma_1}{\Gamma_2}\right)^2}, \tag{51}$$

where

$$\Gamma_1 = \Gamma\left(\frac{S^*(S^*-1)}{2}\right) \tag{52}$$

$$\Gamma_2 = \Gamma\left(\frac{S^*(S^*-1)-1}{2}\right) \tag{53}$$

and $\Gamma(\cdot)$ is the Gamma function (Davis, 1959). Now we have a closed expression for the most cooperative stable state of size $S^*$ emerging from $\mu, \sigma$:

$$\max(\mu^*) \approx \mu + 4\frac{\sigma}{\sqrt{S^*(S^*-1)}}, \tag{54}$$

and the most competitive stable state of size $S^*$ emerging from $\mu, \sigma$:

$$\min(\mu^*) \approx \frac{(S^*)^{1.08}}{14.118} 1 - \sqrt{1 - \frac{2}{S^*(S^*-1)-1}\left(\frac{\Gamma_1}{\Gamma_2}\right)^2} - 1 \tag{55}$$

In (Aguadé-Gorgorió and Kefi, 2024) we showed that the crossing of these two functions ($\max(\mu^*) = \min(\mu^*)$) gives a statistical estimate for the most diverse stable state $S^*$ that can be found given a large pool of species with interaction statistics $(\mu, \sigma)$. The first step in our process is therefore the following: as said in the beginning of this section, we generate a large number of systems with $(\mu, \sigma)$ sampled randomly within the domain of figure 2A. For each $(\mu, \sigma)$ pair, we equate ($\max(\mu^*) = \min(\mu^*)$) as described above, and solve the highly nonlinear equation using a simple brute force method, by which we find the $S^*$ value within 2 and 50 that generates the smallest error $\epsilon$, where $\epsilon = \max(\mu^*) - \min(\mu^*)$. Now the first step is done: we have an estimate for the largest $S^*$ that can appear for every possible system. Of course, whether a state with that diversity or a smaller one is attained will depend on the initial conditions and we cannot know, yet the large number of simulations ensure that statistically this upper bound is found.

We then ask a different question. Given once again the statistics of the original pool alone, $(\mu, \sigma)$, can we predict the fraction of excluded pairs in the interaction matrix of a stable, $A^*$? First, it seems reasonable to assume that the maximum fraction of excluded pairs in the final state will be, at maximum, the maximum fraction of excluded pairs of the original pool $A$. To confirm this hypothesis, we test it numerically, confirming indeed that both the fraction of excluding elements and the fraction of excluding pairs is equal or smaller in $A^*$ than in $A$ (not shown): there is no reason to think that the final stable state would select more exclusionary interactions than the original pool, given that excessive competition is in general detrimental for coexistence.

Now, it suffices to ask, given $(\mu, \sigma)$, what is the fraction of exclusionary pairs in $A$, for which either $A_{ij}$ *or* $A_{ji}$ will be smaller than -1. As reviewed above, the probability that at least one of the elements is smaller than -1 is the one minus the probability that both $A_{ij}$ *and* $A_{ji}$ are larger than -1. Since $A_{ij}$ and $A_{ji}$ are independently drawn from the same Gaussian distribution, the probability that at least one of them is less than $-1$ is similar to the development above:

$$P(A_{ij} < -1 \text{ or } A_{ji} < -1) = 1 - P(A_{ij} \geq -1)P(A_{ji} \geq -1) \tag{56}$$

Using the Gaussian CDF, we have:

$$P(A_{ij} \geq -1) = 1 - \Phi\left(\frac{-1-\mu}{\sigma}\right) \tag{57}$$

$$P(A_{ji} \geq -1) = 1 - \Phi\left(\frac{-1-\mu}{\sigma}\right) \tag{58}$$

Thus, the probability of at least one being smaller than $-1$ is:

$$P(A_{ij} < -1 \text{ or } A_{ji} < -1) = 1 - \left(1 - \Phi\left(\frac{-1-\mu}{\sigma}\right)\right)^2 \tag{59}$$

Since there are $S(S-1)$ unique unordered pairs $(i, j)$ (excluding the diagonal elements), the fraction $f$ of such pairs satisfying the condition is equivalent, giving

$$f = 1 - \left(1 - \Phi\left(\frac{-1-\mu}{\sigma}\right)\right)^2 \tag{60}$$

We have reached the second step of our method. Now, for a given $(\mu, \sigma)$ pair, the first step provided the maximum possible diversity that a stable state can reach, $S^*$, and the second step provides an estimate for the maximum fraction of exclusionary pairs that the original pool $A$ can harbor. To bring these two analytical results together, an easy implementation algorithm is to record each ($S^*$,$f$) pair that our method finds given each $\mu, \sigma$ pair. Each time a larger $f$ value associated to a given $S^*$ value is found, we update it. At the end of a long simulation, this two-sided analytical method (where an estimate solution of $S^*$,$f$ is found by bridging two analytical solutions for $S^*$ and $f$ both as a function of $\mu$ and $\sigma$) has allowed us to find the largest fraction of excluding pairs $f$ associated to the most competitive states for each possible $S^*$ value (Fig. 2C in the main text, red dashed line).

##### 4. Emergent coexistence under other interaction matrices

In Figure 2 we explore the presence of EC in the GLV model where different types of random interaction matrices have been implemented. The different types of interaction matrices and how to build them are discussed in Sections I.C, I.D and I.M. In very general terms, and without aiming at a precise description of the dynamical properties of these different systems, figure 2 shows that EC is a very common outcome not only for random, Gaussian and uncorrelated $A_{ij}$ elements as in the main text, but also for antisymmetric, triangular, sparse and skewed distributions and the more specific single-resource and cross-feeding interaction matrix proposed in the main text. Instead, fully symmetric interactions where $A_{ij} = A_{ji}$ do not yield EC. This is easy to understand for the three-species and indirect effects system described in Section I.B, where heterogeneous interactions are necessary, or the three-species rock-paper-scissors scenario discussed in Section I.M: it appears as if a rock-paper-scissors loop can be stable provided that there is a loop in only one of the two directions: if $A_{ij}$, $A_{jk}$ and $A_{ki}$ are smaller than -1, the opposite elements must be larger than -1 to maintain the system in place. Furthermore, as signaled in (Koch *et al.*, 2024, 2023), there is an intrinsic need for asymmetry in the interactions to allow for certain hierarchy. Consistent with our observations of few low rank exclusions, it could be that strong symmetry of interactions dismantles this weak hierarchies and makes a system all too similar. With this intuitions in place, it remains an open question to analytically show why EC does not happen in fully symmetric interaction matrices.

Additionally, we can see that nested, sparse and antisymmetric phase spaces look similar to the main phase space of the GLV model presented in figure 2A of the main text (Fig. 2A-C). Also the phase space of symmetric interactions looks similar (not shown, but see (Bunin, 2017)), but without EC. Instead, the phase space of the GLV model with Gamma distributed interaction matrix looks different, mainly because the meaning of the axes is different. For the gamma distribution (Fig. 2D), we are modulating the shape $\alpha$ and scale $\theta$ parameters instead of mean $\mu$ and standard deviation $\sigma$. In this context, the mean of the distribution is $\mu = \alpha\theta$, explaining why the phase space reminds us to that of (Hu *et al.*, 2022), where both mean and standard deviation of competition are increased systematically. The standard deviation of the distribution is $\sigma = \theta\sqrt{\alpha}$. Finally, the positive side of the graph does not make sense, because the gamma distribution is positive definite and we add a negative sign to ensure interactions are competitive, explaining why the phase space is symmetrical at $x = 0$.

To plot a possible phase space for the single-nutrient competition and positive cross-feeding interactions, we fix the competition terms as described in Section I.M and modulate the strength of cooperation through the mean and standard deviation of the $C_{ij}$ elements (Fig. 2E). It is important to remember that the elements of $C_{ij}$ are intended to describe facilitation and are sampled from a positive-defined normal distribution in Figure 4 in the main text, but here we generate them from a Gaussian distribution without the only-positive constraint to better understand their implications. In particular, we can see that if the mean of $C_{ij}$ is too negative, the matrix describes competition and the overall system cannot produce species-rich EC states due to excessive competition resulting from adding $C_{ij}$ competition to the additional single-resource and random elements. Instead, species-rich communities start to emerge once $C_{ij}$ elements approach a null mean that is larger than the expectation for the random case, and high-enough heterogeneity, so that enough cooperation is present to overcome the single-resource component, and multispecies communities can survive, also explaining our choice for strictly positive cross-feeding $C_{ij}$ in the main text. Because these multispecies communities will emerge with an interaction matrix that has an exclusionary component through single-resource interactions, they harbor excluding pairs almost by default (strong purple color, Fig. 2F). All in all, these simple preliminary tests provide a glimpse that EC is a common outcome across interaction matrices: it requires that interactions in the pool are heterogeneous and competitive enough, but it does not seem to care about more subtle details of their distributions as long as interactions are not excessively symmetrical.

#### 5. Emergent coexistence in other dynamical models with pairwise interactions

As shown in figure 2A of the main text, Emergent Coexistence is found inside of a large regime of the GLV phase space where previous work had found the presence of multistability (Aguadé-Gorgorió and Kefi, 2024; Altieri *et al.*, 2021; Biroli *et al.*, 2018; Bunin, 2017). The intuition is that both Emergent Coexistence and multistability require strong interactions and strong indirect effects, which is also at the core for why long and persistent abundance fluctuations are found in this phase (Gilpin, 2024; Mallmin *et al.*, 2024; Roy *et al.*, 2020). As seen above, the requirements for EC are slightly more specific, as it requires not only strong competition, but more explicitly $A_{ij} < -1$.

Previous work showed that such a multistability phase, where multiple stable species combinations (*cliques*) are possible, is not a particularity to the GLV model (Aguadé-Gorgorió *et al.*, 2024b). Instead, a qualitatively equivalent phase could be found in complex systems models including positive saturating interactions, an Allee Effect, models of gene regulation or neuron cluster interactions or models of cancer-immune interactions (see supplementary material of (Aguadé-Gorgorió *et al.*, 2024b)). In each of these models, strong and heterogeneous competition led to cliques: heterogeneity in interactions implies the presence of intrincate indirect effects, where a species can be a strong competitor against one species, a positive cooperator against another, and so on, in a strictly disordered fashion. Albeit we still do not have a mathematical definition to characterize this phenomenon more strictly, the observation is that a similar multistability regime is found across models and is not dependent on the linearity of the GLV model.

This suggests an equivalent expectation for emergent coexistence: we hypothesize that emergent coexistence is possible in all pairwise interaction models inside their regime of strong indirect effects and multistability. In fact, there is no reason to suspect that EC is a particularity of the GLV model, but rather an emerging property in ecological communities where species are embedded in strong and densely connected interaction networks. The focus in the GLV model in the main text work is mainly because its simplicity provides analytical and intuitive results.

In Section I.N we have described other archetypic models of species-rich interactions that include saturating responses, multilayer interactions or sublinear growth. Additionally, following the discussion provided at the end of this supplementary material, we also consider in this section and in the same figure a GLV model with external migration where stable states also have to be resistant to reinvasions by species from the pool (see Section II.E). Due to the cost of simulating these models, we restrict our question in them to asking: are there stable states where at least one pair of species cannot coexist in isolation due to competitive exclusion? To explore this, we repeat the basic analysis of Figure 2A but for each of the dynamical models presented in Section I.N.

As discussed in detail in Section II.E, the central property of interest in the GLV model with external migration is that many of the stable states found in this work become unstable, leading to a system where persistent fluctuations exist as a pinball between multiple (un)stable states (Aguadé-Gorgorió and Kefi, 2024; Gilpin, 2024; Mallmin *et al.*, 2024; Roy *et al.*, 2020). Yet, we find in figure 3A that, even if fewer stable states exist due to external reinvasions, those stable states that are present harbor EC very often, with fractions as large as those found in the main text for the GLV model without migration. The intuitive conclusion is that, with or without external migration, strong interactions lead to a regime of dense and dominant indirect effects leading again positive net effects and emergent coexistence.

In the model with saturating interactions, one of the interesting properties is that now excessive cooperation cannot lead to runaway effects nor unbounded growth. Instead, it ensures the coexistence of all species. Because $A_{ij}$ are sampled as in the GLV model above with migration and as the model of the main text (Fig. 2A), the cooperation phase now has an intuitive result. We have shown in Fig. 2A,B and in Section II.A.1 that there is a minimal $\mu, \sigma$ combination necessary for $A$ to carry exclusionary elements. Now, with saturating cooperation, all species will coexist at positive $\mu$. The interesting result, presented in figure 3B, is that now EC is not only present in the regime of the GLV model. Additionally, EC can be found in this cooperative regime, where all or most species are present, and EC is ensured because there will always be EC in the pool of interactions following the results of Section II.A.1. The intuitive conclusion, once again, is that, with or without saturating interactions, EC is possible once interactions are strong enough so that indirect effects dominate and positive net effects can happen. This is possible even when competitive interactions also follow a saturation curve: even when dominance is restricted by a Holling type-2 function, indirect effects accumulate and survival of excluding pairs is possible in larger communities.

We find that EC is also a common, albeit less pervasive outcome in the model with multilayer interactions separated in competition ($B$) and cooperation ($A$) matrices (Fig. 3C). This could be related to the fact that we fix heterogeneity to be relatively low ($\sigma_{A,B} \approx 0.1$). In any case, and following the analysis of the phase space of this model performed previously in (Aguadé-Gorgorió *et al.*, 2024b), we find that the multistability phase is again pervaded by emergent coexistence. Outside of this phase and as discussed in (Aguadé-Gorgorió *et al.*, 2024b), we find that either cooperation dominates and most species pairs coexist without excluding one another, or that competition dominates and the system only reaches final states with one surviving species. The presence of saturating cooperative interactions leads to a different shape of the phase space, but does not alter the fact that, once competition is sufficiently strong and heterogeneous, strong indirect effects become pervasive and excluding species can coexist. Despite this is only an intuition and we lack analytical results for this model, we hypothesize that the main qualitative results discussed in the main text also apply here.

In Figure 3D we plot a phase space for the sublinear growth model where the x-axis controls the mean and the y-axis the standard deviation of the matrix of linear interactions. As discussed in (Hatton *et al.*, 2024; Mazzarisi and Smerlak, 2024), this model leads to increased stability of all possible states once diversity increases, which could have important implications as opposed to the increasing instability leading to permanent fluctuations and extinctions in the GLV model. Beyond stability and focusing on species coexistence, whether more or less species coexist in a given $\mu, \sigma$ range depends strictly on the extinction threshold imposed: in the absence of an arbitrary threshold, species never go extinct in the sublinear growth model and their diversity can become infinitely small. We find in figure 3D that EC becomes an extremely common outcome in the sublinear growth model. The fact that EC is present should not be surprising at this stage: many pairwise interactions can lead to positive indirect effects and rescuing of otherwise-excluded species. We hypothesize that the fact that EC is so common is related to the increased diversity effects of the sublinear growth term (Mazzarisi and Smerlak, 2024): Because many species can coexist at very low abundance provided they are above the given threshold, the sublinear growth model is able to generate states with competition but an arbitrarily large number of coexisting species. Equivalent to the regime where many species coexist thanks to cooperation in figure 3B, the regime where many competitors coexist even if at very low abundance ensures that the presence of at least one excluding pair is almost granted. Using higher extinction thresholds is likely to move the phase space to a regime with coexistence properties more similar to those of the GLV model (Aguadé-Gorgorió *et al.*, 2025).

Beyond the studied models with migration, saturating responses, multilayer interactions or sublinear growth rates, we hypothesize that EC would also be found in other dynamical models that are not restricted to a single equation or dynamical compartment. The two most characteristic scenarios would be those of Consumer-Resource models or Plant-Pollinator interactions. In these models, we hypothesize that coexistence of otherwise excluding species will be possible once interactions are strong, so that consumers or pollinators can indirectly facilitate one another by strongly impacting intermediate consumer or pollinator “enemies”. While we do not have yet definitive results on these much more complex systems, the results above provide strong arguments towards the intuition that EC is a common outcome in systems where pairwise interactions are strong and heterogeneous, irrespective of the specific dynamical rules of these species-rich systems. This further reinforces the notion that the GLV model is only used in the main text as a simple and treatable phenomenological description. Even when additional, more realistic properties are considered, the observation that EC is common is consistently recovered.

**Extended figure S3 caption:** *In purple the fraction of stable states that contain at least one excluding pair, the signature for emergent coexistence. We can see that, in different domains and under different frequencies, EC is a relatively common outcome across dynamical models. Dark dashed lines indicate regime boundaries for which we do not know the analytical expression, whereas red lines indicate the boundaries defined above. In (A) we study the GLV model using the equivalent numerical range of figure 2A of the main text with an additional migration term. Although the fraction of stable states becomes smaller and permanent fluctuations start to dominate, those stable states that are present are still found to harbor EC. This is an expected result as the presence of migration should not be expected to alter the results on positive indirect effects discussed throughout this work. In (B) we study a variation of the GLV model in which interactions are not linear but saturate with species abundances, using again the equivalent numerical range of figure 2A of the main text. This leads to the disappearance of the unbounded growth regime at high competition due to saturating cooperation limiting species outgrowth. Now we can see that the high-cooperation regime contains states where most species are present and EC is almost always ensured: because the* $\sigma_1^c$ *line is trespassed, almost surely at least one excluding pair will be present, and thanks to cooperation it will be maintained in the community. In (C) we show the phase space of the multilayer model where cooperation and competition are separated into interaction matrices. We fix the standard deviation to a relatively small value of 0.1 and modulate the mean of positive and negative interactions in the x-axis and y-axis respectively. The whole phase space was previously analyzed in detail in (Aguade-Gorgorio et al. 2024), and here we consistently find that the multistability domain contains a fraction of stable states where at least one pair of species do not coexist outside of the community. In (C) we plot a phase space of the sublinear growth model within the same parameter range of figure 2A of the main text but of course different equations discussed in Section I.N.3. We find a slightly different organization of the phase space, but again a large regime where most states harbor at least one excluding pair. The sublinear growth model can often sustain more coexisting species than the GLV model provided that the extinction threshold is low-enough. Because of this, we find that most states harbor EC, because they sustain a large number of species out of which it becomes likely that at least one is an excluding pair, similar to what we observe in the high-cooperation domain of panel (B). All the details on model parametrization and discussion of the phase spaces are discussed in I.N.3 and II.A.5.*

In the main text and the results presented above, we have focused on the case where the connectivity of the interaction matrix is set to $C = 1$. This choice simplifies the mathematical framework, allowing us to isolate and examine the roles of key parameters such as $\mu$ (the mean interaction strength) and $\sigma$ (the interaction variability). However, in Section I.D, we have discussed various empirical scenarios where network connectivity can be lower than $C = 1$ in empirical communities. The interesting result for the present work is that EC happens in the GLV model for both high or low connectivity (see section EC), and hence $C$ is not a central parameter in our study. Yet, we present here some preliminary results on the effects of decreasing $C$.

As shown in Figure 2C of the main text, the diversity observed in our simulations is significantly lower than what is typically found in real-world communities, which often contain hundreds or thousands of coexisting species. This discrepancy aligns with the long-standing diversity-stability debate, which discusses why increasing species diversity does not necessarily lead to greater stability of coexisting species in ecological models. The phenomenon by which increasing diversity leads to increased extinctions persists even when alternative growth models, such as sublinear growth dynamics, are considered (Aguadé-Gorgorió *et al.*, 2025; Hatton *et al.*, 2024). The diversity observed in Figure 2C is instead consistent with that of synthetic communities assembled under laboratory conditions, possibly highlighting a less sparse interaction matrix in those systems related to the fact that all species are grown on a single or few limiting resources and hence will most likely interact, whereas the available niches in natural ecosystems with many functional groups are many more than just those few available resources (Chang *et al.*, 2023; Friedman *et al.*, 2017; Venturelli *et al.*, 2018).

It is important to clarify that our study does not attempt to resolve the well-known diversity-stability paradox nor its implications for coexistence or biodiversity. Instead, we refer the reader to the works of (Bunin, 2017; Grilli *et al.*, 2017a; Marcus *et al.*, 2024; Serván *et al.*, 2018), who have extensively analyzed how coexistence and diversity emerge within the Generalized Lotka-Volterra (GLV) framework. Our focus lies elsewhere: we are interested in understanding how strong interactions shape species composition and whether these outcomes can be predicted from pairwise interaction data. The decision to use $C = 1$ was motivated by the desire for mathematical simplicity and consistency with prior studies on the GLV model (Aguadé-Gorgorió and Kefi, 2024; Bunin, 2017; Mallmin *et al.*, 2024). As we discuss in the previous section, our main findings regarding EC remain robust even under different connectivity values.

Nonetheless, an expected trend emerges when we relax the assumption of full connectivity. Unsurprisingly, reducing the connectivity of the interaction matrix increases the number of species that can coexist in the system. In figure 4, we illustrate how the species diversity of stable states in the GLV model changes as connectivity is reduced. It is interesting to note that not only diversity, but the shape of the phase space, changes as we modulate $C$. In particular, we find the following scenario: for large $C \approx 1$ (as in figure 2A of the main text and figure 4A), many species can coexist if $\sigma$ is very low and competition is not exclusionary. This is because low $\sigma$ leads to a neutral scenario where all species competitive capacity is similar. Differences in competition mediated by $\sigma$ rapidly lead to the extinction of some species. At intermediate $C$, we will see that heterogeneity is effectively increasing, and the effect on species diversity will not be so obvious. At very low $C$ we observe a clearly different pattern, where many more species can coexist (Fig. 4C). This non-monotonic trend can be easily understood. If we define $A_{ij} \sim \mathcal{N}(\mu, \sigma)$ as a normally distributed random variable, and define a new variable $B_{ij}$ as follows (the modified matrix with some zeroes):

$$\begin{cases} 0 & \text{with probability } 1 - C, \\ A_{ij} & \text{with probability } C. \end{cases}$$

The mean of the modified interaction matrix is $\mathbb{E}[B_{ij}] = C\mu$, meaning that interactions are on average weaker, while the standard deviation is $\text{std}(B_{ij}) = \sqrt{C\sigma^2 + C(1-C)\mu^2}$ (see supplementary material of (Aguadé-Gorgorió and Kefi, 2024)). The intuition emerging from this non-monotonic standard deviation is that slightly decreasing $C$ increases the heterogeneity of interactions and might therefore decrease coexistence (some strong interactions, some zeroes). Instead, only very large decreases in $C$ decrease the effective heterogeneity by centering most interactions at zero (Fig. 4, bottom panels). The result is that at low connectivity, coexistence is much less sensitive to changes in $\sigma$: increasing $\sigma$ increases the likelihood that strong competitors are in place in $A$ and hence other species go extinct, yet because we are transforming many of these strong $A_{ij}$ elements to zero, we are in fact reducing the effective heterogeneity of interactions and peaking most interaction elements at zero. The monotonic shape of $\text{std}(B_{ij})$ implies that this is only clear at very low connectivity values (Fig. 4, bottom panels). It is beyond the scope of the present work to analytically demonstrate how the phase space and species diversity of the GLV model changes with modified connectivity. Instead, the result that interests us is that, even for variable connectivity and species richness, the GLV model harbors a large domain in the phase space where EC is common. In this regime, reducing connectivity allows for more species to coexist (Fig. 4).

We do not claim that low connectivity is the primary driver of the remarkable diversity observed in natural ecosystems. Extensive theoretical and empirical research has explored the question of explaining the presence and stability of high leveles of natural biodiversity with mathematical arguments and models, and we refer the reader to the broader body of literature on the subject (see e.g. (Calleja-Solanas *et al.*, 2022; Chesson, 2000; Grilli *et al.*, 2017b; Hatton *et al.*, 2024; Ives and Carpenter, 2007; Jacquet *et al.*, 2016; May, 2019; Mazzarisi and Smerlak, 2024; McCann, 2000; Tilman *et al.*, 1998) among an extremely long list). However, it is worth noting that some studies suggest that low connectivity may explain key empirical patterns in microbial and macroorganism communities (Arya *et al.*, 2023; Camacho-Mateu *et al.*, 2024; Marcus *et al.*, 2022). In this context, it is possible that laboratory-assembled microbial communities exhibit lower diversity due to increased interaction connectivity (all species grow on a few resources or cross-fed metabolites), whereas natural ecosystems support a vast number of species, many of which belong to different and distant functional groups and therefore may not interact directly.

### B. Indirect effects, collectivity and condition number

#### 1. Interaction patterns in $A^*$ and coexistence

It is interesting here to recall the step-wise process behind our research. Given the results discussed in the sections above and in the main text, by which we saw that EC is a common outcome in the GLV model under different assumptions and hence does not need in principle any higher-order effect, we started to search for potential explanations and mechanisms. In Section II.D below we describe the observations regarding intransitivity, by which we realized early on that intransitive loops could be present but where not necessary, and perfectly triangular matrices without any loop could already generate EC. In that regard, we started a process to find microscopic (interaction-level) structures in the patterns of interactions that could explain EC, such as those depicted in Figure 1B of the main text.

A powerful example in this regard concerns the work in (Barbier *et al.*, 2021). In it, the authors study the following question: given a pool of species with random interactions, sampled as in our work, is there any non-random signature in the interactions of those species that coexist? The authors found two diffuse or subtle signatures in both theoretical models and empirically estimated interaction matrices from plant communities. These signatures indicate that (1) high-abundance competitors subtly favor each other by not competing directly, and (2) high-abundance competitors tend to target different species. In very general terms, these signatures indicate that competition is somehow distributed across the network, so that e.g. a species is not targeted by two strong competitors at once. We later found that these patterns are also present in cliques, the alternative stable states of the GLV model where we now find EC (Aguadé-Gorgorió and Kefi, 2024). Yet, because these states often contain few species, any statistical pattern is likely to be diffuse. A recent and insightful approach is also considered in (Poley *et al.*, 2025).

A similar exercise would then be to find if there are microscopic patterns in the organization of species interactions that explain EC in particular. To explore this, we performed the following test: we sampled random subsets of interacting species from $A$ following the procedure of Section I.E.2. For each system, we recorded the interaction matrix $A^*$, and studied a given metric of it. We divided the outcomes by systems that were not feasible (row sums of $(I - A^*)^{-1}$ are not positive, species cannot survive at positive abundance) and systems that were feasible. Between the feasible systems, we also divided those with positive abundance but positive eigenvalues (linearly unstable) from those with negative real parts of all eigenvalues (linearly stable). We then ask: for a given studied metric, is there statistically significant differences between each of these systems?

The first relevant part would be to find the conditions that lead to positive abundances (feasible subset of species) irrespective of linear stability, which we study below. The central rationale was that, given that a species $i$ is highly targeted by a competitor $j$ ($A_{ij} < -1$), one would expect to find some signature of other species targeting $j$, so that positive indirect effects emerge in the form of $A_{ij}A_{jk} > 0$. We found that, for a multiplicity of metrics, the differences between feasible subsets of species and unfeasible subsets (negative abundances) where small and, even if there could be certain statistical differences or a diffuse pattern, visual inspection of violin plots did not yield a notable signature that allowed us to hypothesize that a given metric is explanatory for EC specifically, beyond the diffuse metrics linked to feasibility studied in (Aguadé-Gorgorió and Kefi, 2024; Barbier *et al.*, 2021). Some of the studied metrics with weak significant differences are (not shown and not all discussed in detail in this Supplementary Material): (1) the aggregate statistics of interactions $\mu^*, \sigma^*$, where relevant differences emerge for stable states (see section below) but not do not differentiate feasible from unfeasible matrices, (2) the fraction of positive net effects (see section below), (3) skewness and kurtosis of interactions, although feasible states appear to have a slightly more negative kurtosis than unfeasible ones, (4) properties of columns in $A^*$ such as column sum heterogeneity, (5) Collectivity and condition number of feasible and unfeasible matrices, (6) presence of motifs where $A^*_{ij} < -1$ is accompanied by $A^*_{ij}A^*_{jk} > 0$ and more general correlations between the first two elements of the Neumann series $A^*$ and $(A^*)^2$, etc.

Among these and many other studied metrics that did not seem to explain why some subsets with EC were feasible and others were not, metrics of type (6) give an interesting intuition: there seemed to be no microscopic motifs by which a targeted species would be rescued by e.g. a chain of length two. We knew from three-species models (see Section I.B.2) that such motifs of positive indirect effects could explain competitive/emergent coexistence, but we could not find any variant of these in our communities of 4, 5 or more species. This prompted the realization that our reductionist search for microscopic motifs was being blurred by longer chains of effects, which is at the core of the present work: a high collectivity metric indicates that any short chain of direct or indirect effects carries little or no information about the final outcome of the community. When the spectral radius of a matrix is large, the stable abundance of a species cannot be measured in terms of sums of chains of indirect effects. This explained why we hardly could not find any pattern in $A^*$ or $(A^*)^2$ that explained the positive abundances found in the row sums of $(I - A^*)^{-1}$: At high collectivity, $A^*$ and $(I - A^*)^{-1}$ are uncorrelated, and the typical "enemy of my enemy is my friend" motif that we expect from low-dimensional systems disappears, and the same goes with rock-paper-scissors signatures and so on. This is in fact why emergent coexistence is an emergent phenomena: if interactions are strong, long chains of pairwise interactions become so heavy that direct or second-order indirect interactions do not seem to play any role at all. We then measured collectivity in systems with EC to find that most systems harbored collectivity higher than 1, and even those with collectivity smaller than 1 would have e.g. $\phi \approx 0.7$, so that very long chains of interactions play a role in the dynamics and signatures in $A^*$ or $(A^*)^2$ alone are insufficient to predict final abundances.

Additionally to the subtle coexistence signatures found in (Barbier *et al.*, 2021), we highlight here one signature of the interactions between coexisting species that we were able to find among the many different measurements. In brief, the row sums of $A^*$, $\sum_j A^*_{ij}$, are more homogeneous than what one would expect at random, a signature that is not true with the row sums of unfeasible matrices or the column sums of feasible matrices (Fig. 5). To measure this, we measure the standard deviation of $\sum_j A^*_{ij}$ across all columns $i$, and divide it by the average standard deviation of 100 different randomizations of $A^*$. A value of 1 indicates that row sums are not more homogeneous than the random expectation, whereas a value smaller than 1 indicates that row sums are more homogeneous than what one would expect by simply locating interaction strenghts at random, as is done in the original pool of interactions $A$. The observation shown in figure 5 indicates that feasible communities with EC are characterized by the coexistence of species that have similar $\sum_j A^*_{ij}$ values. This has an explicit meaning in the GLV model as $\sum_j A^*_{ij}$ is the total perceived competition if all species had the same abundance. Homogeneity of row sums means that coexisting species must perceive competition similarly, and there cannot be a species that receives much more competition than others. Interestingly, the same is not necessarily true for column sum heterogeneity: we find that column sums are not more homogeneous than expected at random (not shown). This means that the impact that species $j$ has on all other species, $\sum_i A^*_{ij}$, is not necessarily homogeneous. All species must perceive a similar competition, $\sum_j A^*_{ij}$, but it seems irrelevant if this competition is exerted by all species in a similar way or else there are some stronger competitors than others leading to no significant homogeneity in $\sum_i A^*_{ij}$.

#### 2. All-to-all competition but positive net effects

Once we have analysed the properties of $A^*$ that are (and are not) related to the presence of EC and the overall survival of strong competitors, it is also interesting to observe the properties of $(I - A^*)^{-1}$, the matrix capturing the net effects between species. If one recalls the Neumann series,

$$(I - A)^{-1} = 1 + A + A^2 + A^3 + ... \tag{61}$$

which, as discussed in the main text, implies that the stable abundance of a given species is

$$x^*_i = 1 + \sum_j A^*_{ij} + \sum_{j,k} A^*_{ik} A^*_{kj} + \sum_{j,k,l} A^*_{ik} A^*_{kl} A^*_{lj} + ... \tag{62}$$

it appears clear that, even in systems where species interactions are purely competitive, net effects are not necessarily all negative, and multiple positive indirect effects are likely to emerge. Here it becomes useful to measure the so-called Positive Feedback Index introduced in (Liautaud *et al.*, 2019), that counts the fraction of elements in $(I - A^*)^{-1}$ that are positive. We believe it is better avoiding to call these elements *Feedbacks*, as there is no constraint by which these elements must be closed loops so that a species change *feeds back* on its own abundance. To avoid confusion between loops, feedbacks and interaction chains, we simply call this index the fraction of positive net effects. In (Liautaud *et al.*, 2019) the authors found that, in the assumption of purely symmetric interactions that limits the possible outcomes of the system (see Section II.A.4), the fraction of positive net effects could be around 15%. Yet, in our model and for random and uncorrelated interactions, we find that once interactions are strong enough and approach or fall into the EC regime, the fraction of positive net effects easily reaches 50% (Fig. 6). Even more, for weak competition, coexistence states rapidly harbor a large fraction of positive net effects, even before EC is in place. This means that a community will rapidly transition from surviving via self-regulation to surviving via positive direct and indirect interactions. The limit case scenario of 50% net effects can also be seen in Figure 3B in the main text: even when most interactions are competitive (very few $A^*_{ij} > 0$), around half of the net effects are positive (many $(I - A^*)^{-1}_{ij} > 0$). This has also important implications regarding the correlation between the elements of $A^*$ and the elements of $(I - A^*)^{-1}$ (see section below). We are unsure of the explanation by which the fraction of positive net effects measured in (Liautaud *et al.*, 2019) is much lower than in our results. We hypothesize that it could be because their model considers symmetric interactions as well as no species extinctions, which increase the presence of permanent fluctuations and eradicate the likelihood of many stable states with high diversity (see Section II.E and (Aguadé-Gorgorió and Kefi, 2024)). However, we were unable to replicate their results observing a maximum fraction of positive net effects around 15%. All in all, the fact that the fraction of positive net effects can be as high as 50% links emergent coexistence with the weight of indirect effects as captured by the collectivity parameter: if interactions are very weak, long chains of indirect effects will play a weak role, as $A^k$ tends to zero as $k$ increases and species abundances are determined by self-regulation or, at most, by direct interactions. Yet, once interactions are strong enough, long and heavy chains of indirect effects role over direct interactions, and many positive net effects emerge. This leads to unexpected survival (emergent coexistence) of many species pairs that, when isolated from the community and these indirect effects, do not coexist due to strong competition.

#### 3. Indirect effects and an estimate for minimal and maximal collectivity

In Section II.H we have explained the numerical simulations that lead to figure 3A in the main text, for which we sample stable states from the GLV model with different diversity and measure the collectivity metric for those with EC in purple and those without EC in gray. It is useful to have an analytical estimate for how collectivity increases with species diversity, as collectivity gives a direct proxy for the degree to which direct species interactions provide information on community-level net effects.

To do so, we take advantage of the work developed in (Allesina and Tang, 2012) that studies the spectral radius of a random matrix from information on its statistical properties, as discussed in the supplementary material of (Zelnik *et al.*, 2024). These results are used to understand how stability conditions relate to interaction properties of the Jacobian matrix $J$ in (Allesina and Tang, 2012), but we will use the same results to infer the spectral radius of the interaction matrix $A^*$. This does not carry direct information about linear stability (which we review below), but on the relative weight of indirect effects and the degree of collective integration of a community (Zelnik *et al.*, 2024).

Given once again $\mu^*$ and $\sigma^*$ as the mean and standard deviation of the interaction strengths $A^*_{ij}$ between $S^*$ species that survive and coexist stably from the original pool of $S$ species with interaction matrix $A$, random matrix theory predicts a spectral radius of (see e.g. (Allesina and Tang, 2012; Zelnik *et al.*, 2024)):

$$\phi_{\text{rmt}} = \max\{(S^* - 1)C|\mu^*|, (1 + |\gamma^*|)\sigma^* \sqrt{C(S^* - 1)}\} \tag{63}$$

where connectivity $C = 1$ in the main text (see Section I.D for a discussion on variable connectivity) and $|\gamma|$ measures the absolute reciprocity of interactions ($\gamma = \text{corr}(A_{ij}, A_{ji})$, see Section S3.1 of (Liautaud *et al.*, 2019)), which provides a proxy for the degree of symmetry (mostly competitive or mostly cooperative interactions, $\gamma \to 1$) or antisymmetry (mostly predator-prey interactions, $\gamma \to -1$), which we can compute numerically from $A^*$.

The minimal collectivity that we can find in the GLV model equals zero, as there can be systems with no species interactions, so that $\mu^* = 0$ and $\sigma^* = 0$. Yet, these systems cannot generate EC as there are no exclusionary interactions. Instead, to measure collectivity for systems with EC, we must recall once again that we can predict the maximum $\mu^*$ and $\sigma^*$ values that a state with $S^*$ species can harbor (Aguadé-Gorgorió and Kefi, 2024). In fact, we know that larger systems allow for smaller $\mu^*$ values. This provides a key intuition: even if one considers $\phi_{\text{rmt}} \sim (S^* - 1)C|\mu^*|$, the maximum competition that a community can sustain $|\mu^*|$ decreases as $S^*$ increases (Section II.C). This explains why the largest collectivity that we observe is not necessarily exactly linear with $S^*$: increasing the number of species increases the total number and length of chains of indirect effects, but decreases the strength of interactions that a community can sustain and hence the weight of these indirect effects. However, as seen in figure 3 of the main text, the second effect is negligible compared to the first, which can be explained by seeing that we find that maximum collectivity is well estimated by

$$\max(\phi_{\text{EC}}) \approx (S^* - 1)\max(|\mu^*|) \approx (S^* - 1)\left(1 - \frac{(S^*)^{1.08}}{14.118}\min(\sigma^*)\right), \tag{64}$$

where, once again, the smallest $\sigma^*$ state that a community with $S^*$ species emerging from an original pool $A$ has a closed estimate discussed in Section II.C. As we discuss below, this correction for small random matrices is valid for communities of moderate size, but fails when diversity is as small as 3 or 4 species, and Section II.C explains why collectivity can be even higher than our estimate for such small diversities. In systems where competition strength is close to $\mu = -1$ (the neutral case or Hubbell point discussed in (Kessler and Shnerb, 2015)), Emergent Coexistence is possible with $\sigma^* \to 0$ (Figure 2A in the main text). Under this scenario, $\phi$ increases linearly with $S^*$.

Additionally, we provide an estimate for the smallest possible collectivity that a community with EC can sustain. We hypothesize that the smallest collectivity is found for communities with the weakest mean competition possible. For small communities, this is simply zero: even if EC does not exactly happen for $\mu = 0$, $\mu^*$ can be zero as it is only the mean of a small subset sampled from $A$. Yet, as the size of a community increases, we previously showed how the statistics of $A^*$ tend towards those of $A$ (Aguadé-Gorgorió and Kefi, 2024), and the community with EC and weakest . In this context, it suffices to find that the weakest competition possible for EC ($\min(|\mu|)$, Section II.C) is approximately 0.071, so that $\min(\phi_{\text{EC}}) \approx (S^* - 1)0.071$. However, because we do not know of an analytical equation for the transition curve towards unbounded growth (Figure 2A in the main text, vertical dashed curve), we cannot give an analytical expression for where the minimal interaction strength line (Figure 2A in the main text, red line) crosses this curve, and we can only visually estimate that is close to 0.071. Nevertheless, this numerical approximation provides a good bound for the minimal spectral radius of a stable community with Emergent Coexistence (Figure 3A in the main text, bottom red dashed line).

As discussed in the main text, the role of collectivity can also be understood in terms of the correlation between direct and net interactions (Fig. 3B in the main text). In figure 7 we show how, for communities with weak interactions (in gray), direct and net interactions correlate quite well, as there are not many indirect effects. Yet, for communities with strong interactions and EC, direct and net interactions can appear uncorrelated (the central link between $\phi$ and EC discussed in the main text) or even anticorrelated in few-species systems, where a direct competitor is also a strong cooperator by competing with other species in the community. As the number of coexisting species decreases, $A^*$ and $(I - A^*)^{-1}$ tend towards no correlation at all: indirect effects are so strong, that direct interactions become a negligible piece in the community puzzle.

#### 4. An estimate for minimal condition number $\kappa$

If direct interactions are insufficient to predict net effects, the solution is to invert the interaction matrix $A^*$ and estimate $(I - A^*)^{-1}$ numerically. In many fields dealing with high-dimensional systems, the problem of inverting large matrices with minimal error is a classical challenge (Trefethen and Bau, 2022). A metric that captures the precision of matrix inversion is the *condition number* $\kappa$ ((Edelman, 1988; Trefethen and Bau, 2022)). Very generally, $\kappa(I - A^*) = 1$ indicates that an error in estimating the values of $A^*$ will propagate to a similar error in $(I - A^*)^{-1}$. A larger value of $\kappa(I - A^*)$ indicates that small errors in $A^*$ can lead to large errors in $(I - A^*)^{-1}$, blurring our capacity to predict species abundances accurately ((Trefethen and Bau, 2022)) Despite its critical role in understanding predictability, the condition number of ecological interaction matrices has received little attention (but see (Gilpin, 2024) for a very recent work and (Graham, 2003; Strydom *et al.*, 2021) for discussions in adjacent problems).

The *condition number* of a matrix is a key measure of numerical stability, particularly in the context of solving linear systems or inverting matrices. Given that the equilibrium abundances solution for the GLV model is

$$(I - A)\boldsymbol{x} = 1, \tag{65}$$

and so

$$\boldsymbol{x}^* = (I - A^*)^{-1}1, \tag{66}$$

it is key to understand how errors in $A^*$ propagate to the inverse matrix. For such a matrix $(I - A^*)$, its condition number with respect to the 2-norm is defined as:

$$\kappa((I - A^*)) = \frac{s_{\max}}{s_{\min}} \tag{67}$$

where $s_{\max}$ and $s_{\min}$ are the largest and smallest singular values of $(I - A^*)$, respectively. The meaning of these values is discussed very briefly below, and in much more detail in (Edelman, 1988; Trefethen and Bau, 2022).

Intuitively, the condition number quantifies how sensitive the solution $\boldsymbol{x}^*$ of the linear system $(I - A)\boldsymbol{x} = 1$ is to small perturbations in the matrix $(I - A^*)$ (and hence, $A^*$) itself. A high condition number implies that even small errors in data or computation can lead to large errors in the solution, indicating that the matrix is *ill-conditioned.* Conversely, a condition number close to 1 implies that the matrix is *well-conditioned* and stable under numerical operations.

For symmetric positive definite matrices, the singular values coincide with the absolute values of the eigenvalues, and the condition number simplifies to the ratio of the largest to smallest eigenvalue. But, as seen in Section II.A.4; symmetric matrices are precisely the ones that carry less EC states, and we are mostly interested in the asymmetric, uncorrelated case.

To do so, we take advantage of the large amount of work regarding the singular values of random matrices, while reminding that this is by no means a work of fundamental mathematics, for which we refer the reader to works such as (Edelman, 1988; Trefethen and Bau, 2022). What follows is only intended to provide a qualitative understanding for a general reader.

Our main task is to understand how the maximum and minimum singular values of a random matrix, $s_{\max}$ and $s_{\min}$, are related to $S^*$, the dimension of the matrix, and $\mu^*$ and $\sigma^*$. Hence, we will have for now to forget here about the diagonal entries of the matrix, which are no longer random, and link our predictions with the all-random case.

If the matrix $(I - A^*)$ has i.i.d. entries from $\mathcal{N}(0, \sigma^*)$, random matrix theory provides the following asymptotic estimates for large $S^*$ (Edelman, 1988; Trefethen and Bau, 2022):

$$s_{\max} \approx 2\sigma^* \sqrt{S^*} \tag{68}$$

$$s_{\min} \approx \frac{\sigma^*}{\sqrt{S^*}} \tag{69}$$

Now, to find the condition number of systems with EC one would want to compute $\kappa((I-A^*)) \approx s_{\max}/s_{\min}$. Yet, we have to remember that we are studying not a single EC state with given $\mu^*$ and $\sigma^*$ values, but rather an ensemble of many states with variable $\mu^*$ and $\sigma^*$ values. We do not want to give an exact approximation, but rather a numerical bound for how $\kappa$ changes across all these systems.

Let us first study the largest singular value across our ensemble of states, $s_{\max}$. It is useful to recall that $s_{\max}$ provides an intuition of the maximum stretching that our matrix of interactions can produce to a given vector of abundances (Edelman, 1988; Trefethen and Bau, 2022). From all the EC states that we can observe within the domain of figure 2A (within which our simulations are restricted for illustrative purposes), the state that generates the most stretching is the one with the most positive interactions (weaker competition, $\max(\mu^*)$) and the largest possible $\sigma^*$. This state will provide a direction in which stretching is maximum (Fig. 8). Instead, the smallest $s_{\max}$ value will be found for a community that is able to do the least possible stretching, so that interactions are the most competitive (strong competition, $\min(\mu^*)$), and heterogeneity is also small to ensure no cooperators are present (remember that we are searching for the extreme bounds, not exact solutions). In this context, and as seen for figure 8A, the largest singular values across the different states are bound within

$$s_{\max} \in \left(|\min(\mu^*)| + 2\min(\sigma^*)\sqrt{S^*}, |\max(\mu^*)| + 2\max(\sigma^*)\sqrt{S^*}\right) \tag{70}$$

where the largest and smallest $\mu^*$ and $\sigma^*$ values for which we find EC can be simply recorded during a simulation or can be inferred numerically using the analytical derivations discussed in Section II.C.

If we look at figure 8B,C or figure 3C of the main text, we realize that the largest $\kappa$ number will be problematic to compute: because $s_{\min}$ can be arbitrarily low, the largest condition number can be extremely large and fluctuate a lot. As discussed in the main text, this has deep implications for ecological predictability. Let us at least find what is the smallest possible $\kappa$ number for communities with EC. That is, what is the best-case scenario for predictability of equilibrium abundances in these communities.

$$\min(\kappa((I-A^*))) \approx \frac{|\min(\mu^*)| + 2\min(\sigma^*)\sqrt{S^*}}{\max(s_{\min})}. \tag{71}$$

We need therefore to estimate what is the largest possible value for the smallest singular value, $\max(s_{\min})$. In figure 8A we can see a relatively good estimate

$$s_{\min} \in \left(0, \frac{\max(\sigma^*)}{\sqrt{S^*}}\right) \tag{72}$$

The largest possible value for the smallest singular value corresponds to EC states with highest heterogeneity: they are the ones for which the minimal stretching can be larger (Fig. 8A).

This provides us with a first rough estimate for the minimal condition number for communities with EC, which roughly indicates that the best-case scenario for inverse-matrix predictability will behave as

$$\min(\kappa((I - A^*))) \approx \frac{|\min(\mu^*)| + 2\min(\sigma^*)\sqrt{S^*}}{\max(\sigma^*)/\sqrt{S^*}}. \tag{73}$$

Because $\min(\sigma^*)$ can be very close to zero for some small communities with homogeneous interactions (Section I.J, (Aguadé-Gorgorió and Kefi, 2024)), the typical observation of $\min(\kappa((I-A^*))) \sim S^*$ here takes a more nuanced form, where these small communities with relatively homogeneous interactions could go as low as

$$\min(\kappa((I - A^*))) \approx \frac{|\min(\mu^*)|}{\max(\sigma^*)}\sqrt{S^*}. \tag{74}$$

For larger communities where interactions cannot be so homogeneous, we recover the result above, where $\min(\kappa((I - A^*))) \sim S^*$. Unless interactions are extremely weak, a system involving tens to hundreds of species will have, in the best-case scenario, a condition number scaling linearly with diversity. As discussed throughout the text, this poses important numerical limitations in the precision of solving equilibrium abundances from imprecise measurements of direct interactions

In conclusion, communities with EC have a large condition number due to the high strength and heterogeneity of interactions between many species (Purple dots in fig. 8C). The lower analytical bound indicates that the minimal condition number increases with species diversity as $\kappa(I - A^*) \sim S^*$ (Fig. 8C red dashed line, (Edelman, 1988)). Most interaction matrices, however, have $\kappa$ values much larger than that, and only communities with weak and homogeneous interactions have a condition number close to 1 (Gray dots in fig. 8C).

As first additional note on predictability and numerical precision, there might be instances where errors in the measurement of interactions, $A + \epsilon$, behave better (propagate less) by integrating the dynamics with a precise algorithm (section I.E.1) rather than inverting the matrix and calculating net effects (sections I.E.2 or I.I). Because of the qualitative and generalist nature of our work, we leave the accurate comparison of these two methods for future work and refer the reader to e.g. (Gilpin, 2024) where temporal unpredictability and condition number are studied in detail in ecological models.

As a second additional note, it is worth discussing the meaning of numerical instability (high $\kappa$) in the empirical properties of communities as those observed in (Chang *et al.*, 2023). Here is where commenting the methods of *structural stability* becomes particularly relevant (Cenci and Saavedra, 2018; Rohr *et al.*, 2014; Saavedra *et al.*, 2017). Very generally, structural stability asks if a given community state will remain the same (the same species coexisting) under a given change in the interactions or carrying capacities of the system. Hence, the structural stability metric of a community gives a quantitative notion of the volume in the parameter space (typically, of growth rates) within which such combination of species will be observed, and crossing this volume implies that other species might invade or some of the present ones go extinct. In this context, a large $\kappa$ value seems to be somehow related to a small structural stability, meaning that small changes in $A^*$ can lead to great changes in $(I - A^*)^{-1}$ which, in turn, could be related to changes in the composition of the community. While we do not have a mathematical explanation that brings together condition numbers and structural stability, we believe it could be interesting to explore the structural stability of EC communities, their fragility to parameter changes, and the overall link of these with the condition number and predictability. Finally, it is worth commenting that the structural stability approach might also prove valuable for studying EC. In it, one could ask the following question: under which condition the structural stability volume (the space of possible coexistence conditions) of the whole community is larger or non-overlapping with the volume predicted by its species pairs?

#### 5. Empirical applications of using $\phi$ and $k$ to infer predictability

In Section I.I we have proposed two numerical tests for the role of $\phi$ and $k$ in different experimental setups.

To study the role of $\phi$, we try to assemble communities of species that coexist by pairs, and ask if they coexist in the larger community. To study the role of $k$, we try to predict community composition given a matrix of interactions with a certain error in the measurement of interaction strengths, $A^*_{ij} + \epsilon$. In Figure 9 we summarize our findings. In parallel, we refer the reader to very recent works such as (Arya *et al.*, 2025) and (Solé *et al.*, 2024) where the possibilities, fields of application and importance of assembling microbial consortia are discussed in detail.

Regarding the first test, one would expect that communities with $S^* = 2$ species will always past the test: selecting two species that coexist by pairs and assembling them together will obviously yield coexistence. We know from (Friedman *et al.*, 2017) and (Lee *et al.*, 2023) that this test will also work in most scenarios of three species, but will start to fail as we group together more and more species. In figure 9A we show how the collectivity parameter $A^*$ of the species interaction matrix provides an explicit metric for the success or failure of our assembly test. We find that, for very low collectivity $\phi \to 0$, the test will most often succeed, related to communities with very weak indirect effects and few species. Yet, for $\phi \approx 1$, half of the tests will already fail, meaning that a community of coexisting pairs will not result in all species coexisting. Once $\phi \gtrsim 3$, most communities will fail at coexisting, meaning that information of direct interactions will not allow us to predict the coexistence in a complete community. This result harbors an additional important property: here we are assembling communities of coexisting pairs. This means that there are no competitive interactions stronger than $A^*_{ij} = -1$, and we are not necessarily in the domain of EC and strong competition, where we know that chains of indirect effects will be strong. Even before this domain, and for competition weaker than $A^*_{ij} = -1$, communities with large collectivity values will already have strong indirect effects, leading to a failure in the reductionist approach of assembling communities by grouping species that coexist in co-culture. In fact, the test of figure 9A indicates the contrary: assembling many coexisting pairs will most often result in a failed community, whereas the assembly of strong competitors can sometimes yield coexisting communities as shown in the main text. Emergent coexistence and the collectivity parameter, however, imply that we cannot know in principle which of these strong competitors should we culture in groups to ensure a stable community.

In the second test, we predict the coexistence of a community from imprecise measurements, $A^*_{ij} + \epsilon$, and ask if the real dynamics generated from $A^*_{ij}$ also result in community coexistence, or else if the error has propagated into the inverse as much as to predict a false coexistence result. We see in figure 9B that the condition number of the interaction matrix, $k(A^*)$, provides a strong metric to understand the propagation of this error. Fixing $\epsilon \sim \mathcal{N}(\mu = 0, \sigma = 0.1)$, we see that systems with $k \approx 1$ will predict coexistence with almost total precision. Yet, as $k$ increases, this precision drops very abruptly, so that for $k \approx 20$, about 90% of our tests will fail, and real communities will not coexist even if we predicted so. Of course, we find that the decay of predictability with $k$ depends on the error, and for a much more precise $\sigma = 0.01$ we find that communities with $k \approx 20$ yield a success of 50%, compared to the 10% success of figure 9B. All in all, this simple test provides a clear demonstration of how the condition number of an interaction matrix provides a valuable tool to assess the predictability of community composition given a certain error $\epsilon$ in our measurements of pairwise interactions. Experimental tests studying communities with strong and heterogeneous interactions would benefit from measuring the condition number $k$ as a means to infer the validity and error propagation in predicting the likelihood of species coexistence.

#### C. Feedback loops and stability

In Section I.J we have introduced May's complexity-stability limit, also discussed in more detail in (Aguadé-Gorgorió and Kefi, 2024; Bunin, 2017; Mallmin *et al.*, 2024; May, 1972). Very briefly, this limit imposes a maximum combination of interaction strength, interaction heterogeneity, species diversity and network connectivity (combined, the *Complexity* of a community, (Zelnik *et al.*, 2024)) beyond which a Jacobian matrix is almost sure to harbor positive eigenvalues and hence becomes linearly unstable. Because this condition is known to operate for large matrices ($S^* \approx 50$ for $C = 1$, (Aguadé-Gorgorió and Kefi, 2024)), we have also introduced a small-matrix approximation, for which we found in a previous work that communities with a moderate number of species ($S^* \approx 10$) have a slightly larger complexity-stability limit and can hold a larger complexity, which, in our model, translates as communities with higher diversity and competition strength remaining stable. Finally, we have also introduced the Routh-Hurwitz criteria, which are not a statistical approximation from random matrix theory, but rather an exact set of criteria that the coefficients of the characteristic polynomial of the Jacobian matrix must fulfill for the Jacobian to have negative eigenvalues (Dambacher *et al.*, 2003; Levins, 1974; Neutel *et al.*, 2002).

In figure 10 we plot a large set of $10^7$ stable states sampled from the GLV model in the domain of figure 2A of the main text with variable diversities (see Section I.E.2 on how we sample stable states from the GLV model). We paint in gray those states that do not have any EC, which in the end will result in states with weak competition strength, and in purple those states which have EC, meaning they harbor one or more excluding pairs with $A^*_{ij}$ or $A^*_{ji}$ smaller than -1. We know that these states are linearly stable, because we have obtained the eigenvalues of their Jacobian matrices evaluated at each equilibrium. We want to observe how the different conditions on stability (May's bound, the small matrix correction and the second determinant of the Routh-Hurwitz criteria) apply on these states, with a particular focus on states that have a moderate diversity similar to that observed in laboratory experiments of microbial community assembly (Chang *et al.*, 2023; Friedman *et al.*, 2017; Venturelli *et al.*, 2018).

As discussed in the main text, we find that there exist a lot of small-diversity states that have stronger competition strength than what the Random Matrix Theory expectation would indicate (Fig. 10A). This translates into $|\mu^*|$ being larger (more competitive) than $|\mu^c|$, the maximum competition discussed in section I.J and II.A. Briefly put, this simply means that the limits of random matrix theory discussed by May in (May, 1972) do not operate for communities of such a small size. This is somehow an expected result, as the overall complexity-stability debate asks for the conditions under which very diverse ecosystems (think about coral reefs or tropical forests with hundreds of species) would remain stable to perturbations. It does not apply to asking how can a community with three or six or eight species remain stable, mostly because these matrices are so small that the fact that the original pool $A$ is a large and random matrix does not imply that the interaction matrices $A^*$ of these small communities are exactly random anymore. In figure 10B we can see that, as we discussed in detail in our previous publication (Aguadé-Gorgorió and Kefi, 2024), the correction for small matrices predicts much better which communities remain stable and which don't, and now only a few and very small communities are able to overcome the threshold of maximum competition. But once again, it is clear that this boundary is not sharp nor exact, because it also stems from statistical aggregates that only apply for large enough communities where microscopic structures do not play a major role.

This can be seen in more detail by looking at figure 10C, where we plot the sign of the second determinant from the Routh-Hurwitz criteria (Section I.K). Albeit larger determinants result in much more complex stability conditions, the second determinant can be roughly understood as a condition by which feedback loops of order three in the Jacobian (not in the interaction matrix) must be weaker than combinations of order one and two (Section I.K, (Levins, 1974)). Consistent with (Neutel *et al.*, 2002), our preliminary simulations show that once this second determinant is fulfilled all others are mostly fulfilled as well (not shown), so that we focus on this determinant that harbors a simpler explanation in terms of feedbacks. We now see that all states, both with and without emergent coexistence, must fulfill this condition. This signature was previously discussed in detail in (Dambacher *et al.*, 2003; Levins, 1974) and also found in experimental estimates of the Jacobian matrix of food webs, were weak interactions had been found to participate in lowering the weight of these long loops and hence favouring linear stability (Neutel *et al.*, 2002). In short, our results imply that, while random matrix approximations might operate in communities of a certain species richness, linear stability in smaller communities is ensured by a more nuanced community structure, by which interaction strengths of the surviving species follow a pattern so that long loops, even if competition can be strong, do not counterbalance the strength of self-regulation or two-species feedbacks. In the style of (Barbier *et al.*, 2021), that identified fingerprints of coexistence in the interactions between surviving species ($A^*$), it would be interesting to study if systems for which we have estimates of $A^*$ and $x^*$ (and hence, we can build $J^*$) fulfill the RH criteria as previously shown in (Neutel *et al.*, 2002). However, both in-depth analysis of linear stability and the use of empirical estimates $A^*$ matrices are beyond the scope of this qualitative study.

Another interesting way to visualize these results is by plotting together the distance to May's threshold and the sign of the second Routh-Hurwitz determinant for stable (purple) and unstable (gray) subsets of species with size between $S^* = 3$ and $S^* = 15$ (Fig. 11). We can see how there are many stable communities with EC that do not fulfill the random matrix prediction and have stronger competition than expected (x-axis), but still always fulfill the RH condition on balanced feedback loops (y-axis).

#### D. Intransitivity and emergent coexistence

Coexistence under strong competition has often been linked to intransitivity in theoretical models (Gallien *et al.*, 2017; Laird and Schamp, 2006; Levine *et al.*, 2017). Intransitivity can be defined by a lack of hierarchy in the competition matrix: in a three-species model, competitive dominance can be counterbalanced by a rock-paper-scissors loop, where species A outcompetes species B, B outcompetes C but C outcompetes A (Gilpin, 1975; May and Leonard, 1975). More generally, an intransitive network can be characterized by the presence of low rank exclusions, where a species with lower competitive rank (e.g. number of wins minus loses, Section I.L) is able to outcompete a species with higher rank (Chang *et al.*, 2023; Higgins *et al.*, 2017; Laird and Schamp, 2006). Here we analyse and extend the numerical results presented in the main text regarding community intransitivity.

##### 1. Low rank exclusions require 4 exclusions, and even then are rare

In Figure 4A,B of the main text we find that the fraction of low rank exclusions in stable communities with EC is, on average, about $2-3\%$ of the total exclusions in communities of more than about 5 or 6 species, consistent with experimental observations ((Chang *et al.*, 2023; Higgins *et al.*, 2017)). Yet, one of the reasons behind this low value in simulations and experiments is that a single low rank exclusion requires a minimum of 4 exclusionary interactions, which is clearly seen in figure 12. In figure **??** we show the LRE metrics for both the uncorrected version as well as the corrected version of LRE shown in the main text, in which we correct for this issue by measuring low rank exclusions only in communities with four or more exclusionary interactions. In any case, our results show that while some lower ranked species exclude higher ranked ones in systems of moderate size (Fig. 13A), this signature disappears if we do not consider the 4-exclusions requirement: there will be so many communities with 3 or less exclusionary interactions, that it might seem that LRE are absent (Fig. 13A). What is absent is actually the 4-exclusions motif. As an example, we can see how LRE are not possible in systems with only 3 species, showcasing the imprecise use of this metric to assess intransitivity: a community with 3 species can be perfectly intransitive due to a RPS loop, but have a null LRE metric as there will not be the necessary rank differences to observe LRE as proposed in figure 12. Both for the unconditioned case and for the conditioned case (Fig. 13A,B), we can see that large communities very rarely harbor LRE's, consistent with empirical observations (Chang *et al.*, 2023; Higgins *et al.*, 2017). We understand that this condition was not implemented in (Chang *et al.*, 2023; Higgins *et al.*, 2017), so that their results are better described by the extremely low LRE fractions observed in (Fig. 13A). Because we believe the conditioned case describes better the real decay of intransitivity, we have presented the conditioned version in the main text.

#### 2. Rock-Paper-Scissors loops are equivalent to a random expectation

In figure 4C of the main text we plot the fraction of rock-paper-scissors triplets in stable communities with EC. If rock-paper-scissors are necessary for coexistence, a large proportion of triplets should have this motif. The classical example is that of communities with 3 species: if all three species are connected by excluding competition, the only way to avoid one species dominating the others is a rock-paper-scissors loop (Fig. **??**C, dotted circle, (Gilpin, 1975; May and Leonard, 1975)). Yet, for larger communities, we find again that the presence of rock-paper-scissors motifs decreases with species richness. It is important that we are measuring the fraction of triplets that are RPS in communities where there are triplets, so the problem of the LRE metric described above is not present here: in the RPS metric the necessary condition (similar to the 4-exclusions condition above) is that there is a triplet connected by competition, which is how we measure the metric by definition (Section I.L).

We find that the fraction of RPS triplets approaches the null expectation where only 1 in 4 triplets will be RPS, found after shuffling the elements of $A^*$ (gray, Methods, (Chang *et al.*, 2023)). This null expectation is easy to understand following the same rationale as that of figure 12: if one tries drawing all possible triplets connected by pairwise exclusion (see Section I.L for definitions), it will find 2 RPS loops (the clockwise and counterclockwise cases) and 6 one-species-dominates motifs. As shown in figure 4C with the dark dashed line and the gray dots of the shuffled expectation, 1/4 is indeed the likelihood that a triplet is RPS if interactions are drawn at random or, at least, randomized. The implications of this are worth discussing: in the last section of (Chang *et al.*, 2023), the authors ask for the probability that, of 77 observed triplets, none is RPS.

They say that this probability is $P = 0.25^{77} = 4.4 \times 10^{-47}$, which of course an extremely low value. Yet, we believe there is a mistake in this calculation. The probability that a triplet is *not* RPS is $P = 0.75$. Then, the probability that, of 77 triplets, none is RPS is $P = 0.75^{77} = 2.4 \times 10^{-10}$. While this probability is still extremely low and requires that additional mechanisms explain the absence of triplets in laboratory observations of EC, it is 37 orders of magnitude larger than the probability proposed in (Chang *et al.*, 2023). In any case, the consistent observations of EC without RPS triplets require some additional explanation. In the main text and Sections I.M and II.A.4 we propose that this is related to the presence of hierarchical competition.

#### 3. Equivalent results with the Laird-Schamp index

In Section I.L we have presented the study by (Laird and Schamp, 2006), where they propose to measure intransitivity in coexist-exclude tournament networks as the standard deviation in the vector of win counts for a community. A high standard deviation indicates that there might be some species that exclude a lot while others do not, indicative of a highly transitive or hierarchical structure. As we discuss in detail in Section I.L, their proposed metric of $\sigma_{obs} - \sigma_{min}/\sigma_{max} - \sigma_{min}$ has a caveat for communities a small number of wins: in communities with only one exclusionary pair, all vectors would be the same (1 species has 1 win, all other species have zero wins), and so $\sigma_{max}$ and $\sigma_{min}$ would be the same and the metric would diverge, making the index ill-defined for scenarios with few exclusions. In figure 14 we measure the Laird-Schamp heterogeneity index (where possible) as well as the proposed variation of $\sigma_{obs}/\sigma_{max}$, which does not have the well-defined bottom state of totally intransitive competition equals zero as the original metric. In any case, what we are interested in and what we observe in figure 14 is that intransitivity decays as species diversity increases. This provides an intuition that is consistent with that proposed in the main text and in the sections above: intransitive motifs are necessary to ensure coexistence in few-species systems, yet in more diverse communities these motifs get blurred by the very many indirect interaction chains in place.

All in all, intransitive structures only seem essential for species coexistence in very small communities, where they are necessary to avoid dominance of the winner species (Gilpin, 1975; May and Leonard, 1975). In more diverse communities, there exist so many interaction chains that can rescue species from extinction, that intransitivity might be present, but is no longer required for coexistence. Low or high intransitivity, therefore, might be a byproduct of the structure of interactions, potentially explaining seemingly contradictory results across empirical data (Godoy *et al.*, 2017; Soliveres *et al.*, 2015). In limit-case scenarios of nested or single-resource competition discussed in the main text and Sections I.M and II.A.4, intransitive signatures can even disappear.

#### E. On the role of migration and reinvasions

A principal property of the GLV model studied in this work is the absence of species migration. We aim at studying a set of closed experimental ecosystems for which there is no evidence that extinct species can be reinoculated into the assembled community (Chang *et al.*, 2023; Friedman *et al.*, 2017; Venturelli *et al.*, 2018). However, as we discussed in detail in a previous work, the possibility that species can migrate back into the community even if temporarily extinct changes some of the properties of the model (see (Aguadé-Gorgorió and Kefi, 2024) and references therein). This is typically expressed by adding a small migration rate $m$ to the original model

$$\frac{dx_i}{dt} = x_i \left( 1 - x_i + \sum_{j \neq i}^{S} A_{ij} x_j \right) + m. \tag{75}$$

In the simulations, a qualitatively equivalent mechanism instead of adding an external constant migration is to ensure that species cannot go extinct, by adding a minimal abundance below which a species cannot keep decreasing. Mathematically, we had tested subsets of the community to see if they lead to states with positive abundances and linear stability. Adding migration requires that we search for states that are not only feasible and stable, but can also resist the invasion by any species from the original pool that has otherwise gone extinct. Interestingly, the model without migration can lead to states that would be invadable by some species that have however gone extinct during the dynamics or that where not present at initial conditions. The inclusion of species migration in the GLV model modifies the multistability regime where EC is found, by transforming it into a regime where alternative stable states become less common, as some of these states are always invadable by otherwise absent species. As studied in (Mallmin *et al.*, 2024; Roy *et al.*, 2020) or (Arnoulx de Pirey and Bunin, 2024), this leads to a regime where the system approaches a community state but, instead of settling there, there is almost always another species from the pool that can invade this state and so on and so forth, leading to a kind of pinball dynamics between attractors (Aguadé-Gorgorió and Kefi, 2024; Roy *et al.*, 2020). This effectively leads to scenarios where alternative stable states with few surviving species are possible, but large communities with e.g. 10-15 species would become rare. While this might not be directly related to the specific experimental setups under study, the possibility of permanent abundance fluctuations has key implications in ecosystems such as plankton communities (Mallmin *et al.*, 2024; Roy *et al.*, 2020). The presence of EC in the GLV model with migrations has been discussed in Section II.A.5, while the reduction of stable states in favor of persistent fluctuations in this same exact model can be found in (Aguadé-Gorgorió and Kefi, 2024).

### F. Extended Discussion

### G. Emergent coexistence from chains of pairwise interactions

Species-rich microbial communities assembled in laboratory conditions often contain species that do not coexist in pairs due to strong competition (Chang *et al.*, 2023). This questions the bottom-up, reductionist approach of predicting species coexistence from pairwise observations alone (Friedman *et al.*, 2017), and highlights the possibility that additional non-pairwise mechanisms exist (Levine *et al.*, 2017).

We have shown that species-rich models of simple pairwise interactions are consistent with observations of EC. Starting from the simplest possible model with random interactions, we have uncovered a threshold in interaction heterogeneity, beyond which EC becomes pervasive and most stable communities harbor excluding pairs. The model predicts a limit on the fraction of excluding pairs that a community can sustain, that is consistent with observations from three different experimental datasets.

Coexistence between strong competitors is maintained by positive indirect effects, a well-known result for few-species models (Levine, 1999, 1976). Yet, as diversity increases, indirect effects can rapidly become so intrincate that direct interactions no longer predict EC (Zelnik *et al.*, 2024), imposing a fundamental limit to the reductionist approach and an explanation for observations of unexpected coexistence between strong competitors. By bringing together recent mathematical results (Aguadé-Gorgorió and Kefi, 2024; Zelnik *et al.*, 2024) and empirical data (Chang *et al.*, 2023; Friedman *et al.*, 2017; Venturelli *et al.*, 2018), our work reinforces the need to explicitly assess the weight of indirect effects –through collectivity $\phi$ and condition number $k$– when forecasting community behavior.

Finally, we have shown that communities with more than 3 species do not require intransitive competition, aligning again with empirical observations of microbial communities (Chang *et al.*, 2023; Higgins *et al.*, 2017; Koch *et al.*, 2023). In the limit case of nested or single-resource competition, communities can maintain EC even under a total absence of intransitive loops.

### H. Emergent coexistence is an emergent phenomenon

The term *emergence*, although often used loosely across complex systems research, has formal mathematical definitions (Artime and De Domenico, 2022). Specifically, *weak emergence* describes scenarios where large-scale patterns are unexpected, but can –in principle– be computationally derived from lower-scale mechanisms. *Strong emergence*, instead, describes situations where patterns are unrelated or autonomous from any lower-scale mechanism (Artime and De Domenico, 2022).

Our work shows that EC falls in the category of weak emergence: in the presence of strong indirect effects, community composition is driven by, but cannot be analytically predicted from, direct interactions between species. Numerical approximations of species composition and net effects are possible through integration of the dynamics or matrix inversion, albeit with limited precision.

Our work places the phenomenon of EC in a similar conceptual category as deterministic chaos. In chaos, simple dynamical rules produce systems with complex behavior, where small differences in initial conditions can grow exponentially, limiting the capacity for long-term dynamical predictions (May, 1976). Similarly, in EC, unexpected species coexistence can emerge from long chains of simple pairwise interactions. This places EC and indirect effects within broader themes in complex systems, where self-organization, criticality or nonlinear feedbacks can shape emergent behaviors (Boffetta *et al.*, 2002; Levins, 1974; Solé and Bascompte, 2012).

#### I. The limits of bottom-up predictability in species coexistence

Understanding our capacity to predict the future configuration of a system given its present state is a central problem in ecology and across complex systems (Boffetta *et al.*, 2002). In ecology, predictability is often studied in terms of forecasting temporal responses to perturbations, such as species invasions, extinctions, or environmental change (Bender *et al.*, 1984; Daugaard *et al.*, 2022).

Ecological complexity hinders our forecasting capacity in two main ways. First, parametrizing a mathematical model to describe a community of moderate size relies on inferring tens or hundreds of parameters, a tremendously complex task (Arya *et al.*, 2023; Camacho-Mateu *et al.*, 2024; Picot *et al.*, 2023; Rosenbaum and Fronhofer, 2023). Second, even if we could infer the strengths of species interactions, recent work has uncovered scenarios where complex dynamical behavior can limit our capacity to predict temporal responses to perturbations (Gilpin, 2024; Kawatsu, 2024; Mallmin *et al.*, 2024; Zelnik *et al.*, 2024).

Microbial community assembly provides an alternative approach to test ecological predictability, by studying if long-term coexistence is predictable from few-species observations (Chang *et al.*, 2023; Friedman *et al.*, 2017; Pennekamp *et al.*, 2018). Yet, our work establishes explicit and computable limits in the capacity to predict community composition from species interactions, adding a novel perspective on the overall limitations of the temporal and, now, compositional forecasting of ecosystems. This has explicit and practical implications for the rational design and bottom-up assembly of microbial consortia for applications from biomedicine (Eng and Borenstein, 2019) to environmental restoration (Solé *et al.*, 2024).

While weaker or non-random interactions may relax these limitations in small communities, our findings regarding EC suggest profound implications in our capacity to predict the detailed makeup of larger natural communities of hundreds of species. Even more, we hypothesize that incorporating additional mechanisms such as higher-order interactions would likely decrease predictability even further. Instead, a coarse-grained framework is needed –one that infers global community properties without requiring precise species-level information. Towards that goal, theory could benefit from empirical evidence suggesting that, despite high species diversity, microbial communities often organize into a few functional groups (Arya *et al.*, 2023; Buche *et al.*, 2025; Estrela *et al.*, 2022; Goldford *et al.*, 2018).

#### J. Incorporating non-random and non-pairwise interactions

Assuming that interaction strengths are randomly distributed might be a valuable starting point to define null expectations in the absence of additional details of a system (Barbier *et al.*, 2021). Nevertheless, our predictions of EC are robust to different interactions structures, such as sparse or cross-feeding matrices, and are consistent with empirical observations. This places our work within recent successes linking species-rich models with microbial community data (Camacho-Mateu *et al.*, 2024; Grilli, 2020; Hu *et al.*, 2022; Pasqualini *et al.*, 2024).

A natural next step is to study how empirical interaction matrices modulate the weight of indirect effects and hence determine community predictability. For example, recent work has proposed that specific food-web architectures could modulate collectivity (Zelnik *et al.*, 2024), while sparse interactions in turn could limit the length of indirect effects by limiting longer chains. Again, random interactions provide a null expectation for collectivity, to which we could compare $\phi$ and $k$ values measured from empirical data.

Beyond the structure of interaction networks, additional mechanisms such as consumer-resource dynamics (Estrela *et al.*, 2022), higher-order effects (Billick and Case, 1994), nonlinear facilitation (Aguadé-Gorgorió *et al.*, 2024b) or time-dependent interactions (Yin and Rudolf, 2024) can modulate ecological dynamics. Our work does not claim that any of these mechanisms are not present in ecological communities. Instead, we highlight that, even in their absence, emergent behavior is already possible and limits our capacity to forecast community composition.

#### K. Conclusion

Using empirical observations of EC as a case study, our work highlights the critical role of indirect effects in shaping emergent community behavior. Even when interactions are strictly pairwise, the complexity of indirect effects rapidly escalates in communities with just a few strongly interacting species. This imposes a fundamental limit on our ability to predict community composition from pairwise observations: much like deterministic chaos emerging from simple dynamical equations, unexpected coexistence can emerge from long chains of species interactions. As a result, measuring all these interactions in natural communities with hundreds of species is not only an immense challenge: in some cases, it may also be insufficient. Our findings emphasize the need for coarse-grained models that capture global community properties without relying on precise species-level measurements.

**FIGURES**

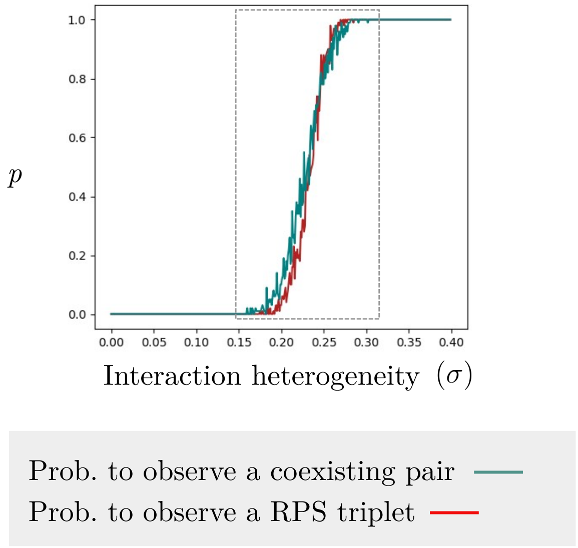


FIG. 1 **Probability of observing certain motifs in a random matrix.** We simulate 100 matrices with 80x80 elements sampled from $\mathcal{N}(\mu, \sigma)$, with $\mu = -1.5$ and $\sigma$ increasing (x-axis). We measure across these matrices the fraction of times in which a matrix has at least one coexisting pair ($A_{ij} > -1$, $A_{ji} > -1$, in teal) and the fraction of times a matrix has at least one rock-paper-scissors triplet (in red). Both motifs provide a proxy for the minimal standard deviation $\sigma$ by which a large matrix will transition from very rare coexistence ($p = 0$) to almost ensured coexistence ($p = 1$). This transition happens quite sharply for both motifs, between $\sigma = 0.2$ and $\sigma = 0.26$, and is consistent with the analytical estimate shown in figure 2B of the main text, bottom panel.

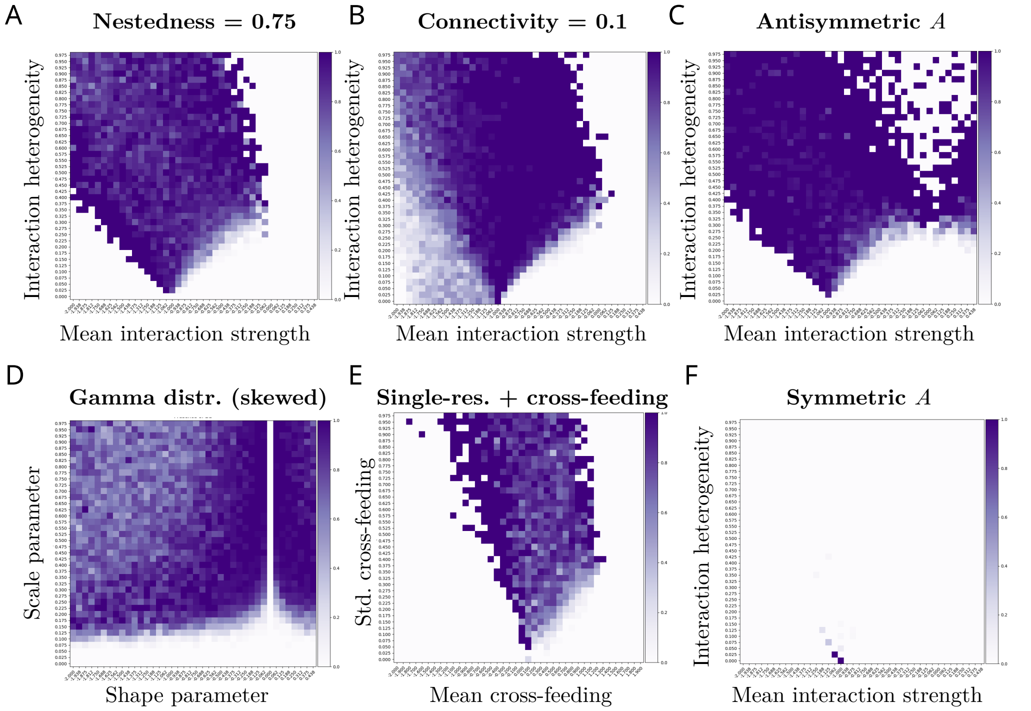


FIG. 2 **Emergent coexistence under different matrix randomizations.** In purple the fraction of stable states that contain at least one excluding pair, using the equivalent numerical procedure of figure 2A of the main text, but under different types of interaction strength randomizations that are not the fully random, uncorrelated $A_{ij} \in \mathcal{N}(\mu, \sigma)$ elements of the main text. We find that EC is a common event (dark purple, most or all states contain at least one excluding pair) across different matrix randomizations even if the shape of the phase spaces can change depending on the model under study. The only exception is in the limit-case scenario of fully symmetric interactions ($A_{ij} = A_{ji}$) where EC is very rare (panel F). In Section I.L we describe in detail the way each of the randomizations is build and the interpretation of the x- and y-axes and simulation outcomes.

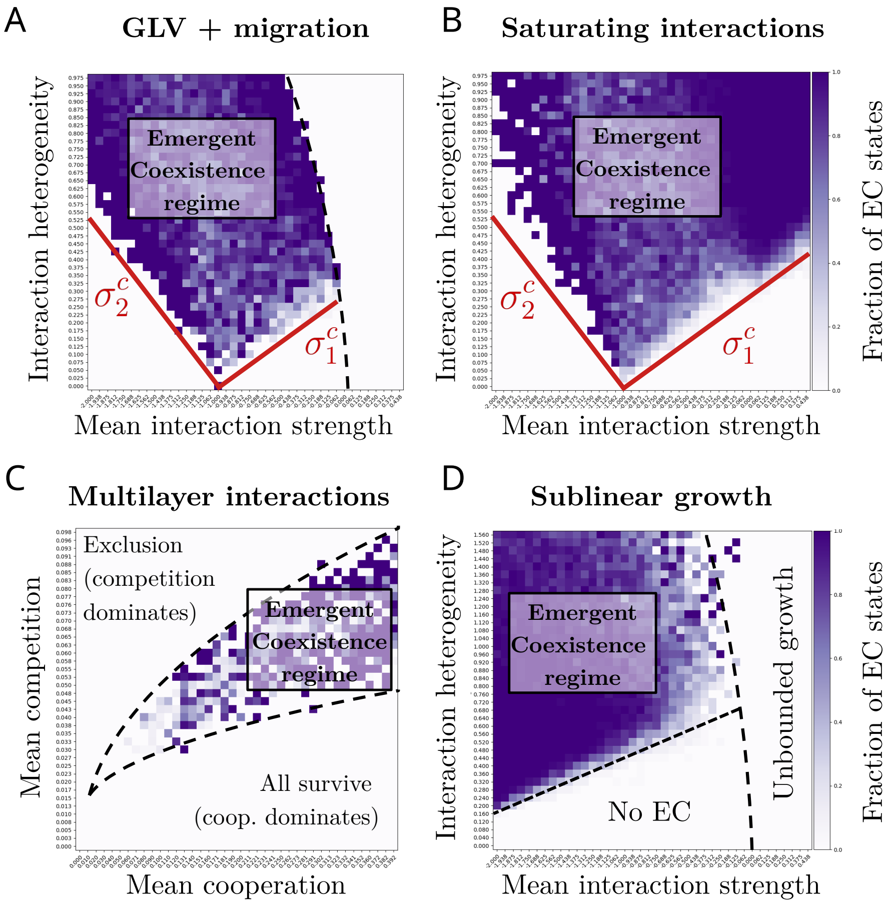


FIG. 3 **Emergent coexistence in different dynamical models.** See Section II.A.5 for the extended figure caption.

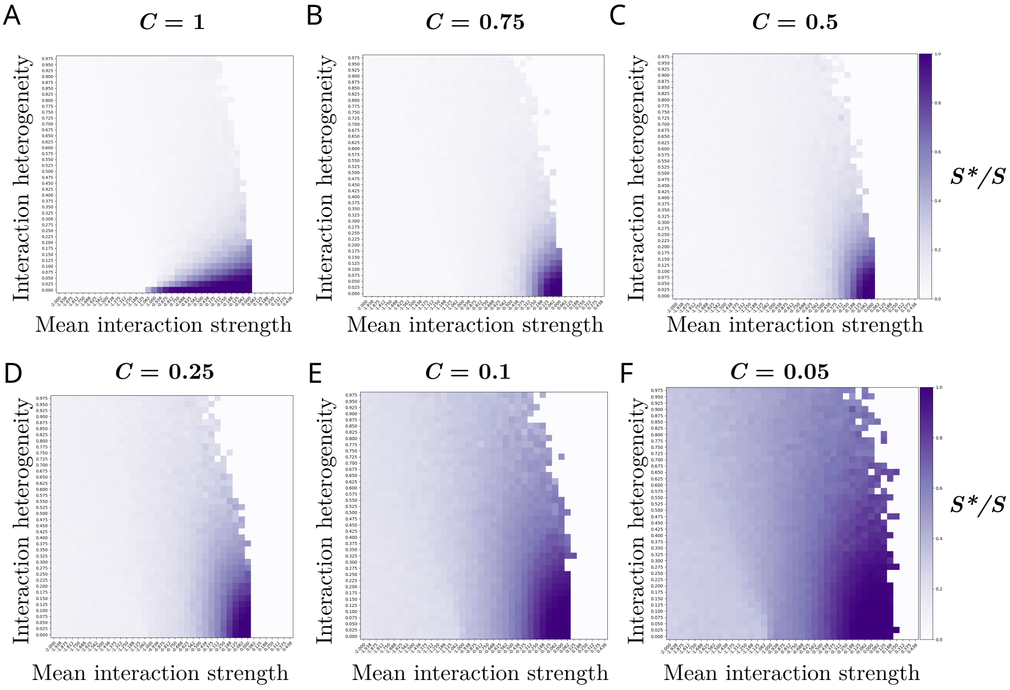


FIG. 4 **Fraction of surviving species under decreasing connectivity.** Phase space of the GLV model under decreasing connectivity. It is important to note that the $\mu$ and $\sigma$ values of the $x$ and $y$ axes do not refer to the mean and variance of the final matrix after the sparsity filter is introduced, but rather to the values of the original $A$ matrix before it is filled with zeros. Although the other method can also be interesting, this provides a useful way of visualizing how connectivity changes the space of possible outcomes. The interesting aspect of the result is related to the fact that reduced connectivity reduces $\mu$ linearly, while it first increases $\sigma$ and later decreases it. This implies that reductions of connectivity as low as $C = 0.5$ do not seem to result in relevant changes in species abundances, whereas much lower connectivity values result in a very homogeneous distribution of effective interactions (most $A_{ij}$ become null), leading to an obvious increase in the possible diversity of the system as many species do not interact directly.

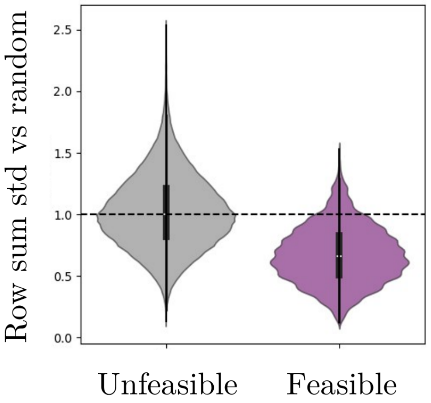


FIG. 5 **Row sums of coexisting species interaction matrices are homogeneous.** Here we plot the violin plots (using the seaborn.violinplot function in python) of the standard deviation of row sums of a $10^6$ matrices of interactions $A^*$ sampled from $A$ with $\mu$ and $\sigma$ sampled randomly within the domain of figure 2A in the main text. We divide the standard deviation by the average standard deviation of 100 randomizations of each matrix generated by shuffling all off-diagonal elements. A value of 1 indicates that the row sums of the interaction matrix are as heterogeneous as the row sums of the randomized versions, whereas a value smaller than 1 indicates that row sums are more homogeneous than the random expectation. Unfeasible communities, where the abundances of some species in a given subset are not positive and hence there is no coexistence, have row sum values similar to those of a random matrix. Feasible communities, instead, have more homogeneous row sums than the random expectation, meaning that there is a tendency or filter for which species can coexist if they all perceive competition $\sum_j A^*_{ij}$ in a similar way.

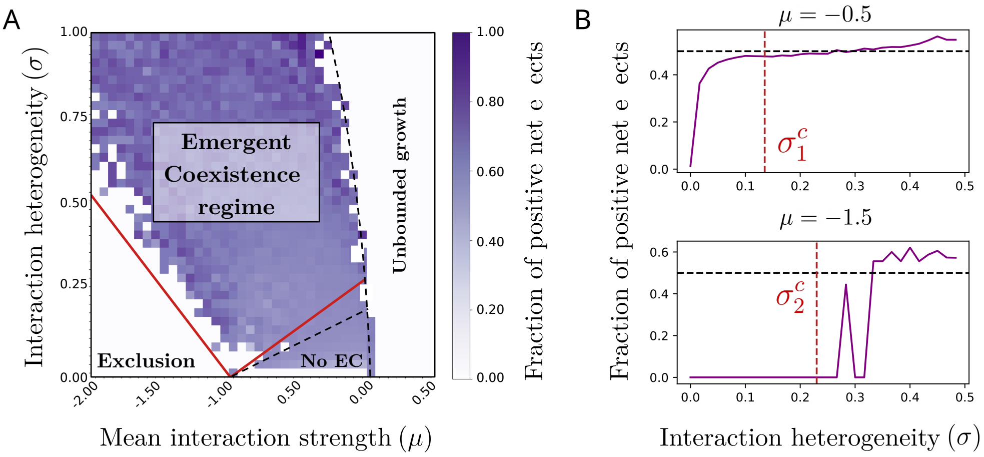


FIG. 6 **Fraction of positive net effects ($(I - A^*)^{-1}_{ij} > 0$)).** Equivalent plot to figure 2A of the main text but, instead of measuring the fraction of states that harbor emergent coexistence, we measure the fraction of positive elements present in $(I - A^*)^{-1}$ given a stable state with interactions encoded in $A^*$ found after simulating the GLV model with pool interactions $A \sim \mathcal{N}(\mu, \sigma)$. In (A) we plot the phase space, to observe that much before EC appears, the fraction of positive net effects in a community rapidly approaches 0.5, meaning that half of the effects between species are in fact facilitative. Equivalent to figure 2A in the main text, in (B) we plot two vertical slices of (A) for $\mu = -0.5$ and $\mu = -1.5$ and increasing $\sigma$.

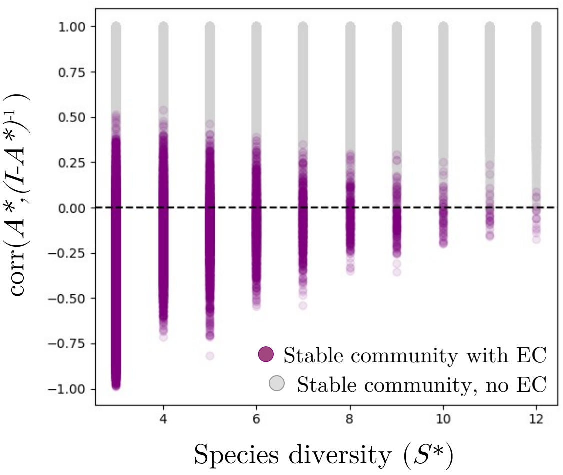


FIG. 7 **Correlation between direct and net effects.** Consistent with discussions in the main text and Section II.B.2, here we measure the Pearson correlation between the matrix of direct interactions and the matrix of net effects. As expected, we find that in communities without EC (gray) characterized by weak interactions the two matrices correlate quite well, meaning that direct interactions provide certain information on net effects, mainly because long indirect effects are negligible. Instead, for communities with EC (purple) that are characterized by the presence of strong interactions, the two matrices do not necessarily correlate: for small communities, they can be anti-correlated, meaning that direct competitors are in fact the species that are providing a rescue effect. For more diverse communities, the two matrices become uncorrelated, leading to the predictability limit proposed in the main text: direct interactions hold little information about net effects and species abundances, mostly because they are negligible when compared to much longer and heavier chains of indirect effects.

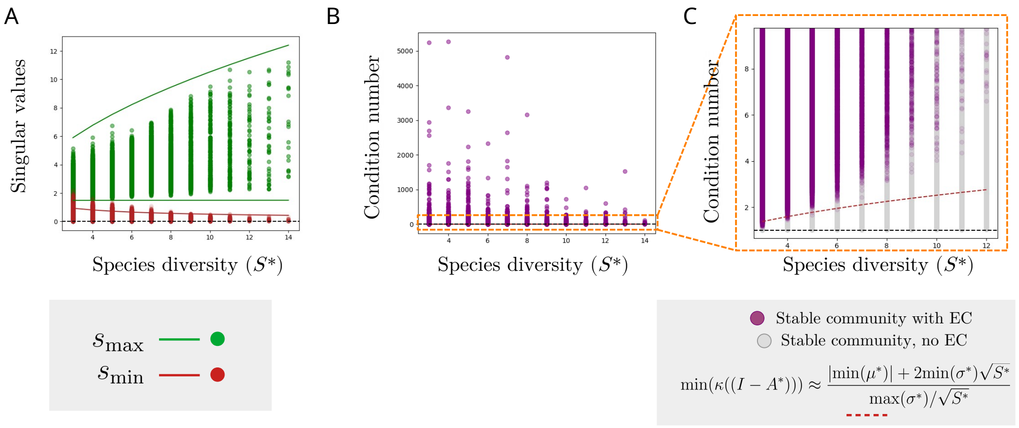


FIG. 8 **Numerical and analytical predictions for the condition number $\kappa$.** In (A) we plot the largest and smallest singular values found for a large number of communities with EC found after simulating $10^6$ systems within the range of figure 2A in the main text. The circles are the values for each state computed with `numpy.linalg.svd`, whereas the lines refer to the maximum and minimum predictions for both $s_{\max}$ and $s_{\min}$ from Random Matrix Theory (Section II.B.4). In (B) we plot the condition number for those states that results from dividing $s_{\max}/s_{\min}$, and in (C) we zoom in and also add the values for communities that do not harbor EC, which allows us to both visualize the analytical prediction proposed in Section II.B.4 together with the observation that $\kappa$ will only remain close to 1 for communities with very weak and homogeneous interaction strengths. As interactions become relatively strong and heterogeneous –as for communities with EC, most communities have a condition number ($\kappa$) much larger than 1. The best expectation is that the condition number increases as $\kappa \sim (S^*)$, but most communities fall much above this analytical estimate. The consequence is that small errors in the measurement of $A^*$ can translate to large errors in the prediction of species abundances via matrix inversion.

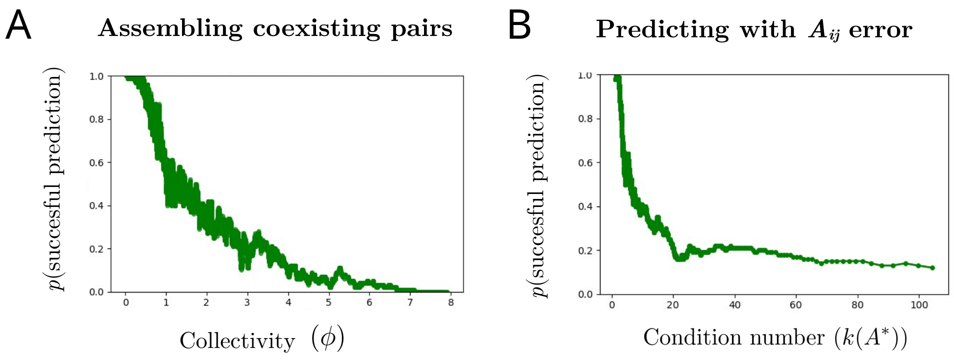


FIG. 9 **Testing the role of $\phi$ and $k$ in possible experimental tests of community assembly.** In (A) we study if assembling pairs of coexisting species will result in a coexisting community as a function of $\phi$. As discussed throughout sections I.I and II.B.5, we see how communities with very small collectivity (few species, weak indirect effects) result in a success, meaning that the final community coexists. Yet, as $\phi$ increases, the likelihood that the reductionist approach succeeds decreases rapidly, so that assembling species that coexist by pairs will not result in a successful, coexisting community. Instead, EC indicates that indirect effects dominate over the community, and assembly tests based on pairwise interactions alone will not succeed. In (B) we test if the predicted coexistence based on imprecise measurements of $A^*_{ij} + \epsilon$, with $\epsilon \sim \mathcal{N}(\mu = 0, \sigma = 0.1)$, successfully predicts the coexistence of the real community with interactions $A^*_{ij}$. We find that, as condition number increases, small errors in our measurements of species interactions rapidly propagate, and for $k \approx 20$, most of the predicted coexisting communities will not coexist in reality.

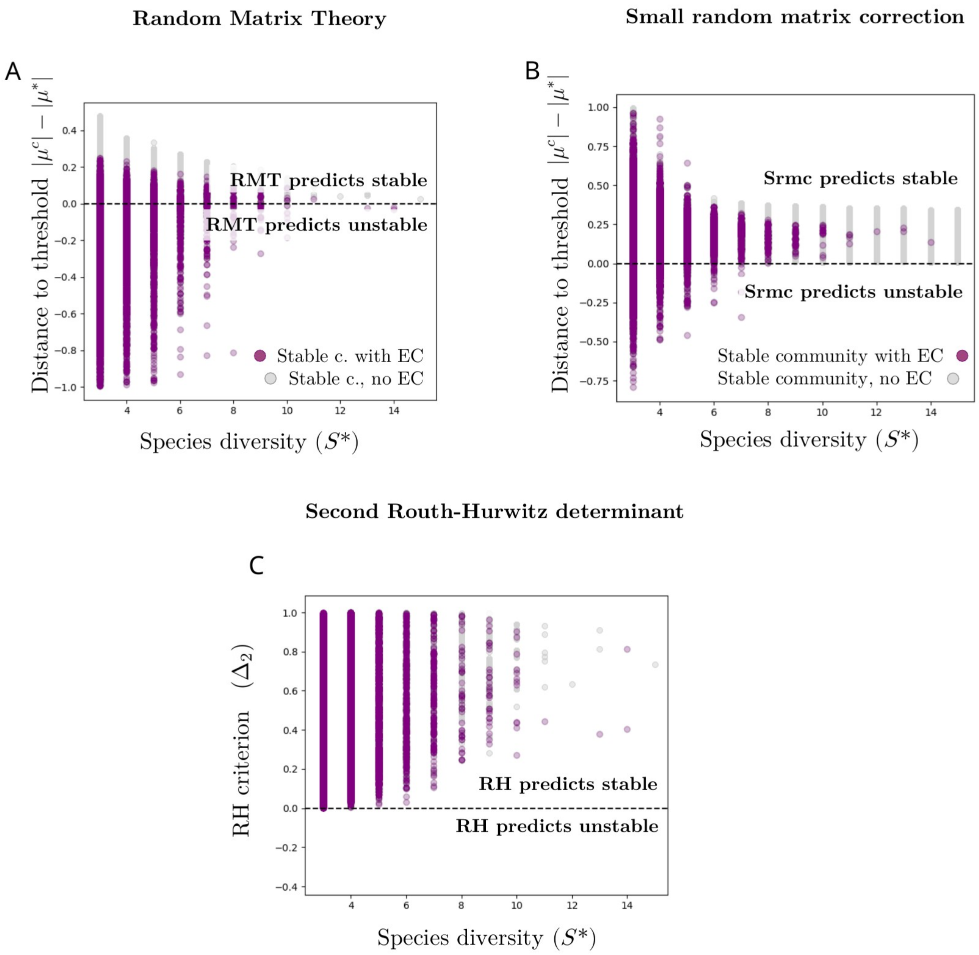


FIG. 10 **Stability predictions for EC states.** Different stability predictions based on the properties of $A^*$ and $J^*$ for communities with (purple) and without (gray) EC, where communities with EC are in fact communities with strong competition. In (A) we see that many communities have stronger competition than what the Random Matrix Theory threshold predicts (Section I.J). Once we correct the threshold using recent approximations for communities of moderate size in (B), we see that more communities fulfill the stability prediction and have absolute interaction strength weaker than the limit prediction $|\mu^c|$. However, it is clear that for the smallest communities these approximations for large and moderate sized communities do not apply, and more subtle patterns allow the states to maintain stability even under strong competition. These patterns are better captured by the Routh-Hurwitz criteria and in particular the condition on the second determinant (Section I.K). We can see that all communities, with and without EC, fulfill the condition for a positive second RH determinant, mostly because it is not based on aggregate statistics of the community but rather the exact values and locations of interaction strengths.

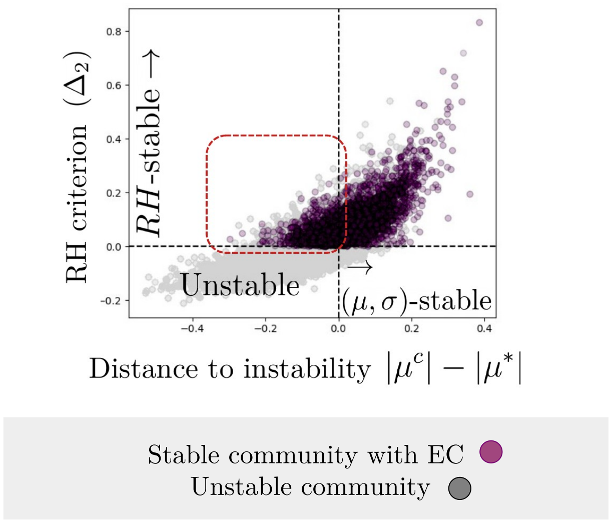


FIG. 11 **Random matrix theory and Routh-Hurwitz predictions on stability.** Bringing together the results of the figure in the previous page, here we plot the distance to May's threshold (x-axis) and the sign of the second Routh-Hurwitz determinant (y-axis) for stable communities with EC (purple) and unstable communities (gray) sampled from the GLV model within the parameter domain of figure 2A in the main text. While many communities have stronger competition than what the Random Matrix Theory approximation predicts (dashed red square, Random Matrix Theory does not operate for such small communities), they all fulfill the RH criteria, providing a microscopic, interaction-structure explanation for linear stability under strong competition.

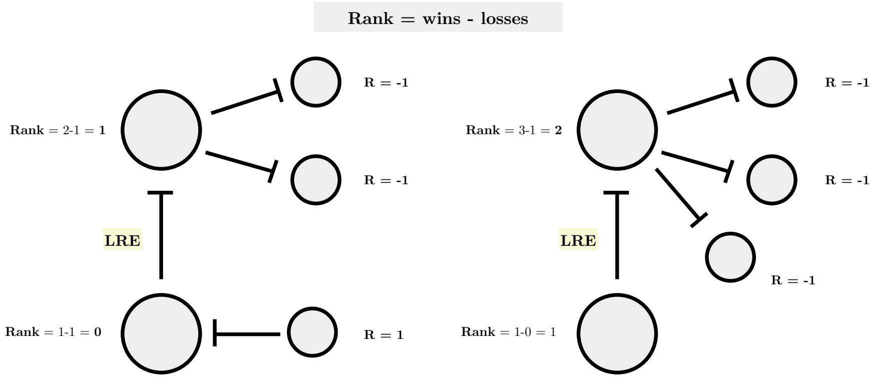


FIG. 12 **Low rank exclusions require at least four exclusionary interactions.** Circles represent species and arrows represent exclusionary interactions. We do not pain the chains of non-exclusionary interactions that we have shown could lead to species coexistence. In any case, we show that for a LRE to happen, we require a minimum of four exclusionary interactions. Because the higher-ranked species will have at least one loss (that of the LRE), it requires additional wins with other species in the community to qualify as a LRE. This shows that LRE are in fact more complex and specific motifs than what the initial intuition proposes, and they might not be a good metric to assess intransitivity in systems with few species or few exclusionary elements.

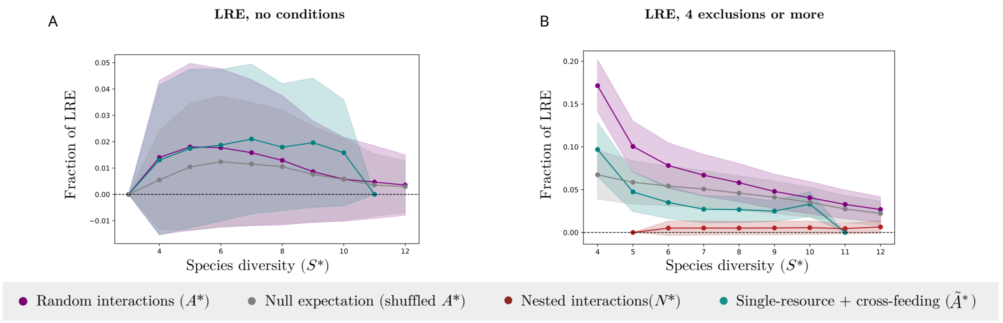


FIG. 13 **Low rank exclusions without and with the 4-exclusions requirement.** LRE requires at least 4 exclusionary interactions (see figure above). If we do not consider this requirement, the apparent fraction of exclusions that is LRE is extremely low (A), mostly because there are a lot of communities with 3 or fewer exclusions that will never harbor a LRE. Instead, in (B) we plot the figure of the main text that corrects for this requirement, and studies the presence of LRE in systems with 4 or more interactions only. Both for the unconditioned case (A) and for the conditioned case (B), we can see that large communities very rarely harbor LRE's, consistent with empirical observations.

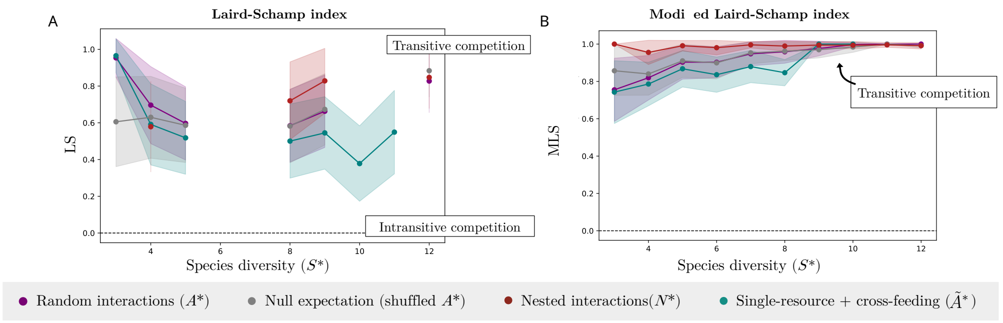


FIG. 14 **Laird-Schamp index with and without $s_{\min}$.** The original intransitivity index used in Laird and Schamp (Laird and Schamp, 2006) fails in our model when the divisor $s_{\max} - s_{\min} = 0$, which happens if there is only one exclusionary element in a system and so there is no such maximum or minimum rank heterogeneity. We can correct this by defining a less restrictive index $s_{obs}/s_{\max}$ (Section II.D.3). If this index equals 1, it means that species ranks are distributed in the most hierarchical (heterogeneous) way, indicative of a transitive competition matrix. The results are not surprising and qualitatively equivalent to the two intransitivity indexes presented in the main text, which indicate that clear intransitive motifs disappear with species diversity, where in fact few exclusions are present and many indirect effects explain the survival of these excluded species in the absence of intransitive motifs.

### BIBLIOGRAPHY

### REFERENCES


Aguadé-Gorgorió, G., A. R. Anderson, and R. Solé (2024a), Iscience **27** (9).

Aguadé-Gorgorió, G., J.-f. Arnoldi, M. Barbier, and S. Kéfi (2024b), Ecology Letters **27** (4), e14413.

Aguadé-Gorgorió, G., and S. Kefi (2024), Journal of Physics: Complexity.

Aguadé-Gorgorió, G., I. Lajaaiti, J.-f. Arnoldi, and S. Kéfi (2025), Oikos **2025** (1), e10980.

Allesina, S., and J. M. Levine (2011), Proceedings of the National Academy of Sciences **108** (14), 5638.

Allesina, S., and S. Tang (2012), Nature **483** (7388), 205.

Altieri, A., and G. Biroli (2022), SciPost Physics **12** (1), 013.

Altieri, A., F. Roy, C. Cammarota, and G. Biroli (2021), Physical Review Letters **126** (25), 258301.

Arnoldi, J.-F., M. Barbier, R. Kelly, G. Barabás, and A. L. Jackson (2022), Methods in Ecology and Evolution **13** (1), 167.

Artime, O., and M. De Domenico (2022), Philosophical Transactions of the Royal Society A **380** (2227), 20200410.

Arya, S., A. B. George, and J. O'Dwyer (2025), Current Opinion in Microbiology **83**, 102580.

Arya, S., A. B. George, and J. P. O'Dwyer (2023), Proceedings of the National Academy of Sciences **120** (48), e2307313120.

Barabás, G., M. J. Michalska-Smith, and S. Allesina (2017), Nature ecology & evolution **1** (12), 1870.

Barbier, M., J.-F. Arnoldi, G. Bunin, and M. Loreau (2018), Proceedings of the National Academy of Sciences **115** (9), 2156.

Barbier, M., C. De Mazancourt, M. Loreau, and G. Bunin (2021), Physical Review X **11** (1), 011009.

Bascompte, J. (2009), Frontiers in Ecology and the Environment **7** (8), 429.

Bascompte, J. (2010), Science **329** (5993), 765.

Bender, E. A., T. J. Case, and M. E. Gilpin (1984), Ecology **65** (1), 1.

van den Berg, N. I., D. Machado, S. Santos, I. Rocha, J. Chacón, W. Harcombe, S. Mitri, and K. R. Patil (2022), Nature ecology & evolution **6** (7), 855.

Billick, I., and T. J. Case (1994), Ecology **75** (6), 1529.

Bimler, M., M. Mayfield, and A. James (2025), .

Biroli, G., G. Bunin, and C. Cammarota (2018), New Journal of Physics **20** (8), 083051.

Bodson, M. (2020), IEEE Control Systems Magazine **40** (1), 45.

Boffetta, G., M. Cencini, M. Falcioni, and A. Vulpiani (2002), Physics reports **356** (6), 367.

Buche, L., L. G. Shoemaker, L. M. Hallett, I. Bartomeus, P. Vesk, C. Weiss-Lehman, M. Mayfield, and O. Godoy (2025), Ecology Letters **28** (1), e70059.

Bunin, G. (2017), Physical Review E **95** (4), 042414.

Calleja-Solanas, V., N. Khalil, J. Gómez-Gardeñes, E. Hernández-García, and S. Meloni (2022), Physical Review E **106** (6), 064307.

Camacho-Mateu, J., A. Lampo, M. Sireci, M. A. Muñoz, and J. A. Cuesta (2024), Proceedings of the National Academy of Sciences **121** (5), e2309575121.

Castillo-Alvino, H., and M. Marvá (2020), Journal of Biological Dynamics **14** (1), 222.

Castledine, M., J. Pennycook, A. Newbury, L. Lear, Z. Erdos, R. Lewis, S. Kay, D. Sanders, D. Sünderhauf, A. Buckling, *et al.* (2024), Microbiology **170** (9), 001489.

Cenci, S., and S. Saavedra (2018), Physical Review E **97** (1), 012401.

Chang, C.-Y., D. Bajić, J. C. Vila, S. Estrela, and A. Sanchez (2023), Science **381** (6655), 343.

Chen, Y.-C. (2017), Biostatistics & Epidemiology **1** (1), 161.

Chesson, P. (1990), Theoretical Population Biology **37** (1), 26.

Chesson, P. (2000), Annual review of Ecology and Systematics **31** (1), 343.

Clark, R. N. (1992), IEEE Control Systems Magazine **12** (3), 119.

Courchamp, F., L. Berec, and J. Gascoigne (2008), *Allee effects in ecology and conservation* (OUP Oxford).

Czárán, T., and E. Szathmáry (2000), The geometry of ecological interactions **116**, 134.

Dal Bello, M., H. Lee, A. Goyal, and J. Gore (2021), Nature Ecology and Evolution **5** (10), 1424.

Dambacher, J. M., H.-K. Luh, H. W. Li, and P. A. Rossignol (2003), The American Naturalist **161** (6), 876.

Daugaard, U., S. B. Munch, D. Inauen, F. Pennekamp, and O. L. Petchey (2022), Ecology Letters **25** (9), 1974.

Davis, P. J. (1959), The American Mathematical Monthly **66** (10), 849.

Demmel, J. W. (1987), Numerische Mathematik **51**, 251.

Domínguez-García, V., V. Dakos, and S. Kéfi (2019), Proceedings of the National Academy of Sciences **116** (51), 25714.

Donohue, I., H. Hillebrand, J. M. Montoya, O. L. Petchey, S. L. Pimm, M. S. Fowler, K. Healy, A. L. Jackson, M. Lurgi, D. McClean, *et al.* (2016), Ecology letters **19** (9), 1172.

Dormand, J. R., and P. J. Prince (1980), Journal of computational and applied mathematics **6** (1), 19.

Dormann, C. F. (2007), Plant Ecology **191**, 171.

Dormann, C. F., and S. H. Roxburgh (2005), Proceedings of the Royal Society B: Biological Sciences **272** (1569), 1279.

Dunne, J. A. (2006), Ecological networks: linking structure to dynamics in food webs **1**, 27.

Edelman, A. (1988), SIAM journal on matrix analysis and applications **9** (4), 543.

El Ghaoui, L. (2002), Linear algebra and its applications **343**, 171.

Eng, A., and E. Borenstein (2019), Current opinion in biotechnology **58**, 117.

Engel, E. C., and J. F. Weltzin (2008), Plant Ecology **195**, 77.

Estrela, S., J. C. Vila, N. Lu, D. Bajić, M. Rebolleda-Gómez, C.-Y. Chang, J. E. Goldford, A. Sanchez-Gorostiaga, and Á. Sánchez (2022), Cell Systems **13** (1), 29.

Feng, Y., S. Soliveres, E. Allan, B. Rosenbaum, C. Wagg, A. Tabi, E. De Luca, N. Eisenhauer, B. Schmid, A. Weigelt, *et al.* (2020), Methods in Ecology and Evolution **11** (1), 117.

Fortuna, M. A., D. B. Stouffer, J. M. Olesen, P. Jordano, D. Mouillot, B. R. Krasnov, R. Poulin, and J. Bascompte (2010), Journal of animal ecology , 811.

Friedman, J., L. M. Higgins, and J. Gore (2017), Nature ecology & evolution **1** (5), 0109.

Galla, T. (2018), Europhysics Letters **123** (4), 48004.

Gallien, L., N. E. Zimmermann, J. M. Levine, and P. B. Adler (2017), Ecology Letters **20** (7), 791.

Ghoul, M., and S. Mitri (2016), Trends in microbiology **24** (10), 833.

Gilpin, M. E. (1975), The American Naturalist **109** (965), 51.

Gilpin, W. (2024), arXiv preprint arXiv:2403.19186.

Giral Martínez, J., M. Barbier, and S. De Monte (2024), bioRxiv , 2024.

Godoy, O., D. B. Stouffer, N. J. Kraft, and J. M. Levine (2017), "Intransitivity is infrequent and fails to promote annual plant coexistence without pairwise niche differences,".

Goldford, J. E., N. Lu, D. Bajić, S. Estrela, M. Tikhonov, A. Sanchez-Gorostiaga, D. Segrè, P. Mehta, and A. Sanchez (2018), Science **361** (6401), 469.

Graham, M. H. (2003), Ecology **84** (11), 2809.

Grilli, J. (2020), Nature communications **11** (1), 4743.

Grilli, J., M. Adorisio, S. Suweis, G. Barabás, J. R. Banavar, S. Allesina, and A. Maritan (2017a), Nature communications **8** (1), 14389.

Grilli, J., G. Barabás, M. J. Michalska-Smith, and S. Allesina (2017b), Nature **548** (7666), 210.

Grilli, J., T. Rogers, and S. Allesina (2016), Nature communications **7** (1), 12031.

Guimaraes Jr, P. R. (2020), Annual Review of Ecology, Evolution, and Systematics **51** (1), 433.

Hardin, G. (1960), science **131** (3409), 1292.
Hatton, I. A., O. Mazzarisi, A. Altieri, and M. Smerlak (2024), Science **383** (6688), eadg8488.
Higgins, L. M., J. Friedman, H. Shen, and J. Gore (2017), BioRxiv , 175737.
Holling, C. S. (1959), The canadian entomologist **91** (5), 293.
Horn, R. A., and C. R. Johnson (1994), *Topics in matrix analysis* (Cambridge university press).
Hu, J., D. R. Amor, M. Barbier, G. Bunin, and J. Gore (2022), Science **378** (6615), 85.
Hurwitz, A. (1895), Mathematische Annalen **46** (2), 273.
Hutchinson, G. E. (1953), Proceedings of the Academy of Natural Sciences of Philadelphia **105**, 1.
Ives, A. R., and S. R. Carpenter (2007), science **317** (5834), 58.
Jacquet, C., C. Moritz, L. Morissette, P. Legagneux, F. Massol, P. Archambault, and D. Gravel (2016), Nature communications **7** (1), 12573.
Johnson, R. A., D. W. Wichern, *et al.* (2002), .
Kawatsu, K. (2024), Proceedings of the National Academy of Sciences **121** (27), e2322939121.
Kéfi, S., V. Domínguez-García, I. Donohue, C. Fontaine, E. Thébault, and V. Dakos (2019), Ecology letters **22** (9), 1349.
Kéfi, S., C. J. Lortie, and L. A. Cavieres (2024), "The importance of facilitative interactions in mediating climate change impact on biodiversity,".
Kerr, B., M. A. Riley, M. W. Feldman, and B. J. Bohannan (2002), Nature **418** (6894), 171.
Kessler, D. A., and N. M. Shnerb (2015), Physical Review E **91** (4), 042705.
Kessler, D. A., and N. M. Shnerb (2025), Physical Review E **111** (3), 034408.
von Kiedrowski, G. (1993), Bioorganic chemistry frontiers , 113.
Koch, F., A.-M. Neutel, D. K. Barnes, and K. T. Allhoff (2024), bioRxiv , 2024.
Koch, F., A.-M. Neutel, D. K. Barnes, K. Tielborger, C. Zarfl, and K. T. Allhoff (2023), Communications Biology **6** (1), 690.
Kvrivan, V., and J. Eisner (2006), Theoretical Population Biology **70** (4), 421.
Laigle, I., I. Aubin, C. Digel, U. Brose, I. Boulangeat, and D. Gravel (2018), Oikos **127** (2), 316.
Laird, R. A., and B. S. Schamp (2006), The American Naturalist **168** (2), 182.
Lajaaiti, I., S. Kéfi, and J.-F. Arnoldi (2024), Proceedings of the Royal Society B **291** (2032), 20240930.
Landi, P., H. O. Minoarivelo, Å. Brännström, C. Hui, and U. Dieckmann (2018), Population ecology **60** (4), 319.
Lee, H., B. Bloxham, and J. Gore (2023), Proceedings of the National Academy of Sciences **120** (35), e2212113120.
Lele, K., B. E. Wolfe, and L. H. Uricchio (2024), bioRxiv , 2024.
Levine, J. M. (1999), Ecology **80** (5), 1762.
Levine, J. M., J. Bascompte, P. B. Adler, and S. Allesina (2017), Nature **546** (7656), 56.
Levine, S. H. (1976), The American Naturalist **110** (976), 903.
Levins, R. (1974), Annals of the New York Academy of Sciences **231** (1), 123.
Liautaud, K., E. H. van Nes, M. Barbier, M. Scheffer, and M. Loreau (2019), Ecology letters **22** (8), 1243.
Lubiana Botelho, L., C. Jeynes-Smith, S. A. Vollert, and M. Bode (2025), Ecology Letters **28** (1), e70034.
Mallmin, E., A. Traulsen, and S. De Monte (2024), Proceedings of the National Academy of Sciences **121** (11), e2312822121.
Marcus, S., A. M. Turner, and G. Bunin (2022), PLoS computational biology **18** (7), e1010274.
Marcus, S., A. M. Turner, and G. Bunin (2024), arXiv preprint arXiv:2405.11360.
Martínez, J. G., S. De Monte, and M. Barbier (2024), arXiv preprint arXiv:2411.14969.
May, R. M. (1972), Nature **238** (5364), 413.
May, R. M. (1976), Nature **261** (5560), 459.
May, R. M. (2019), *Stability and complexity in model ecosystems* (Princeton university press).
May, R. M., and W. J. Leonard (1975), SIAM journal on applied mathematics **29** (2), 243.

Mazzarisi, O., and M. Smerlak (2024), Physical Review E **110** (5), 054403.

McCann, K. S. (2000), Nature **405** (6783), 228.

Mehta, P., and R. Marsland III (2021), arXiv preprint arXiv:2110.04965.

Neutel, A.-M., J. A. Heesterbeek, and P. C. De Ruiter (2002), Science **296** (5570), 1120.

Neutel, A.-M., J. A. Heesterbeek, J. Van de Koppel, G. Hoenderboom, A. Vos, C. Kaldeway, F. Berendse, and P. C. De Ruiter (2007), Nature **449** (7162), 599.

Neutel, A.-M., and M. A. Thorne (2014), Ecology letters **17** (6), 651.

Newman, M. (2018), *Networks* (Oxford university press).

Ortiz, A., N. M. Vega, C. Ratzke, and J. Gore (2021), The ISME Journal **15** (7), 2131.

Pasqualini, J., A. Maritan, A. Rinaldo, S. Facchin, E. Savarino, A. Altieri, and S. Suweis (2024), arXiv preprint arXiv:2406.07465.

Payrató-Borras, C., L. Hernández, and Y. Moreno (2019), Physical Review X **9** (3), 031024.

Pearl Mizrahi, S., H. Lee, A. Goyal, E. Owen, and J. Gore (2025), bioRxiv , 2025.

Pennekamp, F., M. Pontarp, A. Tabi, F. Altermatt, R. Alther, Y. Choffat, E. A. Fronhofer, P. Ganesanandamoorthy, A. Garnier, J. I. Griffiths, *et al.* (2018), Nature **563** (7729), 109.

Pfeiffer, T., and S. Bonhoeffer (2004), The American Naturalist **163** (6), E126.

Picot, A., S. Shibasaki, O. J. Meacock, and S. Mitri (2023), Current Opinion in Microbiology **75**, 102354.

Pilosof, S., M. A. Porter, M. Pascual, and S. Kéfi (2017), Nature Ecology & Evolution **1** (4), 0101.

Piñero, J., and R. Solé (2018), Entropy **20** (2), 98.

Arnoulx de Pirey, T., and G. Bunin (2024), Physical Review X **14** (1), 011037.

Poley, L., T. Galla, and J. W. Baron (2025), Physical Review E **111** (1), 014318.

Qian, J. J., and E. Akçay (2020), Nature ecology & evolution **4** (3), 356.

Rao, C., K. Z. Coyte, W. Bainter, R. S. Geha, C. R. Martin, and S. Rakoff-Nahoum (2021), Nature **591** (7851), 633.

Rodriguez-Brenes, I. A., N. L. Komarova, and D. Wodarz (2013), Trends in ecology & evolution **28** (10), 597.

Rohr, R. P., S. Saavedra, and J. Bascompte (2014), science **345** (6195), 1253497.

Rosenbaum, B., and E. A. Fronhofer (2023), Ecosphere **14** (4), e4503.

Routh, E. J. (1877), *A treatise on the stability of a given state of motion: particularly steady motion. Being the essay to which the adams prize was adjudged in 1877, in the University of Cambridge* (Macmillan and Company).

Roxburgh, S. H., and J. B. Wilson (2000), Oikos **88** (2), 395.

Roy, F., M. Barbier, G. Biroli, and G. Bunin (2020), PLoS computational biology **16** (5), e1007827.

Saavedra, S., R. P. Rohr, J. Bascompte, O. Godoy, N. J. Kraft, and J. M. Levine (2017), Ecological Monographs **87** (3), 470.

Schmitz, D. A., T. Wechsler, I. Mignot, and R. Kümmerli (2024), ISME communications **4** (1), ycae045.

Serván, C. A., J. A. Capitán, J. Grilli, K. E. Morrison, and S. Allesina (2018), Nature ecology & evolution **2** (8), 1237.

Sinervo, B., and C. M. Lively (1996), Nature **380** (6571), 240.

Solé, R., and J. Bascompte (2012), *Self Organization in Complex Ecosystems* (Princeton University Press).

Solé, R., V. Maull, D. R. Amor, J. P. Mauri, and C.-P. Núria (2024), ACS Synthetic Biology.

Soliveres, S., F. T. Maestre, W. Ulrich, P. Manning, S. Boch, M. A. Bowker, D. Prati, M. Delgado-Baquerizo, J. L. Quero, I. Schöning, *et al.* (2015), Ecology letters **18** (8), 790.

Strauss, S. Y. (1991), Trends in Ecology & Evolution **6** (7), 206.

Strogatz, S. H. (2018), *Nonlinear dynamics and chaos with student solutions manual: With applications to physics, biology, chemistry, and engineering* (CRC press).

Strydom, T., G. V. Dalla Riva, and T. Poisot (2021), Frontiers in Ecology and Evolution **9**, 623141.

Suweis, S., F. Simini, J. R. Banavar, and A. Maritan (2013), Nature **500** (7463), 449.

Szathmáry, E., and I. Gladkih (1989), Journal of Theoretical Biology **138** (1), 55.

Szathmáry, E., and J. M. Smith (1997), Journal of theoretical biology **187** (4), 555.

Tilman, D., C. L. Lehman, and C. E. Bristow (1998), The American Naturalist **151** (3), 277.

Toni, B. (2014), in *New Frontiers of Multidisciplinary Research in STEAM-H (Science, Technology, Engineering, Agriculture, Mathematics, and Health)* (Springer) pp. 205–240.

Trefethen, L. N., and D. Bau (2022), *Numerical linear algebra* (SIAM).

Valverde, S., J. Piñero, B. Corominas-Murtra, J. Montoya, L. Joppa, and R. Solé (2018), Nature ecology & evolution **2** (1), 94.

Van Der Hofstad, R. (2024), *Random graphs and complex networks*, Vol. 2 (Cambridge university press).

Vandermeer, J. (1980), The American Naturalist **116** (3), 441.

Venturelli, O. S., A. V. Carr, G. Fisher, R. H. Hsu, R. Lau, B. P. Bowen, S. Hromada, T. Northen, and A. P. Arkin (2018), Molecular systems biology **14** (6), e8157.

Vincent, T. L., D. Scheel, J. S. Brown, and T. L. Vincent (1996), The American Naturalist **148** (6), 1038.

Wootton, J. T. (1994), Annual review of ecology and systematics , 443.

Wright, E. S., and K. H. Vetsigian (2016), Nature communications **7** (1), 11274.

Yin, H., and V. H. Rudolf (2024), Ecology Letters **27** (7), e14481.

Yodzis, P. (1988), Ecology **69** (2), 508.

Yu, X., M. F. Polz, and E. J. Alm (2019), The ISME journal **13** (6), 1602.

Zelnik, Y. R., N. Galiana, M. Barbier, M. Loreau, E. Galbraith, and J.-F. Arnoldi (2024), Ecology Letters **27** (1), e14358.
